# A Speedy Route to Multiple Highly Potent SARS-CoV-2 Main Protease Inhibitors

**DOI:** 10.1101/2020.07.28.223784

**Authors:** Kai S. Yang, Xinyu R. Ma, Yuying Ma, Yugendar R. Alugubelli, Danielle A. Scott, Erol C. Vatansever, Aleksandra K. Drelich, Banumathi Sankaran, Zhi Z. Geng, Lauren R. Blankenship, Hannah E. Ward, Yan J. Sheng, Jason C. Hsu, Kaci C. Kratch, Baoyu Zhao, Hamed S. Hayatshahi, Jin Liu, Pingwei Li, Carol A. Fierke, Chien-Te K. Tseng, Shiqing Xu, Wenshe Ray Liu

**Affiliations:** The Texas A&M Drug Discovery Laboratory, Department of Chemistry, Texas A&M University, College Station, TX 77843, USA; Department of Microbiology and Immunology, University of Texas Medical Branch, Galveston, TX 77555, USA; Molecular Biophysics and Integrated Bioimaging, Berkeley Center for Structural Biology, Lawrence Berkeley National Laboratory, Berkeley, CA 94720, USA; Department of Biochemistry and Biophysics, Texas A&M University, College Station, TX 77843, USA; Department of Pharmaceutical Sciences, UNT Health Science Center, Fort Worth, TX 76107, USA; Department of Molecular and Cellular Medicine, College of Medicine, Texas A&M University, College Station, TX 77843, USA

**Author notes:** Contribute equally to the paper (Kai S. Yang, Xinyu R. Ma, Yuying Ma, Yugendar R. Alugubelli, Danielle A. Scott, Erol C. Vatansever, Aleksandra K. Drelich). Correspondence should be addressed to Carol A. Fierke, Chien-Te K. Tseng, Shiqing Xu, and Wenshe Ray Liu.

**Keywords:** COVID-19, SARS-CoV-2, main protease, 3C-like protease, reversible covalent inhibitors

## Abstract

The COVID-19 pathogen, SARS-CoV-2, requires its main protease (SC2M^Pro^) to digest two of its translated polypeptides to form a number of mature proteins that are essential for viral replication and pathogenesis. Inhibition of this vital proteolytic process is effective in preventing the virus from replication in infected cells and therefore provides a potential COVID-19 treatment option. Guided by previous medicinal chemistry studies about SARS-CoV-1 main protease (SC1M^Pro^), we have designed and synthesized a series of SC2M^Pro^ inhibitors that contain β-(*S*-2-oxopyrrolidin-3-yl)-alaninal (Opal) for the formation of a reversible covalent bond with the SC2M^Pro^ active site cysteine C145. All inhibitors display high potency with IC_50_ values at or below 100 nM. The most potent compound MPI3 has as an IC_50_ value as 8.5 nM. Crystallographic analyses of SC2M^Pro^ bound to 7 inhibitors indicated both formation of a covalent bond with C145 and structural rearrangement from the apoenzyme to accommodate the inhibitors. Virus inhibition assays revealed that several inhibitors have high potency in inhibiting the SARS-CoV-2-induced cytopathogenic effect in both Vero E6 and A549 cells. Two inhibitors MP5 and MPI8 completely prevented the SARS-CoV-2-induced cytopathogenic effect in Vero E6 cells at 2.5-5 μM and A549 cells at 0.16-0.31 μM. Their virus inhibition potency is much higher than some existing molecules that are under preclinical and clinical investigations for the treatment of COVID-19. Our study indicates that there is a large chemical space that needs to be explored for the development of SC2M^Pro^ inhibitors with extreme potency. Due to the urgent matter of the COVID-19 pandemic, MPI5 and MPI8 may be quickly advanced to preclinical and clinical tests for COVID-19.

## INTRODUCTION

Coronaviruses (CoVs) are a group of related RNA viruses that cause diseases in a wide range of vertebrates including humans and domestic animals.^1^ Before 2003, there were only two CoVs, HCoV-229E and HCoV-OC43, known as human pathogens.^2, 3^ The SARS pandemic in 2003 led to the revelation of SARS-CoV-1, a pathogen causing a severe respiratory infection.^4^ The subsequent surge in CoV research resulted in the discovery of two additional human CoVs, HCoV-NL63 and HCoV-HKU1, that are mildly pathogenic.^5, 6^ One addition to this group was MERS-CoV that emerged in 2012 as a pathogen causing a severe respiratory infection.^7^ Although SARS-CoV-1 and MERS-CoV are highly lethal pathogens, the public health, social, and economic damages that they have caused are diminutive in comparison to that from SARS-CoV-2, a newly emerged human CoV pathogen that causes COVID-19.^8^ Rival only to the 1918 influenza pandemic, the COVID-19 pandemic has led to catastrophic impacts worldwide. As of July 13^th^, 2020, the total global COVID-19 cases have surpassed 12 million with more than 570,000 deaths.^9^ To alleviate catastrophic damages of COVID-19 on public health, society and economy, finding timely treatment options is of paramount importance.

Similar to all other CoVs, SARS-CoV-2 is an enveloped, positive-sensed RNA virus with a genome of nearly 30 kb in size.^10^ Its genome encodes 10 open reading frames (ORFs). The largest ORF, ORF1ab encompasses more than two thirds of the whole genome. Its translated products, ORF1a (∼500 kDa) and ORF1ab (∼800 kDa),^11^ are very large polypeptides that undergo proteolytic cleavage to form 15 mature proteins. These are nonstructural proteins (Nsps) that are essential for the virus to modulate human cell hosts for efficient viral protein expression, genome replication, virion packaging, viral genome transcription and replication, and viral genomic RNA processing to evade the host immune system. The proteolytic cleavage of ORF1a and ORF1ab is an autocatalytic process. Two internal polypeptide regions, Nsp3 and Nsp5, possess cysteine protease activity that cleaves themselves, and all other Nsps, from the two polypeptides. Nsp3 is commonly referred to as papain-like protease (PL^Pro^), and Nsp5 as 3C-like protease (3CL^Pro^) or, more recently, main protease (M^Pro^).^12^ Although we have yet to understand SARS-CoV-2 biology and COVID-19 pathogenesis, previous studies of SARS-CoV-1 have established that activity of both PL^Pro^ and M^Pro^ is essential to viral replication and pathogenesis. Of the two proteases, M^Pro^ processes 12 out of the total 15 Nsps; inhibition of this enzyme is anticipated to have more significant impacts on the viral biology than that of PL^Pro^. Therefore, small molecule medicines that potently inhibit SARS-CoV-2 M^Pro^ (SC2M^Pro^) are potentially effective treatment options for COVID-19.^13, 14, 15^ In this work we report our progress in the development of potent SC2M^Pro^ inhibitors.

## RESULTS

### The design of β-(*S-*2-oxopyrrolidin-3-yl)-alaninal (Opal)-based, reversible covalent inhibitors for SC2M^Pro^

Although we are at the inaugural stage of learning medicinal chemistry to inhibit SC2M^Pro^, much has been learned from studies of SARS-CoV-1 M^Pro^ (SC1M^Pro^) that shares 96% sequence identity with SC2M^Pro^.^16^ SC1M^Pro^ has a large active site that consists of several smaller pockets for the recognition of residues at P1, P2, P4, and P3’ positions in a protein substrate (Figure 1A).^17^ P4 is typically a small hydrophobic residue while P2 and P3’ are large. For all Nsps that are processed by SC1M^Pro^ and SC2M^Pro^, Gln is the P1 residue at their cleavage sites. In order to bind the P1 Gln, SC1M^Pro^ forms strong van der Waals interactions with the Gln side chain, and also utilizes two hydrogen bonds with the Gln side chain amide oxygen and *α*-carbonyl oxygen atoms (Figure 1B). Previous efforts in the development of irreversible covalent inhibitors for SC1M^Pro^ primarily focused on fixing the P1 residue as a more potent β-*S*-2-oxopyrrolidine-containing Gln analog, and changing the scissile backbone amide to an alkene Michael acceptor in order to react with the active site cysteine C145, as well as varying substituents on two sides to improve potency.^18^ The enhanced potency from the use of the β-*S*-2-oxopyrrolidine-containing Gln analog is most probably due to the reduction of entropy loss during the binding of SC1M^pro^ to the more rigid lactam compared to the flexible Gln. Although converting the scissile backbone amide to a Michael acceptor in a SC1M^Pro^ ligand turns it into a covalent inhibitor, it eliminates the critical hydrogen bond between the P1 *α*-carbonyl oxygen and SC1M^Pro^. Therefore, most Michael acceptor inhibitors developed for SC1M^Pro^ and recently for SC2M^Pro^ tend to have efficacy with low micromolar or submicromolar IC_50_ values rather than low nanomolar levels.^13, 18^ To maintain this critical hydrogen bond and exploit a covalent interaction with the active site cysteine C145 to form a hemiacetal for high affinity, both aldehyde and ketoamide moieties have been used to replace the P1 *C*-side *α*-amide to develop potent reversible covalent inhibitors for SC1M^Pro^. For aldehyde-based inhibitors, a typical potent inhibitor contains Opal at the P1 site that consists of a β-*S*-2-oxopyrrolidine side chain and an *α*-aldehyde for both taking advantage of strong interactions with the SC1M^Pro^ P1-binding pocket and the formation of a reversible covalent bond with C145 (Figure 1C). Typical examples of this design include GC376 that was originally developed for M^Pro^ from feline infectious peritonitis (FIP) CoV and two inhibitors, 11a and 11b, that were recently developed for SC2M^Pro^.^19, 20^ Given its relative simplicity, we have followed a similar scheme according to structure diagrams shown in Figure 1D to design and synthesize reversible covalent inhibitors for SC2M^Pro^ and pursued structural variations at P2, P3, and R positions for improved potency.

**Figure 1:**
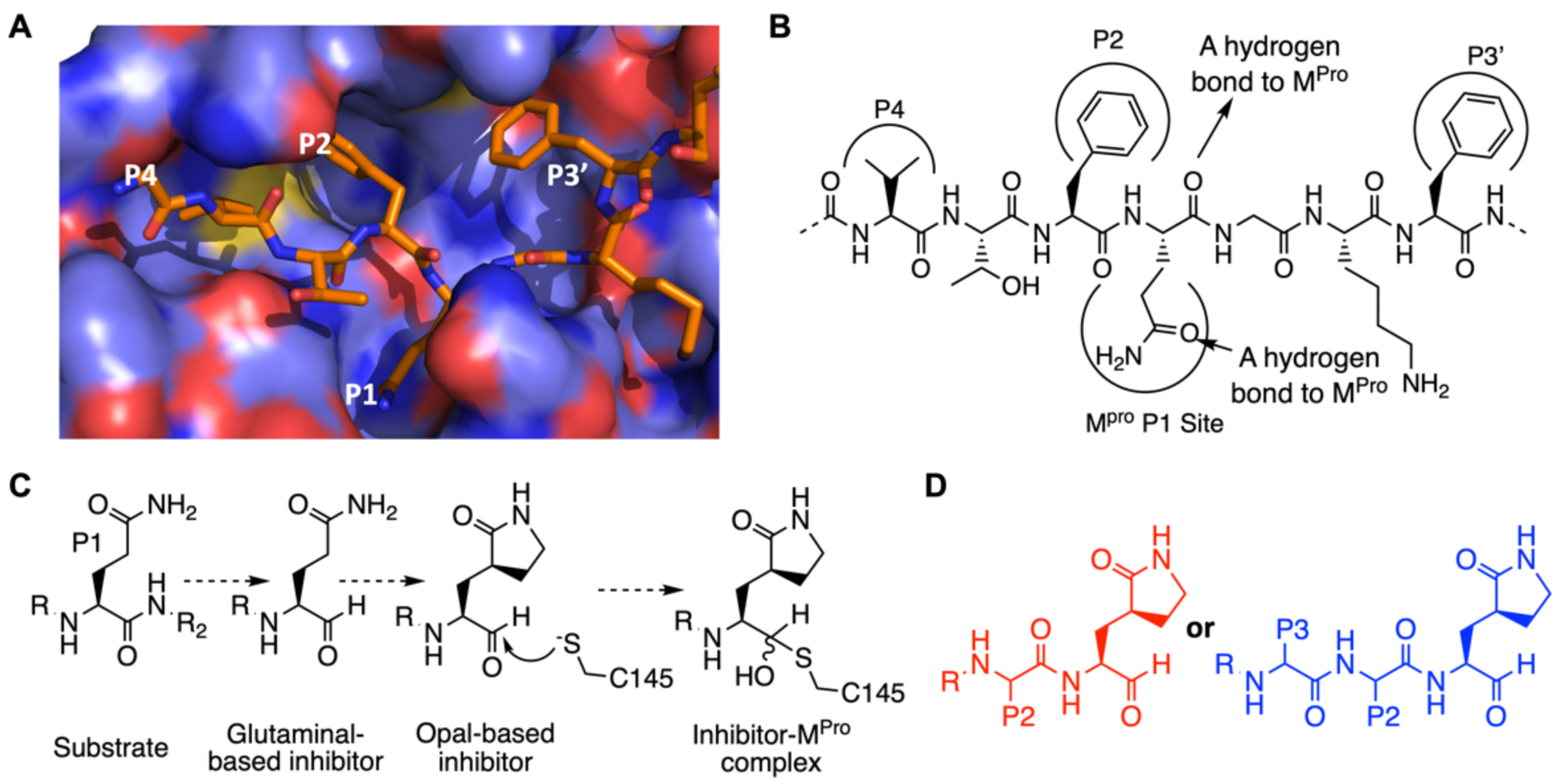
The design of SC2M^Pro^ inhibitors based on medicinal chemistry learned from SC1M^Pro^ studies. (A) The structure of SC1M^Pro^ complexed with a peptide substrate (based on the pdb entry 5b6o). Active site cavities that bind P1, P2, P4, and P3’ residues in the substrate are labeled. (B) A schematic diagram that shows interactions between SC1M^Pro^ and a substrate. (C) A scheme in which a substrate P1 residue is converted to glutaminal and then β-(*S*-2-oxopyrrolidin-3-yl)-alaninal (Opal) to form a reversible covalent inhibitor that reacts with the SC1M^Pro^ active site cysteine C145. (D) Scaffold structures of Opal-based inhibitors designed for SC2M^Pro^.

### Synthesis and IC50 characterization of SC2M^Pro^ inhibitors (MPIs)

GC376 (Figure 2A) has confirmed potency against SC1M^Pro^.^21^ We purchased it as a potential SC2M^Pro^ inhibitor. We designed two similar dipeptidyl compounds MPI1-2 (Figure 2A) and synthesized them according to a synthetic scheme shown in Supplementary Scheme 1. Both MPI1 and MPI2 have Phe at the P2 site which was previously shown to contribute to strong bonding to SC1M^Pro^.^18^ MPI2 has also an *o-*fluoro-*p-*chlorocinnamyl group as an *N*-terminal cap. This group is more rigid than the CBZ group and therefore possibly introduces a strong interaction with the P4-binding pocket in SC2M^Pro^.^22^ To characterize IC_50_ values of all three molecules for inhibition of SC2M^Pro^, we recombinantly expressed a 6×His-SUMO-SC2M^Pro^ fusion protein in *E. coli* and purified and digested this protein with SUMO protease to obtain intact SC2M^Pro^ with more than 95% purity. We used a previously described fluorescent peptide assay (see supplementary data) to measure the IC_50_ values for GC376, MPI1, and MPI2 as 31 ± 4, 100 ± 23, and 103 ± 14 nM, respectively (Figure 2B).^15^ Our determined IC_50_ value for GC376 agrees well with that from Ma *et al*.^23^ In the light of the publication of inhibitors 11a and 11b that showed similar IC_50_ values as 53 ± 5 and 40 ± 2 nM, respectively,^24^ we shifted our focus from the synthesis of bipeptidyl inhibitors to that of tripeptidyl inhibitors. By adding one more residue to the design of inhibitors, additional interactions with SC2M^Pro^ might be achieved to improve potency. In the design of SC1M^Pro^ inhibitors, Leu, Phe, and Cha (cyclohexylalanine) are three residues used frequently at the P2 site and Val and Thr(tBu) (O-tert-butyl-threonine) are two residues used frequently at the P3 site.^18^ Installation of these residues at two sites and including CBZ as a N-terminal cap led to the design of six compounds MPI3-8 (Figure 2A). We added one additional compound MPI9 that has an *o-*fluoro-*p-*chlorocinnamyl cap to this series to compare the effect of the two N-terminal caps on the inhibitor potency for SC2M^Pro^. We synthesized all 7 compounds according to a synthetic scheme presented in Supplementary Scheme 2 and characterized their IC_50_ values using the fluorescent peptide assay. As shown in Figure 2B, all inhibitors have IC_50_ values below 100 nM, except for MPI8 that has an IC_50_ value as 105 ± 22 nM. The most potent compound is MPI3 with an IC_50_ value as 8.5 ± 1.5 nM, followed by MPI4 and MPI5 with IC_50_ values as 15 ± 5 and 33 ± 2 nM, respectively. We also synthesized 11a (named as MPI10 in our series) according to the procedure in Dai *et al*. and used it as a positive control in our enzyme and viral inhibition analyses.^24^ Using our fluorescent peptide assay, we determined the IC_50_ value of 11a as 31 ± 3 nM. As far as we know, MPI3 is the most potent SC2M^Pro^ inhibitor that has been reported so far. From the perspective of enzyme inhibition, Leu and Val are optimal residues at P2 and P3 sites in an inhibitor for improved affinity for SC2M^Pro^ and CBZ also enhances affinity compared to the *o-* fluoro-*p-*chlorocinnamyl as a N-terminal capping group.

**Figure 2:**
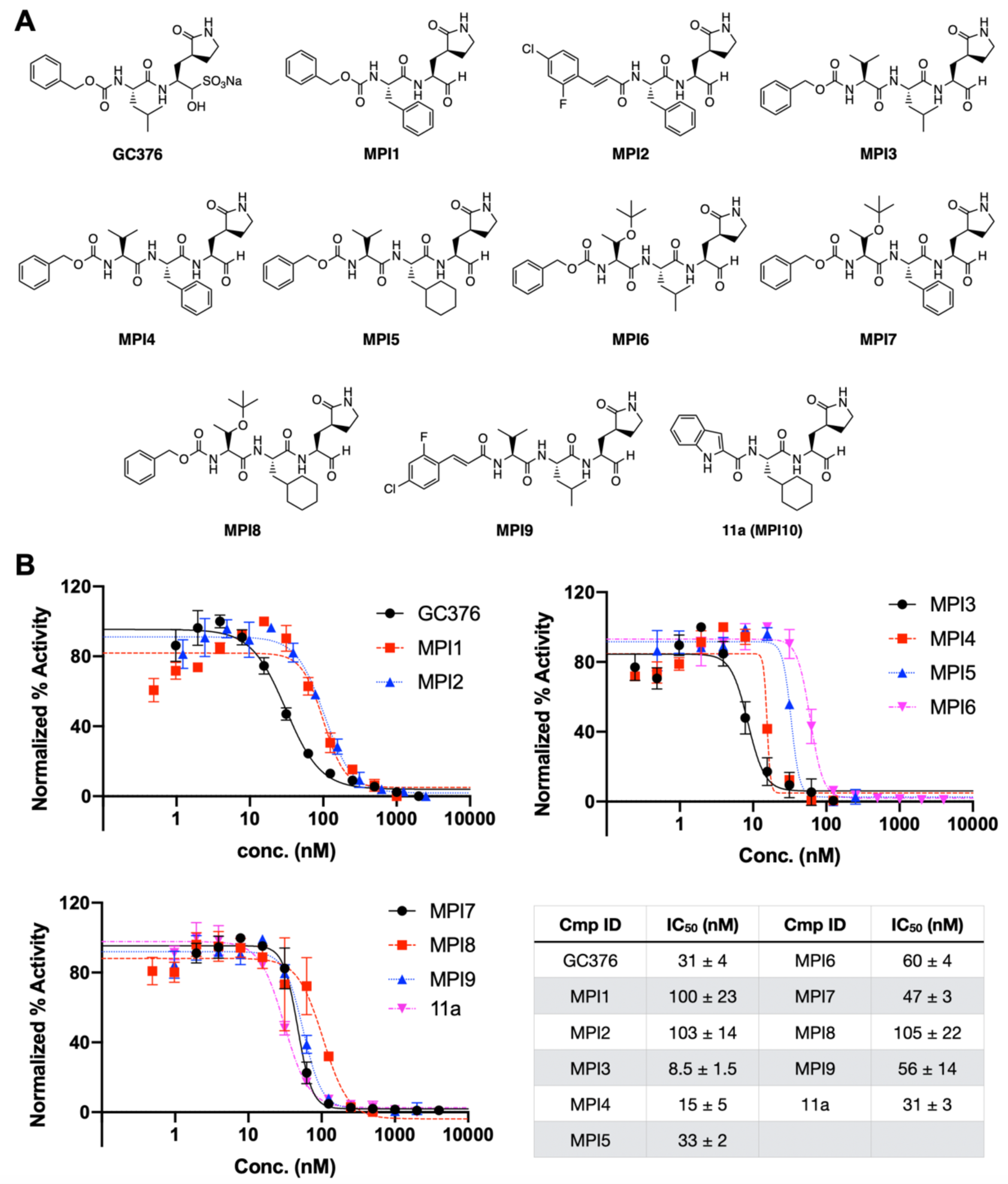
SC2M^Pro^ inhibitors and their determined IC_50_ values. (A) Structures of GC376 and 10 Opal-based inhibitors. (B) The inhibition curves of all 11 inhibitors toward SC2M^Pro^. The determined IC_50_ values are presented in the associated table.

### Structural characterization of SC2M^Pro^ interactions with Opal-based inhibitors

In order to understand how our designed inhibitors interact with SC2M^Pro^ at its active site, we screened crystallization conditions for apo-SC2M^Pro^, soaked apo-SC2M^Pro^ crystals with different inhibitors, and determined the crystal structures of these inhibitors in complex with SC2M^Pro^. We used Hampton Research Crystal Screen and Index kits to perform initial screening and identified several conditions that yielded single crystals of apo-SC2M^Pro^. For all conditions, crystals were in a thin plate shape (Supplementary Figure S1). The best crystallization condition contained 0.2 M dibasic ammonium phosphate and 17% PEG 3,350. We refined the structure of apo-SC2M^Pro^ against diffraction data to 2.0 Å resolution. In the apoenzyme crystals, SC2M^Pro^ existed as a monomer in the crystallographic asymmetric unit and packed relatively densely. The active site of each monomer stacked upon another monomer (two representative monomers are shown in red and blue respectively in Figure 3A). This close contact and dense protein packing made the diffusion of inhibitors to the active site quite slow. We soaked apo SC2M^Pro^ crystals with all 9 inhibitors that we synthesized and collected and processed their X-ray diffraction data for structural determination. For crystals that we soaked with the inhibitors for just 2 h, we did not find observable ligand electron density at the enzyme active site. For 7 inhibitors including MPI1 and MPI3-8, we performed two-day soaking and observed clear electron density in the difference maps in the active site of the enzyme. For MPI2 and MPI9, we were not able to determine structures of their complexes with SC2M^Pro^ due to cracking of the crystals upon soaking with the inhibitors. For MPI3, the electron density around the P1, P2, and P3 residues were well defined, and the covalent interaction between the C145 side chain thiolate and the Opal aldehyde to form a hemiacetal was clearly observable (Figure 3B). The electron density around CBZ was very weak indicating flexible CBZ binding around the enzyme P4-binding pocket. Figure 3C shows the superposition of apo SC2M^Pro^ and the SC2M^Pro^-MPI3 complex structures. The two structures display very little overall variation with RMSD as 0.2 Å. Around the active site in the two structures, large structural rearrangements exist for residues M49 and N142 and the loop region that contains P168. In apoenzyme, the side chain of M49 folds into the P2-binding pocket. It flips toward the solvent to make space available for the binding of the P2 Leu in MPI3. The side chain of N142 rotates by almost 180° between the two structures and adopts a conformation in the SC2M^Pro^-MPI3 complex that closely caps the P1-binding site for strong van der Waals interactions with the Opal residue in MPI3. In the SC2M^Pro^-MPI3 complex, the P168-containing loop is pushed away from its original position in the apoenzyme, probably by interaction with the CBZ group, which triggers a position shift for the whole loop. Except for M49, N142, and the P168-containing loop, structural orientations of all other residues at the active site closely resemble each other in the two structures. In the active site, MPI3 occupies the P1, P2, and P4-binding pockets and leaves the large P3’-binding pocket empty (Figure 3D). Extensive hydrogen bonding and van der Waals interactions in addition to the covalent interaction with C145 contribute to the strong binding of MPI3 to SC2M^Pro^ (Figure 3E). Residues F140, N142, H163, E166, and H172 form a small cage to accommodate the Opal side chain. Three hydrogen bonds form between the Opal lactam amide and the E166 side chain carboxylate, H163 imidazole, and F140 backbone carbonyl oxygen. The precise fitting of Opal into the P1-binding pocket and the formation of three hydrogen bonds explain the preferential binding of the Opal side chain to this pocket. In the SC2M^Pro^-MPI3 complex, M49 flips from the P2-binding pocket to leave space for the binding of the P2 Leu in MPI3. Residues H41, M49, M165 and D187, backbones of the M165-containing strand, and the D-187-containing loop form a hydrophobic pocket that is in a close range of van der Waals interactions with the P2 Leu in MPI3. We observe Leu as the best residue in this position probably due to this close van der Waals interaction range for the recognition of the P2 Leu side chain. The enzyme has no P3-binding pocket. However, the P3 Val in MPI3 positions its side chain in van der Waals interaction distance to E166 and P168. In the structure, CBZ narrowly fits into the P4-binding pocket and the channel formed between the P168-and Q192-containing loops. The P168 loop rearranges its position from that in apoenzyme to accommodate the CBZ group. The CBZ group also has weak electron density. These observations indicate that CBZ is not an optimal structural moiety for interaction at these sites. Besides interactions involving side chains and the CBZ group in MPI3, its two backbone amides and carbamate form 6 hydrogen bonds with the enzyme. Two of them are formed between the P3 Val in MPI3 and the backbone amino and carbonyl groups of E166 in SC2M^Pro^. One water molecule mediates a hydrogen bond bridge between the P2 Leu amino group in MPI3 and the Q189 side chain amide in SC2M^Pro^. For the P1 Opal residue in MPI3, its *α*-amino group forms a hydrogen bond with the H164 *α*-carbonyl oxygen in the enzyme. The original aldehyde oxygen in MPI3 forms two hydrogen bonding interactions, one with the *α*-amino group of G143 and the other the C145 *α*-amine in SC2M^Pro^. The two hydrogen bonds are probably the reason that Opal-based reversible covalent inhibitors are typically stronger than Michael acceptor inhibitors, in which the original scissile amide is replaced with an alkene, for inhibition of M^Pro^ enzymes. In the structures of SC2M^Pro^ complexes with the other 6 inhibitors, we observed similar structure rearrangements at M49, N142, and the P168-containing loop to accommodate inhibitors and a covalent interaction with C145 (Supplementary Figure S2 and Supplementary Table S1). The structural superposition of all seven inhibitor complexes with SC2M^Pro^ show a similar binding mode for all inhibitors to the SC2M^Pro^ active site (Figure 4F). The electron density around the CBZ group for all inhibitors is not well defined. Some inhibitor-enzyme complex structures have relatively weak electron density at P2 and P3 residues in their inhibitors possibly due to low occupancy or flexible conformations at these positions.

**Figure 3:**
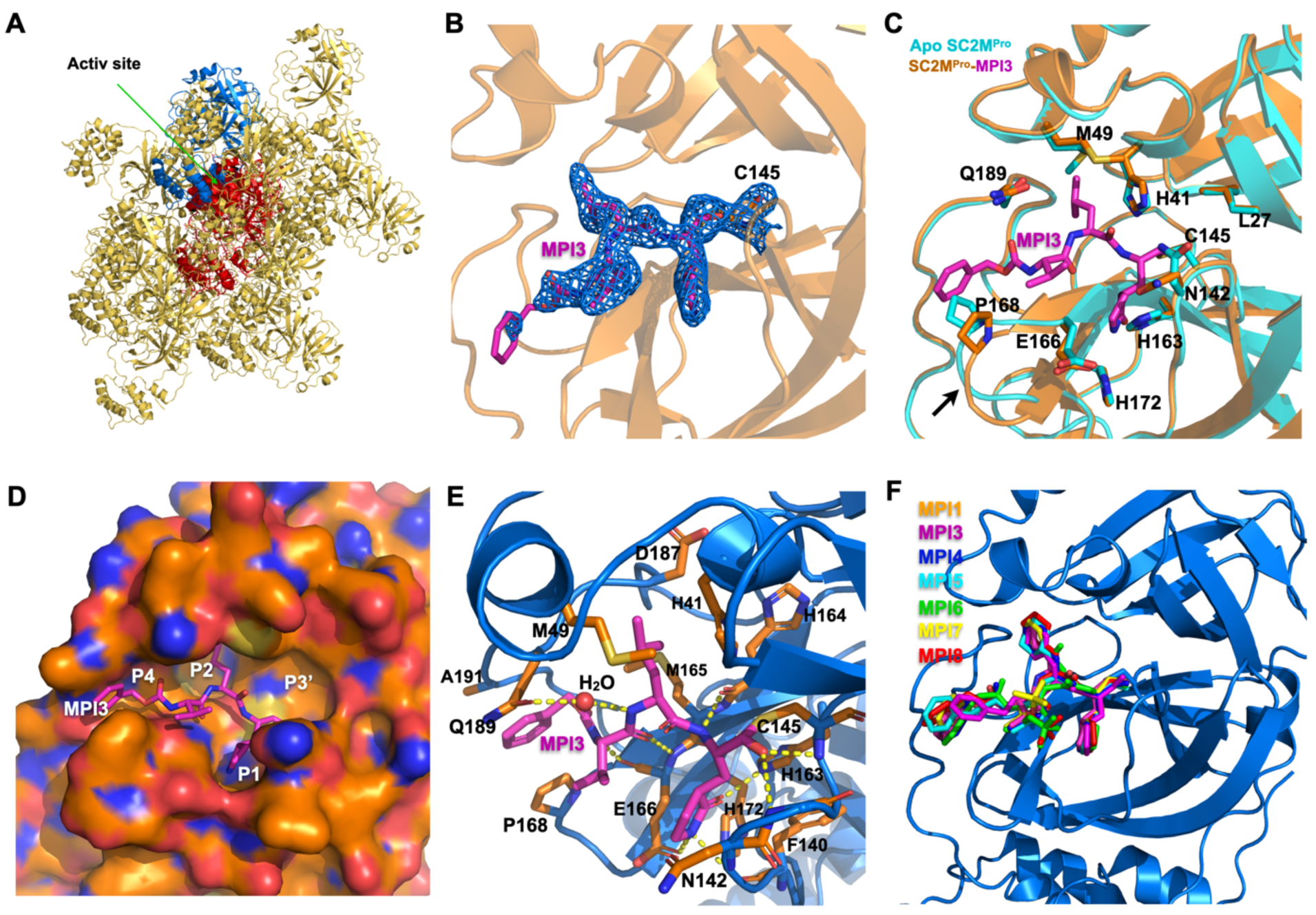
X ray crystallography analysis of SC2M^Pro^ in its apo-form and complexes with different inhibitors. (A) The packing of apo-SC2M^Pro^ in its crystals. An asymmetric unit monomer is colored in red in the center. Its active site is presented as a concaved surface. Another monomer that stacks upon the active site of the red monomer is colored in blue. (B) A contoured 2Fo-Fc map at the 1 *σ* level around MPI3 and C145 in the active site of SC2M^Pro^. A covalent bond between MPI3 and C145 is observable. (C) The structure overlay between apo-SC2M^Pro^ and the SC2M^Pro^-MPI3 complex. A black arrow points to a region that undergoes structure rearrangement in the SC2M^Pro^-MPI3 complex from apoenzyme to accommodate MPI3. (D) The occupation of the active site cavity of SC2M^Pro^ by MPI3. The enzyme is shown in its surface presentation mode. (E) Extensive hydrogen bonding and van del Waals interactions between SC2M^Pro^ and MPI3. The backbone of SC2M^Pro^ is colored in marine blue and side chain carbon atoms in orange. Hydrogen bonds between MPI3 and SC2M^Pro^ are depicted as yellow dashed lines. (F) The overlay of 7 Opal-based inhibitors at the active site of SC2M^Pro^. Inhibitors are colored according to their color-coded names shown in the Figure. All images were made using the program PyMOL.

**Figure 4:**
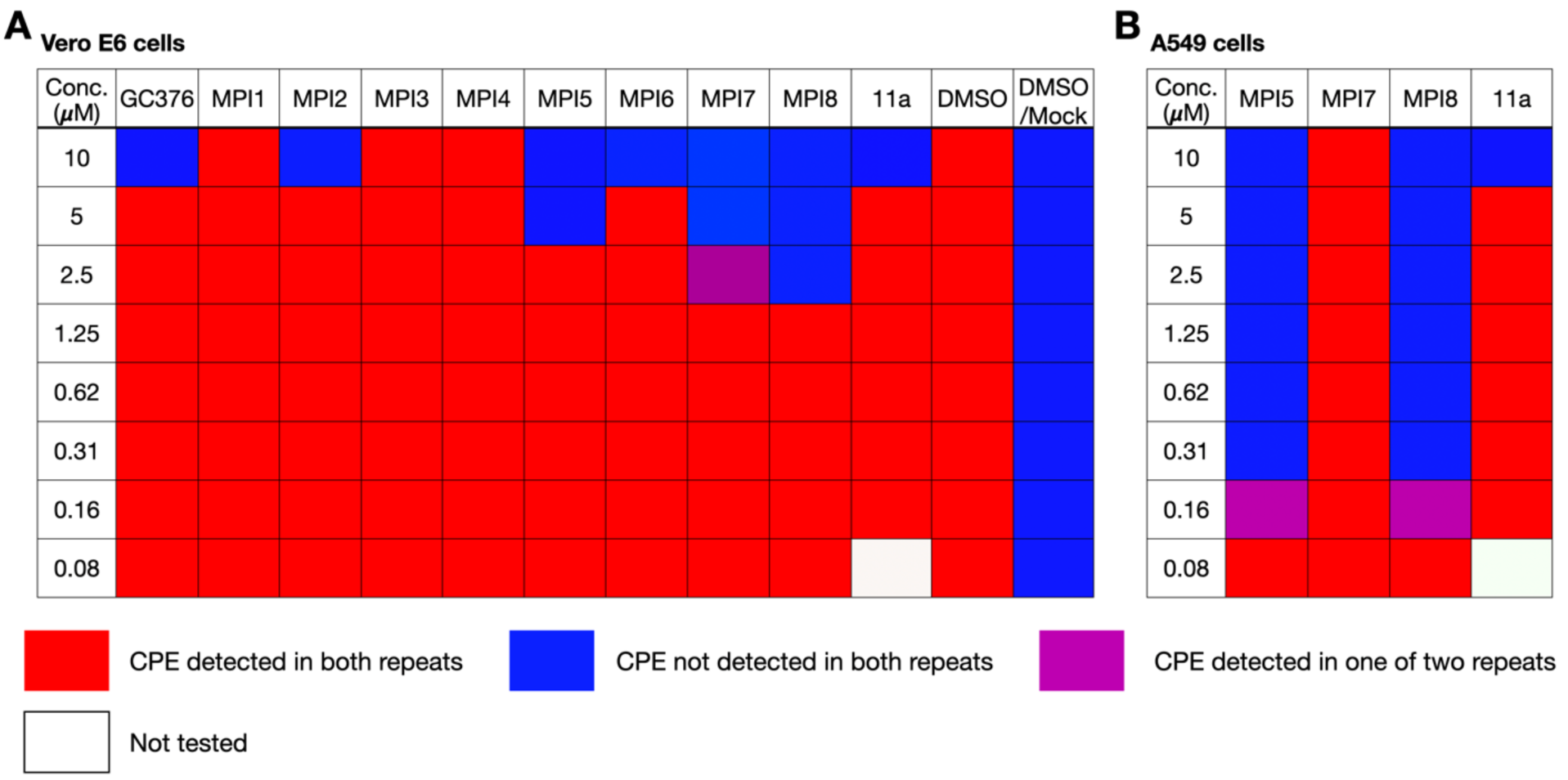
The SARS-CoV-2 viral inhibition results of selected inhibitors in (A) Vero E6 and (B) A549 cells. CPE is an abbreviation for cytopathogenic effect.

### SARS-CoV-2 inhibition analysis of GC376, MPI1-8, and 11a

To evaluate our molecules’ ability to inhibit SARS-CoV-2, we conducted a live virus-based microneutralization assay in Vero E6 cells. Vero E6 is a kidney epithelial cell line isolated from African Green Monkey. It has been used widely as a model system for human CoV studies.^25^ We tested 10 molecules including GC376, MPI1-8, and 11a in a concentration range from 80 nM to 10 μM and recorded cytopathogenic effect (CPE) observed in SARS-CoV-2-infected Vero E6 cells that we cultured in the presence of different concentrations of inhibitors. 11a was included as a positive control. For each condition, we conducted two repeats. Although it was disappointing that MPI3 was not able to completely prevent CPE at all tested concentrations, several inhibitors abolished CPE: GC376, MPI2, MPI6, and 11a at 10 μM, MPI5 at 5 μM, MPI7 at 2.5-5 μM, and MPI8 at 2.5 μM (Figure 4A). Three compounds MPI5, MPI7 and MPI8 performed better than GC376 that has been recently explored by Anivive Lifesciences for the treatment of COVID-19 and 11a that has been considered for COVID-19 clinical studies.^15^ Since we only recorded complete abolition of CPE, the real EC_50_ values for these compounds are expected to be much lower than lowest observed concentrations for CPE abolishment. Encouraged by our results in Vero E6 cells, we tested the three most potent compounds MPI5, MPI7, and MPI8 and also 11a in A549 cells. The A549 cell line was derived from human alveolar epithelial cells. It mimics the SARS-CoV-2 infection of the human respiratory tract system better than Vero E6.^26^ We tested a same concentration range for all four compounds. MPI7 was not able to completely abolish CPE at all tested conditions. However, both MPI5 and MPI8 performed much better than in Vero E6 cells with complete abolition of CPE at 160-310 nM and much better than 11a (Figure 4B). 11 displayed potency similar to that shown in Vero E6 cells. Given that real EC_50_ values are expected to be lower than the lowest observed concentration for CPE abolishment, MPI5 and MPI8 are, as far as we know, the most potent anti-SARS-CoV-2 small molecules in infected cells that have been reported so far.

## DISCUSSION

Guided by previous medicinal chemistry studies about SC1M^Pro^, we designed and synthesized a number of Opal-based dipeptidyl and tripeptidyl inhibitors that potently inhibit SC2M^Pro^, an essential enzyme for SARS-CoV-2, the pathogen of COVID-19. As the most potent inhibitor of SC2M^Pro^, MPI3 displayed an IC_50_ value of 8.5 nM. As far as we know, this is the lowest reported IC_50_ for known SC2M^Pro^ inhibitors. During the search of optimal conditions for IC_50_ characterizations, we noticed that 10 nM was the lowest SC2M^Pro^ concentration that could provide reliable activity.^15^ Since the determined IC_50_ value for MPI3 is lower than the used enzyme concentration, the real K_d_ value of MPI3 is expected to be much lower than 8.5 nM. X-ray crystallography analysis of the SC2M^Pro^-MPI3 complex revealed that MPI3 fits precisely in the P1-and P2-binding pockets at the SC2M^Pro^ active site. Strong van der Waals interactions at the P1-and P2-binding pockets, 9 hydrogen bonds with active site residues, and the covalent interaction with C145 necessitate high affinity of MPI3 to SC2M^Pro^. The structures of SC2M^Pro^ complexes with MPI3 and other inhibitors showed a relatively loosely bound N-terminal capping group. We expect that optimization at this site will contribute to the generation of more potent inhibitors in the future. Although MPI3 is the most potent inhibitor for the enzyme, its cellular activity in inhibiting SARS-CoV-2 is much lower than several other inhibitors we have generated. A likely reason is its lower cellular stability. MPI3 has Leu and Val at its P2 and P3 sites respectively. Both are native amino acids that are expected to be targeted by both extracellular and cellular proteases. Since Leu and Val are optimal residues at two sites, modest changes based on these structures will be necessary for both maintaining high potency in inhibiting SC2M^Pro^ and improving cellular stability for enhanced cellular activity in inhibiting the virus. As such, Val and Leu analogs at these two sites need to be explored. Since both MPI5 and MPI8 show high anti-SARS-CoV-2 activity in both Vero E6 and A549 cells and each has Cha at their P2 site, we suggest maintaining Opal and Cha at P1 and P2 sites and varying the residue at P3 and the N-terminal capping moiety to improve anti-SARS-CoV-2 activity in cells. Based on our structures of SC2M^Pro^ complexes with 7 inhibitors, the P1 Opal occupies precisely the P1-binding pocket in SC2M^Pro^ and three hydrogen bonds to the Opal lactam amide are critical in maintaining strong binding to SC2M^Pro^. Chemical space to manipulate the P1 residue in an inhibitor for improved binding to SC2M^Pro^ is minimal. But one direction that may be explored is to introduce additional heteroatom(s) to Opal for the formation of hydrogen bond(s) with the N142 side chain amide. In the SC2M^Pro^-MPI3 complex, the N142 side chain flips by about 180° from its position in apoenzyme to form a closed P1-binding pocket. However, only van der Waals interactions with Opal are involved with N142. Given the close distance between the Opal side chain and the side chain amide of N142, some hydrogen bonds may be designed for improved potency. In all our designed inhibitors, an Opal aldehyde is involved in the formation of a covalent interaction with C145. This design, although necessary for the formation of a hemiacetal covalent complex, effectively excludes the exploration of the P3’-binding site in SC2M^Pro^ for improved potency in a designed inhibitor. Figure 4D illustrates that the P3’-binding pocket is completely empty. In our early discussion, we argued that it is critical to maintain the hydrogen bond between the scissile amide oxygen in a substrate and SC2M^Pro^ for high affinity. Changing the scissile amide to an aldehyde in an inhibitor is effective in maintaining this hydrogen bond and allows a covalent interaction with C145. Two hydrogen bonds formed between the hemiacetal alcohol and SC2M^Pro^ contribute to high potency of this group of molecules. One possibility to keep all of these described potentials that are associated with the Opal aldehyde in an inhibitor but to reach out to the P3’-binding site for even stronger binding to SC2M^Pro^ is to convert the aldehyde to an *α*-ketoamide. Lower reactivity of *α*-ketoamide in comparison to aldehyde toward C145 to generate a hemiacetal might be compensated and surpassed by adding chemical moieties to the *α*-ketoamide for strong interactions with the P3’-binding pocket. All these possibilities are worth exploring.

In our study, cell-based anti-SARS-CoV-2 activity of our designed inhibitors do not correlate with their IC_50_ values in inhibiting SC2M^Pro^. This is expected since cellular stability and other features of these inhibitors are very different. However, information regarding both enzyme inhibition IC_50_ values and anti-SARS-CoV-2 activity is critical for the design of a new generation of inhibitors that perform excellent in both aspects. Given that MPI3 has already reached a single digit nanomolar IC_50_ value and MPI5 and MPI8 display high potency in inhibiting SARS-CoV-2, merging features of the three molecules will lead to inhibitors with extreme potency in inhibiting the virus. Our antiviral assays indicated that MPI5 and MPI8 performed much better than GC376 and 11a, two molecules that have been explored for COVID-19 preclinical and clinical tests. Due to the urgent matter of COVID-19, we urge preclinical and clinical tests of both MPI5 and MPI8 in the treatment of COVID-19 and are actively exploring them.

## ACKNOWELDGEMENTS

This work was supported in part by National Institutes of Health (grants R01GM127575 and R01GM121584 to W. R. Liu and grant R01AI145287 to P. Li) and Welch Foundation (grant A-1715 to W.R. Liu and grant A-1987 to C. A. Fierke). We are grateful to Prof. Thomas Meek for allowing us to use the LC-MS system in his group for purification and characterization of some of our compounds. The ALS-ENABLE beamlines are supported in part by the National Institutes of Health, National Institute of General Medical Sciences, grant P30 GM124169-01 and the Howard Hughes Medical Institute. The Advanced Light Source is a Department of Energy Office of Science User Facility under Contract No. DE-AC02-05CH11231.

## AUTHOR CONTRIBUTIONS

W.R.L. conceived the project. W.R.L., S.X., C.K.T. C.A.F., J.L., and P.L. designed experiments. K.S.Y., X.R.M., Y.M., Y.R.A., D.A.S., E.C.V., A.K.D., Z.Z.G., L.R.B., H.E.W., Y.J.S., J.C.H., K.C.K., B.Z., and H.S.H. performed the experiments. S.X., C.K.T, C.A.F., and W.R.L. wrote the manuscript. All authors approved the final manuscript before submission.

## SUPPLEMENTARY MATERIALS

Please see the associated Supplementary Materials for details about compound synthesis, protein expression, inhibition characterization, X-ray crystallography analysis, virus microneutralization assays, Nuclear magnetic resonance and mass spectrometry characterization of synthesized compounds and related spectra, X-ray data collection and structure refinement data tables, and associated Supplementary Figures that show synthetic schemes and structures of SC2M^Pro^ complexes with different inhibitions.

