## Supplementary Materials for "A Speedy Route to Multiple Highly Potent SARS-CoV-2 Main Protease Inhibitors"

†Contribute equally to the paper

Keywords: COVID-19, SARS-CoV-2, main protease, 3C-like protease, reversible covalent inhibitors

### Materials & Methods

#### Recombinant SC2M$^{pro}$ protein expression and purification

The construct pET28a-His-SUMO-SC2M$^{pro}$ construct was made based on the a pET28a plasmid modified with N-terminal His-SUMO tag. The gene encoding SC2M$^{pro}$ was amplified from a previous plasmid pBAD-sfGFP-M$^{pro}$ using the forward primer 5′-CGCGGATCCGGGTTTCGCAAG-3′ and the reverse primer 5′-CCGCTCGAGTTACTGAAAAGTTACGCC-3′. The amplified PCR product was digested with BamHI and XhoI and ligated into the vector pET28a-His-SUMO plasmid digested with the same restriction enzymes. The gene sequence of His-SUMO-SC2M$^{pro}$ was verified by sequencing at Eton Bioscience Inc.

The pET28a-His-SUMO-SC2M$^{pro}$ construct was transformed into *E. coli* strain BL21(DE3). Cells were cultured at 37°C in 6 L 2xYT medium with kanamycin (50 µg/mL) for 3 h and induced with isopropylβ-D-1-thiogalactoside (IPTG) at final concentration of 1 mM when the $OD_{600}$ reached 0.8. After 3 h, cells were harvested by centrifugation at 12,000 rpm, 4°C for 30 min. Cell pellets were resuspended in 150 mL buffer A (20 mM Tris, 100 mM NaCl, 10 mM imidazole, pH 8.0) and then lysed by sonication on ice. The lysate was clarified by centrifugation at 16,000 rpm, 4°C for 30 min. The supernatant was loaded onto a nickel-chelating column with High Affinity Ni-Charged Resin (GenScript) and washed with 10 column volumes of buffer A to remove unspecific binding proteins, followed by elution using buffer B (20 mM Tris, 100 mM NaCl, 250 mM imidazole, pH 8.0). The protein eluates were subjected to buffer exchange with buffer C (20 mM Tris, 10 mM NaCl, 1mM dithiothreitol (DTT), pH 8.0) by using HiPrep 26/10 desalting column (GE Healthcare). The His-SUMO-SC2M$^{pro}$ proteins were digested with SUMO protease overnight at 4°C. The digested protein was applied to nickel-chelating column again to remove the His-tagged SUMO protease, the His-SUMO tag, and protein with uncleaved His-SUMO tag. The tag-free SC2M$^{pro}$ protein was loaded onto an anion-exchange column with Q Sepharose, Fast Flow (GE Healthcare) equilibrated with buffer C for further purification. The column was eluted by buffer D (20 mM Tris, 1 M NaCl, 1 mM DTT, pH 8.0) with a linear gradient ranging from 0 to 500 mM NaCl (10 column volumes buffer). Fractions eluted from the anion-exchange column were condensed and loaded to size exclusion column with HiPrep 16/60 Sephacryl S-100 HR (GE Healthcare) pre-equilibrated with buffer E (20 mM Tris, 100 mM NaCl, 1 mM DTT, 1 mM EDTA, pH 7.8). The eluted SC2M$^{pro}$ protein in buffer E was concentrated to 20 mg/mL and stored in -80°C for further use.

#### $IC_{50}$ analysis

The assays were carried out with 20 nM enzyme (except for MPI3, for which 10 nM enzyme was used) and 10 µM substrate at 37 °C with continuous shaking. All the analyses were carried out in triplicate. The substrate (DABCYL-Lys-Thr-Ser-Ala-Val-Leu-Gln-Ser-Gly-Phe-Arg-Lys-Met-Glu-EDANS) was purchased from Bachem and stored as 1 mM solution in 100% DMSO. Enzyme activity was monitored by fluorescence with excitation at 336 nm and emission at 455 nm wavelength. The dilution buffer (used for enzyme and substrate dilution) is 10 mM $Na_xH_yPO_4$ ,10 mM NaCl, 0.5 mM EDTA, pH 7.6. Final composition of the assay buffer is 10 mM $Na_xH_yPO_4$, 10 mM NaCl, 0.5 mM EDTA, 2 µM DTT (coming from enzyme stock solution), pH 7.6 with 1.25% DMSO. All the inhibitors were stored as 10 mM in 100% DMSO solutions in -20 °C freezer.

For $IC_{50}$ analysis, the inhibitor was diluted to 400-fold times higher than the highest working concentration to make the secondary stock solution (i.e. if the highest working concentration of inhibitor is 2 μM, then the inhibitor was diluted from its 10 mM stock solution to 800 μM in DMSO). 10 μL from this secondary stock solution was added to the 990 uL of dilution buffer. Serial dilutions were carried out in dilution buffer containing 1% DMSO to ensure all the inhibitor serial dilutions are at 1% DMSO. 25 μL of each inhibitor solution were added to 96-well plate with multichannel pipettor. Next, 25 μL of 80 nM enzyme solution (diluted from 10 μM enzyme storage solution in 10 mM $Na_xH_yPO_4$ ,10 mM NaCl, 0.5 mM EDTA, pH 7.6, 1 mM DTT with dilution buffer) were added by multichannel pipettor and mixed by pipetting up and down three times. Then, the enzyme-inhibitor solution was incubated at 37 $^oC$ for 30 minutes. During incubation period, 20 μM of the substrate solution is prepared by diluting from 1 mM stock solution with dilution buffer. When the incubation period is over, 50 μL of the 20 μM substrate solution added to each well by multichannel pipettor and the assay started. Data recording were stopped after 30 minutes. Data treatment were done with Graph Pad Prism 8.0 software. First 0-300 seconds were analyzed by linear regression for initial slope analyses. Then, the initial slopes were normalized and $IC_{50}$ values were determined by inhibitor vs response - Variable slope (four parameters).

### Crystallization of SC2M$^{pro}$

A freshly prepared SC2M$^{pro}$ protein solution at a concentration of 10 mg/mL was cleared by centrifugation at 14,000 rpm, 10 min. Next, a basic screen with the commercially available screening kits (Hampton Research Index$^{TM}$, Crystal Screen$^{TM}$ 1 and 2, PEGRx$^{TM}$ 1 and 2, PEG/Ion$^{TM}$ 1 and 2) were performed employing the sitting-drop vapor-diffusion method at 18°C. 1.0 μL of SC2M$^{pro}$ protein solution and 1.0 μL of reservoir buffer were mixed to equilibrate against 100 μL reservoir solution. Crystals appeared overnight under over 50 conditions. The most promising crystal was found under condition No.44 of PEG/Ion$^{TM}$ (0.2 M Ammonium phosphate dibasic, 20% w/v PEG3350, pH8.0). Subsequent optimization was performed by adjusting the temperature and concentration of protein and precipitant. The best plate-like crystals were obtained at 25°C from 0.2 M Ammonium phosphate dibasic, 17% w/v PEG3350, pH8.0, with a SC2M$^{pro}$ protein concentration of 14 mg/ml. Overnight growing crystals were washed with cryo-protectant containing mother liquor plus gradually increasing glycerol (5%, 10%, 15%, 20%, 25% and 30%). Cryo-protected crystals were fished for data collection.

### Crystallization of SC2M$^{pro}$ in complex with inhibitors

Soaking was performed to produce SC2M$^{pro}$-inhibitor complex crystals. Overnight growing SC2M$^{pro}$ crystals were washed with reservoir solution three times in situ. Subsequently, the crystals were washed three times with reservoir solution plus 0.5 mM inhibitor and 2% DMSO (Inhibitors were dissolved to 25 mM in 100% DMSO). The mixture was incubated at 25°C for 48 h. The cryo-protectant solution contained mother liquor plus 30% glycerol, 0.5 mM inhibitor and 2% DMSO. Cryo-protected crystals were fished for data collection.

### Data collection and Structure Determination

All data were collected on a Rigaku R-AXIS IV++ image plate detector and at the Advanced Light Source (ALS) beamline 5.0.2 using a Pilatus3 6M detector. The diffraction data were indexed, integrated and scaled with iMosflm.[1] All crystals are in space group C121. All the structures were determined by molecular replacement using the structure model of the free enzyme of the SARS-CoV-2 (2019-nCoV) main protease [Protein Data Bank (PDB) ID code 6Y2E] as the search model using Phaser in the Phenix package.[2-3] *JLigand* and *Sketcher* from the CCP4 suite were employed for the generation of PDB and geometric restraints for the inhibitors. The inhibitors were built into the $F_o$-$F_c$ density by using *Coot*.[4] Refinement of all the structures was performed with Real-space Refinement in Phenix.[3] Details of data quality and structure refinement are summarized in Table S1. All structural figures were generated with PyMOL (https://www.pymol.org).

**SARS-CoV-2 inhibition by a cell-based assay**

A slightly modified cytopathic effect (CPE)-based microneutralization assay was used to evaluate the drug efficacy against SARS-CoV-2 infection. Briefly, confluent African green monkey kidney cells (Vero E6) or human alveolar epithelial A549 cells stably expressing human ACE2 viral receptor, designated A549/hACE2, grown in 96-wells microtiter plates were pre-treated with serially 2-folds diluted individual drugs for two hours before infection with 100 or 500 infectious SARS-CoV-2 (US_WA-1 isolate) particles in 100 µL EMEM supplemented with 2% FBS, respectively. Cells pre-treated with parallelly diluted DMSO with or without virus were included as positive and negative controls, respectively. After cultivation at 37 °C for 3 (Vero E6) or 4 days (A549/hACE2), individual wells were observed under the microcopy for the status of virus-induced formation of CPE. The efficacy of individual drugs was calculated and expressed as the lowest concentration capable of completely preventing virus-induced CPE in 100% (EC100) or 50% (EC50) of the wells. All compounds were dissolved in 100% DMSO as 10 mM stock solutions before subjecting to dilutions with culture media.

### The synthesis of inhibitors MPI1-9

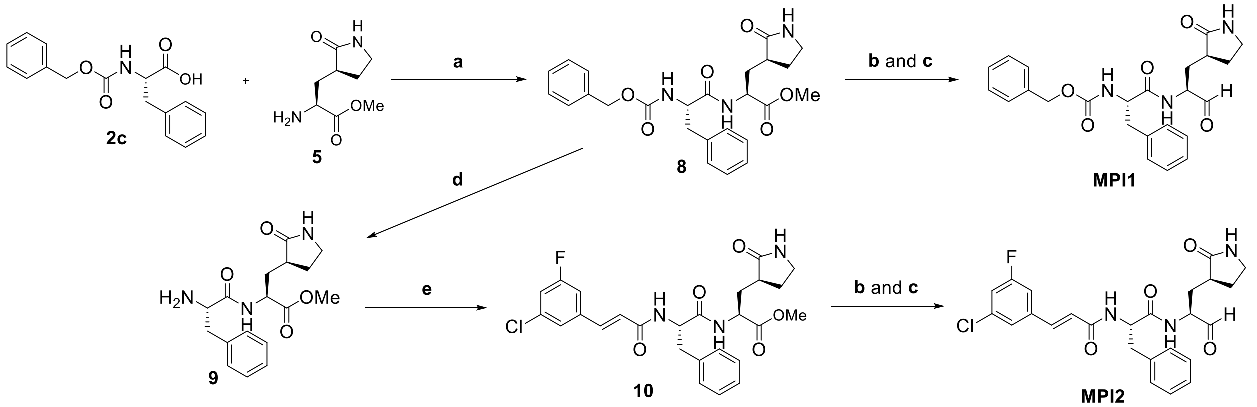


**Scheme S1.** The synthesis of compounds **MPI1** and **MPI2**. Reaction conditions: (a) HATU, DIPEA; (b) $LiBH_4$, THF; (c) Dess-Martin periodinane (DMP), $NaHCO_3$, $CH_2Cl_2$; (d) Pd/C, $H_2$; (e) (*E*)-4-Chloro-2-fluorocinnamic acid, HATU, DIPEA.

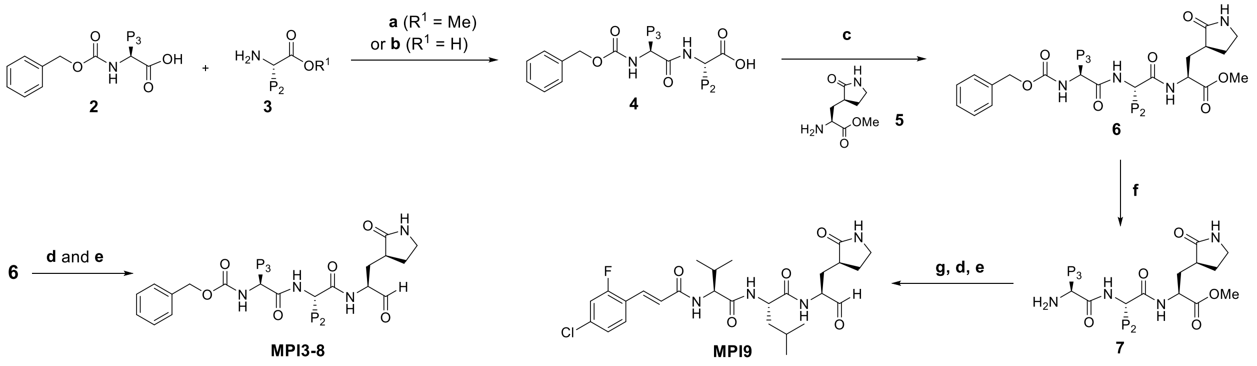


**Scheme S2.** The synthesis of compounds **MPI3-9.** Reaction conditions: (a) i) HATU, DIPEA; ii) LiOH; (b) $ClCO_2Et$, $Et_3N$; (c) HATU, DIPEA; (d) $LiBH_4$, THF; (e) Dess-Martin periodinane (DMP), $NaHCO_3$, $CH_2Cl_2$; (f) Pd/C, $H_2$; (g) (*E*)-4-Chloro-2-fluorocinnamic acid, HATU, DIPEA.

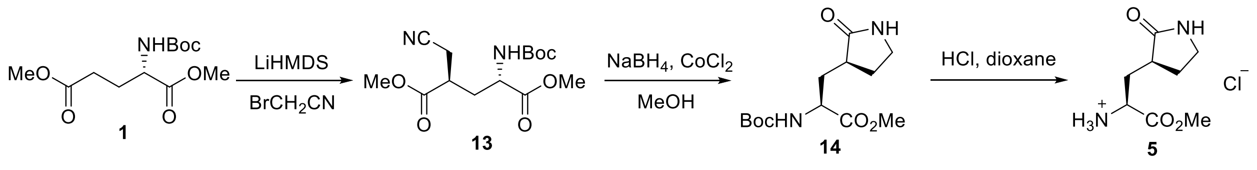


**Scheme S3.** The synthesis of compound **5**.[5]

#### Dimethyl (2S,4R)-2-((tert-butoxycarbonyl)amino)-4-(cyanomethyl)pentanedioate (13)

A solution of *N*-Boc-glutamic acid dimethyl ester (3 g, 11 mmol, 1 equiv.) in anhydrous THF (20 mL) was cooled under -78 °C. Then 24 mL of 1 M LiHMDS solution in THF (24 mmol, 2.18 equiv.) was added to the solution dropwise. After addition, the solution was stirred under -78 °C for 1 h. Meanwhile, boromoacetonitrile was stirred with activated basic alumina for 2 h and then filtered. Freshly dried and filtered bromoacetonitrle (1.4 g, 11.8 mmol, 1.06 equiv.) was then added dropwise to the dianion solution. The solution was then stirred under -78 °C for 3~5 h, until TLC confirms complete consumption of the starting material. Then the reaction was quenched with pre-cooled methanol (1 mL) in one portion and stirred under the same temperature for 30 min. The methoxide solution was then quenched with pre-cooled AcOH/THF (1 mL in 6 mL THF) in one portion and stirred for another 30 min under the same temperature. Then the cooling bath was removed. The reaction mixture was allowed to warm up to room temperature and poured into 50 mL of saturated brine solution. The layers were separated, and the organic layer was then concentrated to give dark oil. Then to the residue was added 4 g of silica gel, 1 g of activated charcoal and 50 mL of dichloromethane. The slurry was stirred for 1 h, and then filtered and washed with another 50 mL of dichloromethane. The filtrate was then concentrated to give brown oil, which was used without further purification.

**Methyl (S)-2-((tert-butoxycarbonyl)amino)-3-((S)-2-oxopyrrolidin-3-yl)propanoate (14)**

To a pre-cooled solution of $CoCl_2 \cdot 6H_2O$ (1.54 g, 6.5 mmol) and **13** (11 mmol, crude) in methanol under 0 °C was added $NaBH_4$ (44 mmol, 1.67 g) in portions over 30 min. The reaction was exothermic and produces copious amount of hydrogen and black precipitate. The reaction mixture was stirred under room temperature for 24 h, and then concentrated on vacuo. The residue oil was then poured into 10% citric acid and filtered. The filtrate was then extracted with ethyl acetate twice. The organic layer was then dried over anhydrous $Na_2SO_4$ and concentrated. The residue was purified by flash chromatography to afford **14**as light-yellow oil (2.1 g, 66%). $^1$H NMR (400 MHz, $CDCl_3$): δ 6.58 (s, 1H), 5.58 (d, J = 8.5 Hz, 1H), 4.19-4.36 (m, 1H), 3.71 (s, 3H), 3.23-3.39 (m, 2H), 2.36-2.54 (m, 2H), 2.04-2.19 (m, 1H), 1.73-1.90 (m, 1H), 1.41 (s, 9H).

**(S)-1-Methoxy-1-oxo-3-((S)-2-oxopyrrolidin-3-yl)propan-2-amine hydrochloride (5)**

To a solution of **14** (2.1 g, 7.3 mmol)in 1,4-dioxane (10 mL) was added dropwise a HCl solution in 1,4-dioxane (4 M, 10 mL). The resulting solution was stirred at room temperature for 1 h. Then residue was then concentrated *on vacuo* to afford **5** as light-yellow hydroscopic crystal (1.5 g, 92%). $^1$H NMR (400 MHz, $d_6$-DMSO): δ 8.72 (s, 3H), 7.97 (s, 1H), 4.13-4.24 (m, 1H), 3.76 (s, 3H), 3.12-3.24 (m, 2H), 2.54-2.65 (m, 1H), 2.23-2.34 (m, 1H), 2.01-2.10 (m, 1H), 1.83-1.92 (m, 1H), 1.62-1.73 (m, 1H).

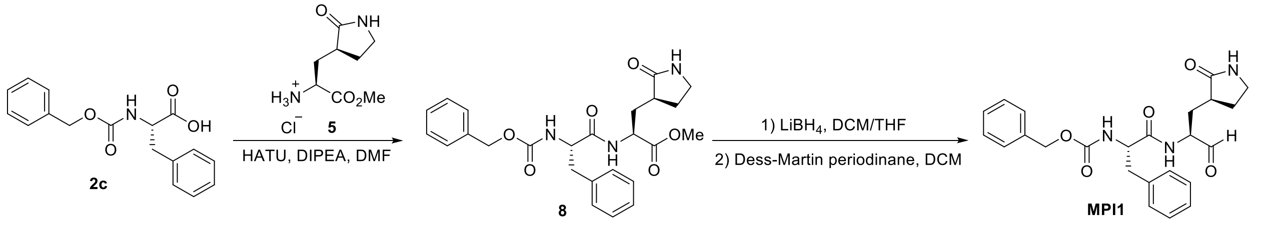


**Scheme S4.** The synthesis of compound **MPI1**.

**Methyl (*S*)-2-((*S*)-2-(((benzyloxy)carbonyl)amino)-3-phenylpropanamido)-3-((*S*)-2-oxopyrrolidin-3-yl)propanoate (8)**

To a solution of **2c** (2 mmol, 0.44 g) and **5** (2 mmol, 0.44 g) in anhydrous DMF (10 mL) was added DIPEA (4 mmol, 0.52 g) and was cooled to 0 °C. HATU (2.2 mmol, 0.84 g) was added to the solution under 0 °C and then stirred at room temperature overnight. The reaction mixture was then diluted with ethyl acetate (50 mL) and washed with saturated $NaHCO_3$ solution (2×20 mL), 1 M HCl solution (2×20 mL), and saturated brine solution (2×20 mL) sequentially. The organic layer was dried over anhydrous $Na_2SO_4$ and then concentrated *on vacuo*. The residue was then purified with flash chromatography (50-100% EtOAc in hexanes as the eluent) to afford **8** as white solid (520 mg, 56%). $^1$H NMR (400 MHz, $d_6$-DMSO): δ 8.59 (d, J = 7.9 Hz, 1H), 7.66 (s, 1H), 7.53 (d, J = 8.5 Hz, 1H), 7.03-7.43 (m, 10H), 4.93 (q, J = 12.3, 11.8 Hz, 2H), 4.31-4.42 (m, 1H), 4.22-4.31 (m, 1H), 3.63 (s,3H), 2.91-3.18 (m, 3H), 2.68-2.79 (m, 1H), 2.23-2.36 (m, 1H), 2.01-2.17 (m, 2H), 1.53-1.67 (m, 2H).

**Benzyl ((*S*)-1-oxo-1-(((S)-1-oxo-3-((*S*)-2-oxopyrrolidin-3-yl)propan-2-yl)amino)-3-phenylpropan-2-yl)carbamate (MPI1)**

To a solution of **8** (0.1 mmol, 47 mg) in anhydrous dichloromethane (5 mL) was added a solution of $LiBH_4$ in anhydrous THF (2 M, 0.1 mL, 0.2 mmol) at 0 °C. The resulting solution was stirred at the same temperature for 3 h. Then a saturated solution of $NH_4Cl$ (5 mL) was added dropwise to quench the reaction. The layers were separated, and the organic layer was washed with saturated brine solution (2×10 mL), dried over anhydrous $Na_2SO_4$. and evaporated to dryness. The residue was then dissolved in anhydrous dichloromethane (5 mL) and cooled to 0 °C. Dess-Martin periodinane (0.2 mmol, 85 mg) was added to the solution. The reaction mixture was then stirred at room temperature overnight. Then the reaction was quenched with a saturated $NaHCO_3$ solution containing 10% $Na_2S_2O_3$. The layers were separated. The organic layer was then washed with saturated brine solution (2×10 mL), dried over anhydrous $Na_2SO_4$ and evaporated *on vacuo*. The residue was then purified with flash chromatography (1-10% methanol in dichloromethane as the eluent) to afford **1i** as white solid (30 mg, 65%). $^1$H-NMR (400 MHz, $CDCl_3$): δ 9.26 (s, 1H), 8.18 (s, 1H), 7.46 – 7.08 (m, 10H), 5.54 (s, 1H), 5.44 (d, J = 9.2 Hz, 1H), 5.11 (s, 2H), 4.65 – 4.53 (m, 1H), 4.30 – 4.16 (m, 1H), 3.37 – 3.24 (m, 2H), 3.23 – 3.14 (m, 1H), 3.07 (dd, J = 13.6, 6.5 Hz, 1H), 2.40 – 2.28 (m, 1H), 2.27 – 2.19 (m, 1H), 1.92 – 1.73 (m, 3H); $^{13}$C NMR (100 MHz, $CDCl_3$) δ 199.98, 180.16, 172.13, 155.96, 136.39, 129.62, 128.73, 128.65, 128.29, 128.15, 127.14, 67.11, 58.05, 56.14, 40.72, 38.96, 38.17, 31.08, 29.53, 28.88; ESI-MS calcd for $C_{24}H_{28}N_3O_5$ $(M+H^+)$: 438.2; found 438.3.

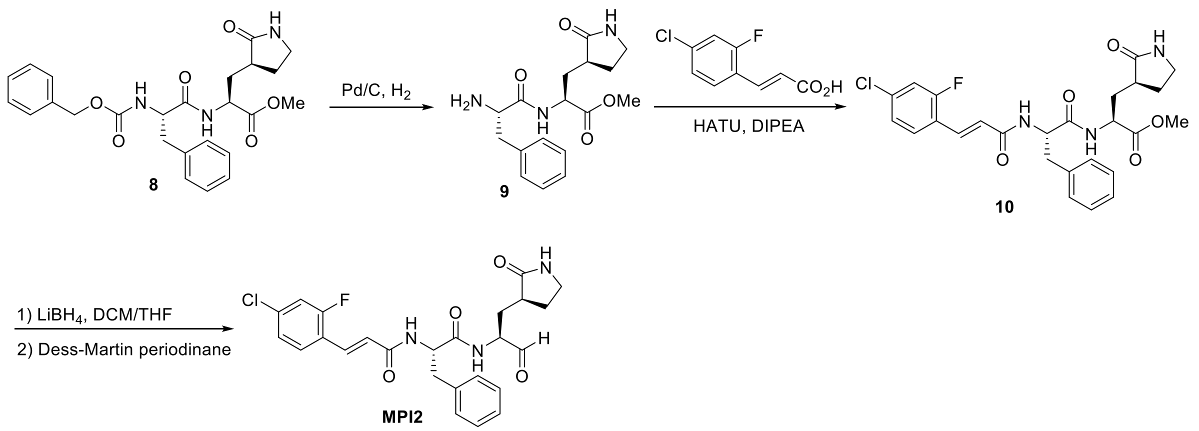


**Scheme S5.** The synthesis of compound **MPI2**.

#### Methyl (*S*)-2-((*S*)-2-((*E*)-3-(3-chloro-5-fluorophenyl)acrylamido)-3-phenylpropanamido)-3-((*S*)-2-oxopyrrolidin-3-yl)propanoate (10)

To a solution of **8** (0.25 mmol, 116 mg) in methanol was added 10 % Pd/C (26 mg). The mixture was then stirred with hydrogen balloon at room temperature for 3 h. The catalyst was then filtered off and the solution was evaporated on vacuo to afford **9** as white solid, which was used without purification. To a solution of **9** in 2 mL dry DMF was added DIPEA (0.5 mmol, 65 mg) and cooled to 0 °C. Then HATU (0.3 mmol, 114 mg) was added to the solution at the same temperature. The solution was stirred at room temperature overnight. The reaction mixture was then diluted with ethyl acetate (20 mL) and washed with saturated $NaHCO_3$ solution (2×10 mL), 1 M HCl solution (2×10 mL), and saturated brine solution (2×10 mL) sequentially. The organic layer was dried over anhydrous $Na_2SO_4$ and then concentrated *on vacuo*. The residue was then purified with flash chromatography (1-10% methanol in dichloromethane as the eluent) to afford **10** as white solid (80 mg, 62%). $^1$H-NMR (400 MHz, $CDCl_3$): δ 7.92 (d, J = 6.7 Hz, 1H), 7.62 (d, J = 15.8 Hz, 1H), 7.40 (t, J = 8.3 Hz, 1H), 7.32 – 7.17 (m, 5H), 7.11 (td, J = 8.7, 7.4, 3.0 Hz, 2H), 6.61 – 6.43 (m, 2H), 5.89 (s, 1H), 5.02 (dd, J = 8.2, 5.9 Hz, 1H), 4.51 – 4.33 (m, 1H), 3.72 (s, 3H), 3.38 – 3.25 (m, 2H), 3.25 – 3.11 (m, 2H), 2.44 – 2.31 (m, 1H), 2.27 – 2.19 (m, 1H), 2.17 – 2.05 (m, 1H), 1.95 – 1.74 (m, 2H).

#### (*E*)-3-(3-chloro-5-fluorophenyl)-*N*-((*S*)-1-oxo-1-(((*S*)-1-oxo-3-((*S*)-2-oxopyrrolidin-3-yl)propan-2-yl)amino)-3-phenylpropan-2-yl)acrylamide (MPI2)

To a solution of **10** (0.1 mmol, 52 mg) in anhydrous dichloromethane (5 mL) was added a solution of $LiBH_4$ in anhydrous THF (2 M, 0.1 mL, 0.2 mmol) at 0 °C. The resulting solution was stirred at the same temperature for 3 h. Then a saturated solution of $NH_4Cl$ (5 mL) was added dropwise to quench the reaction. The layers were separated, and the organic layer was washed with saturated brine solution (2×10 mL), dried over anhydrous $Na_2SO_4$. and evaporated to dryness. The residue was then dissolved in anhydrous dichloromethane (5 mL) and cooled to 0 °C. Dess-Martin periodinane (0.2 mmol, 85 mg) was added to the solution. The reaction mixture was then stirred at room temperature overnight. Then the reaction was quenched with a saturated $NaHCO_3$ solution containing 10% $Na_2S_2O_3$. The layers were separated. The organic layer was then washed with saturated brine solution (2×10 mL), dried over anhydrous $Na_2SO_4$ and evaporated *on vacuo*. The residue was then purified with flash chromatography (1-10% methanol in dichloromethane as the eluent) to afford **MPI2** as white solid (27 mg, 55%). $^1$H-NMR (400 MHz, DMSO-$d_6$): δ 9.30 (s, 1H), 8.67 (d, J = 7.6 Hz, 1H), 8.58 (d, J = 8.2 Hz, 1H), 7.68 – 7.61 (m, 2H), 7.52 (dd, J = 10.8, 2.1 Hz, 1H), 7.41 – 7.32 (m, 2H), 7.30 – 7.15 (m, 5H), 6.81 (d, J = 16.0 Hz, 1H), 4.75 – 4.67 (m, 1H), 4.21 – 4.12 (m, 1H), 3.18 – 3.04 (m, 3H), 2.89 (dd, J = 13.8, 9.3 Hz, 1H), 2.25 – 2.05 (m, 2H), 1.88 (ddt, J = 13.9, 11.3, 5.6 Hz, 1H), 1.67 – 1.55 (m, 2H); $^{13}$C NMR (101 MHz, CDCl3) δ 199.87, 179.97, 171.97, 165.18, 162.26, 159.71, 136.30, 133.45, 130.13, 129.55, 128.59, 127.05, 124.96, 123.37, 121.40, 116.82, 57.99, 54.27, 40.62, 38.83, 38.10, 29.42, 28.75; ESI-MS: calcd for $C_{25}H_{26}ClFN_3O_4$ (M+H$^+$): 486.1; found 486.1.

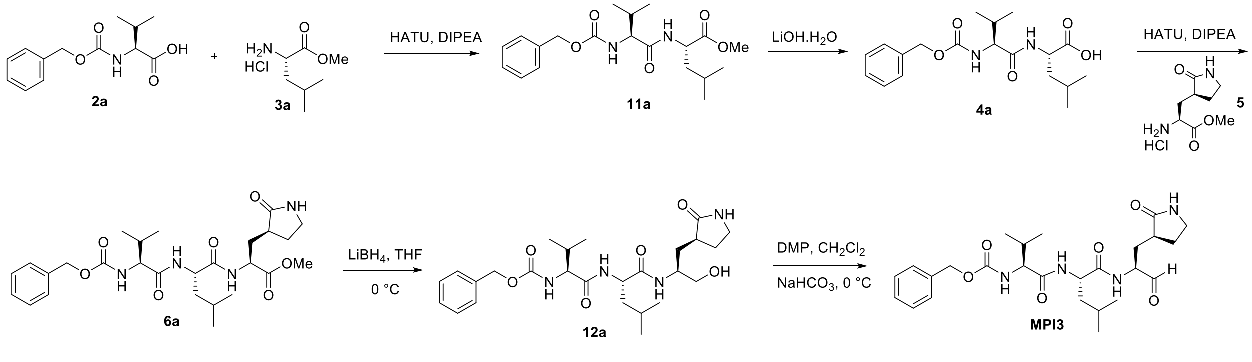


**Scheme S6.** The synthesis of compound **MPI3**.

#### (*S*)-Methyl 2-((*S*)-2-(((benzyloxy)carbonyl)amino)-3-methylbutanamido)-4-methylpentanoate (11a)

The amino acid methyl ester hydrochloride **3a** (1.0 g, 5.52 mmol) and the Cbz-protected amino acid **2a** (1.88 g, 6.08 mmol) were dissolved in dry DMF (20 mL) and the reaction was cooled to 0 °C. HATU (2.52 g, 6.62 mmol) and DIPEA (3.92 mL, 22.08 mmol) were added, and the reaction mixture was allowed warm up to room temperature and stirred for 12 h. The mixture was then poured into water (50 mL) and extracted with ethyl acetate (4×20 mL). The organic layer was washed with aqueous hydrochloric acid 10% v/v (2×20 mL), saturated aqueous $NaHCO_3$ (2×20 mL), brine (2×20 mL) and dried over $Na_2SO_4$. The organic phase was evaporated to dryness and the crude material purified by silica gel column chromatography (15-50% EtOAc in n-hexane as the eluent) to afford **11a** white solid (1.82, 69%). $^1$H NMR ($CDCl_3$, 400MHz) δ 7.28-7.22 (m, 5H), 6,28 (d, *J*=7.72 Hz, 1H), 5.35 (d, =864 Hz, 1H), 5.03 (s, 2H), 4.56-451 (m, 1H), 3.97 (t, *J*=8.12 Hz, 1H), 3.65 (s, 3H), 2.19-1.98 (m, 1H), 1.62-1.43 (m, 3H), 0.92-0.83 (m, 12H); $^{13}$C NMR ($CDCl_3$, 100MHz) δ 173.2, 171.1, 156., 136.2, 128.5 (2C), 128.2, 128 (2C), 67.0, 60.2, 52.3, 50.7, 1., 31.3, 2.8, 22.7, 21.9, 19.1, 17.8.

#### (*S*)-2-((*S*)-2-(((Benzyloxy)carbonyl)amino)-3-methylbutanamido)-4-methylpentanoic acid (4a)

The peptide **11a** (500 mg, 1.14 mmol) was dissolved in THF/$H_2O$ (1:1, 10 mL).LiOH (114 mg, 2.86 mmol) was added at 0 °C. The mixture was stirred at room temperature overnight. Then THF was removed *on vacuum* and the aqueous layer was acidified with 1 M HCl and extracted with dichloromethane (3 x 10 mL). The organic layer was dried over anhydrous $Na_2SO_4$ and concentrated to yield **4a** as white solid (315 mg, 65%). $^1$H NMR ($CDCl_3$, 400 MHz): δ 7.26-7.23 (m, 5H), 6.65 (d, *J*=7.88, 1H), 5.68 (d, *J*=4.64 Hz, 1H), 5.03 (s, 2H), 4.56-4.45 (m, 1H), 3.97 (t, *J*=7.92 Hz, 1H), 2.02-1.96 (m, 1H), 1.66-1.46 (m, 3H), 0.85 (dd, *J*= 7.36, 13.3 Hz, 12H); $^{13}$C NMR ($CDCl_3$, 100 MHz): δ 176.0, 171.8, 156.7, 136.1, 128.5 (2C), 128.2, 128.0 (2C), 67.2, 60., 50.8, 41.1, 31.1, 24.8, 22.8, 21.8, 19.1, 18.0.

**(5*S*,8*S*,11*S*)-Methyl 8-isobutyl-5-isopropyl-3,6,9-trioxo-11-(((S)-2-oxopyrrolidin-3-yl)methyl)-1-phenyl-2-oxa-4,7,10-triazadodecan-12-oate (6a)**

The methyl (S)-2-amino-3-((S)-2-oxopyrrolidin-3-yl)propanoate hydrochloride **5** (150 mg, 0.657 mmol) and the peptide **11a** (270 mg, 0.743 mmol) were dissolved in dry DMF (10 mL) and the reaction was cooled to 0 °C. HATU (308 mg, 0.788 mmol) and DIPEA (0.48 mL, 2.63 mmol) were added, and the reaction mixture was allowed warm up to room temperature and stirred for 12 h. The mixture was then poured into water (20 mL) and extracted with ethyl acetate (4×20 mL). The organic layer was washed with aqueous hydrochloric acid 10% v/v (2×20 mL), saturated aqueous $NaHCO_3$ (2×20 mL), brine (2×20 mL) and dried over $Na_2SO_4$. The organic phase was evaporated to dryness and the crude material purified by silica gel column chromatography afford **6a**as white solid (250 mg, 70%). $^1$H NMR ($CDCl_3$, 400 MHz) δ 7.89 (d, *J*=7.16 Hz, 1H), 7.28-7.20 (m, 5H), 7.00 (d, *J*=8.24 Hz, 1H), 6.65 (Brs, 1H), 5.47 (d, *J*=8.8 Hz, 1H), 5.01 (s, 2H), 4.59-4.51 (m, 1H), 4.46-4.38 (m, 1H), 3.94 (t, *J*=7.84 Hz, 1H), 3.63 (s, 3H), 3.28-3.18 (m, 2H), 2.38-2.22 (m, 2H), 2.29-1.98 (m, 2H), 1.79-1.70 (m, 1H), 1.68-1.51 (m, 2H), 1.50-1.25 (m, 2H), 0.86-0.81 (m, 12H);$^{13}$C NMR ($CDCl_3$, 100 MHz) δ 179.8, 172.5, 172.1, 171.2, 156.5, 136.2, 128.5 (2C), 128.2, 128.0 (2C), 67.1, 60.5, 52.4, 51.7, 51.1, 42.0, 40.5, 38.3, 33.1, 31.1, 28.1, 24.6, 22.8, 22.0, 19.7, 19.1.

**Benzyl ((S)-1-(((S)-1-(((S)-1-hydroxy-3-((S)-2-oxopyrrolidin-3-yl)propan-2-yl)amino)-4-methyl-1-oxopentan-2-yl)amino)-3-methyl-1-oxobutan-2-yl)carbamate (12a)**

To a stirred solution of compound **6a** (120 mg, 0.254 mmol) in THF (8 mL) was added $LiBH_4$ (2.0 M in THF, 0.636 mL, 1.27 mmol) in several portions at 0 °C under a nitrogen atmosphere. The reaction mixture was stirred at 0 °C for 1 h, then allowed to warm up to room temperature, and stirred for an additional 2 h. The reaction was quenched by the drop wise addition of 1.0 M HCl (aq) (1.2 mL) with cooling in an ice bath. The solution was diluted with ethyl acetate and $H_2O$. The phases were separated, and the aqueous layer was extracted with ethyl acetate (3×15 mL). The organic phases were combined together, dried over $MgSO_4$, filtered, and concentrated on a rotorvap to give a yellow oily residue. Column chromatographic purification of the residue (6% MeOH in CH2Cl2 as the eluent) afforded a white solid (80 mg, 70%).$^1$H NMR ($CDCl_3$) δ 7.62 (d, *J*=6.48 Hz, 1H), 7.30-7.27 (m, 5H), 6.57 (d, *J*=7.6 Hz, 1H), 5.76 (Brs, 1H), 5.30 (d, *J*=7.84 Hz, 1H), 5.05 (s, 2H), 4.4-4.38 (m, 1H), 3.93 (t, *J*=7.0 Hz, 2H), 3.60-3.53 (m, 2H), 3.43 (s, 3H), 3.27-3.25 (m, 2H), 2.39-2.32 (m, 2H), 2.15-2.01 (m, 1H), 1.95-1.75 (m, 3H), 1.68-1.49 (m, 6H), 0.92-0.84 (m, 12H).

**Benzyl ((S)-3-methyl-1-(((S)-4-methyl-1-oxo-1-(((S)-1-oxo-3-((S)-2-oxopyrrolidin-3-yl)propan-2-yl)amino)pentan-2-yl)amino)-1-oxobutan-2-yl)carbamate (MPI3)**

To a solution of **12a** (70 mg, 0.142 mmol) in $CH_2Cl_2$ (6 mL) was added $NaHCO_3$ (48 mg, 4 equiv) and the Dess-Martin reagent (180 mg, 0.427 mmol, 3 equiv). The resulting mixture was stirred at rt for 12 h. Then the reaction was quenched with a saturated $NaHCO_3$ solution containing 10 % $Na_2S_2O_3$. The layers were separated. The organic layer was then washed with saturated brine solution, dried over anhydrous $Na_2SO_4$ and concentrated *on vacuum*. The residue was then purified with flash chromatography afford **MPI3** as white solid (45 mg, 64%). $^1$H NMR ($CDCl_3$, 400 MHz): δ 9.41 (s, 1H), 8.18 (d, *J*=5.12 Hz, 1H), 7.28-7.26 (m, 5H), 6.63 (d, *J*=7.0 Hz, 1H), 6.15 (Brs, 1H), 5.40 (d, *J*=6.88 Hz, 1H), 5.02 (s, 2H), 4.52-4.50 (m, 1H), 4.27-4.25 (m, 1H), 3.94-3.92 (m, 2H), 3.27-3.23 (m, 2H), 2.35-2.27 (m, 2H), 2.07-2.05 (m, 1H), 1.90-1.81 (m, 2H), 1.76-1.71 (m, 1H), 1.57-1.43 (m, 3H), 0.85 (dd, *J*=6.76, 14.72 Hz, 12H);$^{13}$C NMR ($CDCl_3$, 100 MHz): δ 199.6, 180.0, 173.2, 171.4, 156.6, 136.2, 128.6 (2C), 128.2, 128.1 (2C), 67.1, 60.6, 57.5, 51.2, 41.7, 40.6, 38.0, 31.0, 29.8, 28.4, 24.8, 22.9, 21.9, 19.2, 17.8.

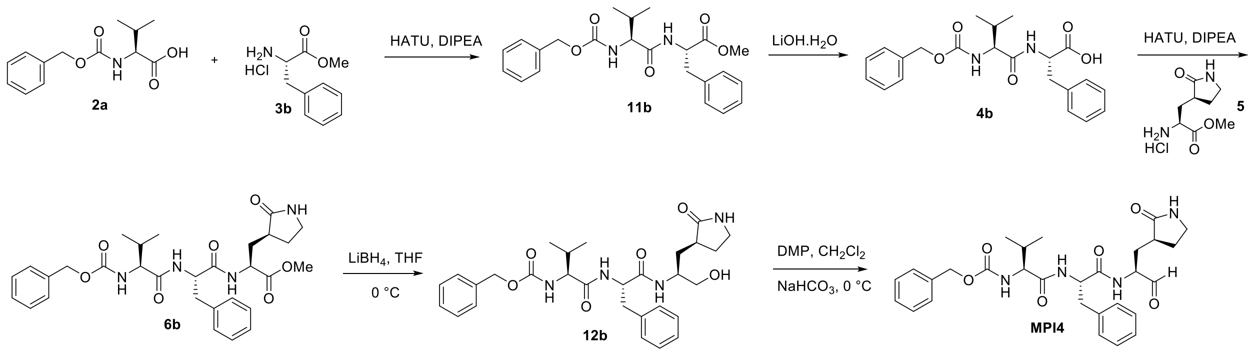


**Scheme S7.** The synthesis of compound **MPI4**.

**Methyl ((benzyloxy)carbonyl)-L-valyl-L-phenylalaninate (11b)**

To a solution of **2a** (5 mmol, 1.25 g) and **3b** (5 mmol, 1.07g) in anhydrous DMF (20 mL) was added DIPEA (10 mmol, 1.29 g) and was cooled to 0 °C. HATU (5.5 mmol, 2.09 g) was added to the solution under 0 °C and then stirred at room temperature overnight. The reaction mixture was then diluted with ethyl acetate (100 mL) and washed with saturated $NaHCO_3$ solution (2×50 mL), 1 M HCl solution (2×50 mL), and saturated brine solution (2×50 mL) sequentially. The organic layer was dried over anhydrous $Na_2SO_4$ and then concentrated *on vacuo*. The residue was then purified with flash chromatography (15-50% EtOAc in hexanes as the eluent) to afford **11b** as white soild (1.52 g, 74%).

**((Benzyloxy)carbonyl)-L-valyl-L-phenylalanine (4b)**

**11b** (1 mmol, 470 mg) was dissolved in 5 mL of THF. A solution of LiOH·$H_2O$ (2 mmol, 84 mg) in 5 mL $H_2O$ was added to the solution. The mixture was stirred at room temperature overnight. Then THF was removed *on vacuo* and the aqueous layer was acidified with 1 M HCl and extracted with dichloromethane (3×10 mL). The organic layer was dried over anhydrous $Na_2SO_4$ and evaporated to give **4b** as white solid (312 mg, 76 %). $^1$H-NMR (400 MHz, $CD_3OD$): δ 7.29-7.42

(m, 5H), 7.16-7.29 (m, 5H), 5.11 (s, 2H), 4.73 – 4.68 (m, 1H), 3.94 (d, J = 7.3 Hz, 1H), 3.21 (dd, J = 13.9, 5.2 Hz, 1H), 3.01 (dd, J = 13.9, 8.6 Hz, 1H), 2.06 – 1.95 (m, 1H), 0.91 (dd, J = 8.5, 6.7 Hz, 6H).

**Methyl (5S,8S,11S)-8-benzyl-5-isopropyl-3,6,9-trioxo-11-(((S)-2-oxopyrrolidin-3-yl)methyl)-1-phenyl-2-oxa-4,7,10-triazadodecan-12-oate (6b)**

To a solution of **4b** (0.4 mmol, 160 mg) and **5** (0.4 mmol, 88 mg) in anhydrous DMF (2 mL) was added DIPEA (0.8 mmol, 103 mg) and was cooled to 0 °C. HATU (0.44 mmol, 167 mg) was added to the solution under 0 °C and then stirred at room temperature overnight. The reaction mixture was then diluted with ethyl acetate (20 mL) and washed with saturated $NaHCO_3$ solution (2×10 mL), 1 M HCl solution (2×10 mL), and saturated brine solution (2×10 mL) sequentially. The organic layer was dried over anhydrous $Na_2SO_4$ and then concentrated *on vacuo*. The residue was then purified with flash chromatography (1-10% methanol in dichloromethane as the eluent) to afford **6b** as white solid (151 mg, 67%). $^1$H-NMR (400 MHz, $CDCl_3$): δ 7.71 (d, J = 7.1 Hz, 1H), 7.42 – 7.30 (m, 5H), 7.25 – 7.13 (m, 5H), 6.78 (d, J = 8.5 Hz, 1H), 5.83 (s, 1H), 5.26 (d, J = 8.7 Hz, 1H), 5.10 (d, J = 4.2 Hz, 2H), 4.83 (q, J = 6.9 Hz, 1H), 4.51 – 4.41 (m, 1H), 3.96 (dd, J = 8.6, 6.2 Hz, 1H), 3.70 (s, 3H), 3.37 – 3.25 (m, 2H), 3.11 (d, J = 6.3 Hz, 2H), 2.44 – 2.31 (m, 1H), 2.26 – 2.13 (m, 1H), 2.13 – 1.99 (m, 2H), 1.93 – 1.74 (m, 2H), 0.91 (d, J = 6.8 Hz, 3H), 0.82 (d, J = 6.8 Hz, 3H).

**Benzyl ((S)-3-methyl-1-oxo-1-(((S)-1-oxo-1-(((S)-1-oxo-3-((S)-2-oxopyrrolidin-3-yl)propan-2-yl)amino)-3-phenylpropan-2-yl)amino)butan-2-yl)carbamate (MPI4)**

To a solution of **6b** (0.1 mmol, 57 mg) in anhydrous dichloromethane (5 mL) was added a solution of $LiBH_4$ in anhydrous THF (2 M, 0.1 mL, 0.2 mmol) at 0 °C. The resulting solution was stirred at the same temperature for 3 h. Then a saturated solution of $NH_4Cl$ (5 mL) was added dropwise to quench the reaction. The layers were separated, and the organic layer was washed with saturated brine solution (2×10 mL), dried over anhydrous $Na_2SO_4$. and evaporated to dryness. The residue was then dissolved in anhydrous dichloromethane (5 mL) and cooled to 0 °C. Dess-Martin periodinane (0.2 mmol, 85 mg) was added to the solution. The reaction mixture was then stirred at room temperature overnight. Then the reaction was quenched with a saturated $NaHCO_3$ solution containing 10 % $Na_2S_2O_3$. The layers were separated. The organic layer was then washed with saturated brine solution (2×10 mL), dried over anhydrous $Na_2SO_4$ and evaporated *on vacuo*. The residue was then purified with flash chromatography (1-10% methanol in dichloromethane as the eluent) to afford **MPI4** as white solid (30 mg, 65 %). $^1$H-NMR (400 MHz, $CDCl_3$): δ 9.26 (s, 1H), 8.18 (s, 1H), 7.46 – 7.08 (m, 10H), 5.54 (s, 1H), 5.44 (d, J = 9.2 Hz, 1H), 5.11 (s, 2H), 4.65 – 4.53 (m, 1H), 4.30 – 4.16 (m, 1H), 3.37 – 3.24 (m, 2H), 3.23 – 3.14 (m, 1H), 3.07 (dd, J = 13.6, 6.5 Hz, 1H), 2.40 – 2.28 (m, 1H), 2.27 – 2.19 (m, 1H), 1.92 – 1.73 (m, 3H); $^{13}$C NMR (100 MHz, $CDCl_3$) δ 199.98, 180.16, 172.13, 155.96, 136.39, 129.62, 128.73, 128.65, 128.29, 128.15, 127.14, 67.11, 58.05, 56.14, 40.72, 38.96, 38.17, 31.08, 29.53, 28.88; ESI-MS calcd for $C_{24}H_{28}N_3O_5$ ($M+H^+$): 438.2; found 438.3.

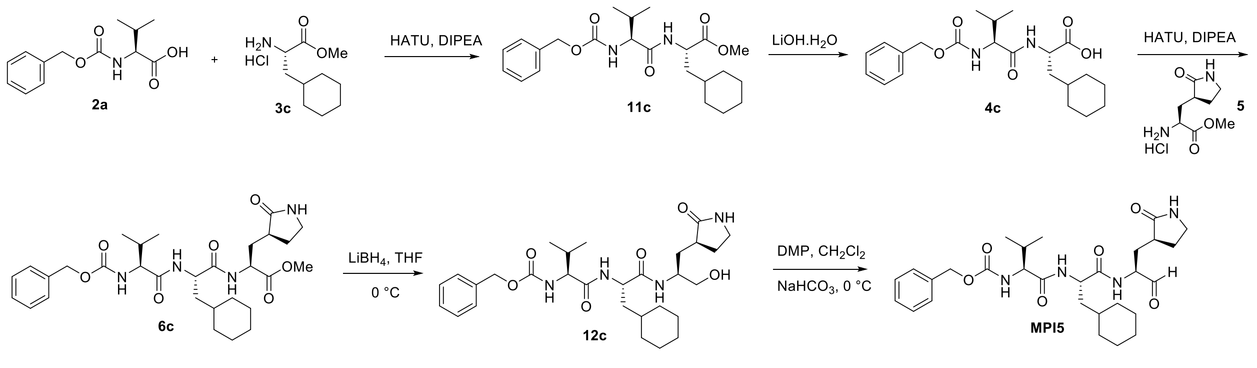

**Scheme S8.** The synthesis of compound **MPI5**.

**Methyl (S)-2-((S)-2-(((benzyloxy)carbonyl)amino)-3-methylbutanamido)-3-cyclohexylpropanoate (11c)**.

To a solution of **2a** (2g, 7.95 mmol, 1.0 equiv) in anhydrous DMF (15 mL) at 0°C, and then**3c** (1.8 g, 7.95 mmol, 1,0 equiv), HATU (4.5 g, 12.0 mmol, 1.5 equiv), DIPEA (7.0 mL, 40.0 mmol, 5.0 equiv) was added sequentially. The mixture was stirred at room temperature for 6 h. The mixture was diluted with EtOAc and washed with water, 1M HCl, sat. NaCl, dried over $Na_2SO_4$, and concentrated. The residue was purified by column chromatography (EtOAc: Hexane = 1:2 v/v) to afford the pure product **11c** as a white solid (2.7 g, 81%). $^1H$ NMR (400 MHz, $CD_3OD$-$d_4$) δ 7.4 – 7.3 (m, 5H), 5.1 (d, $J$ = 1.6 Hz, 2H), 4.5 (dd, $J$ = 9.6, 5.6 Hz, 1H), 4.0 (d, $J$ = 7.5 Hz, 1H), 3.7 (s, 3H), 2.2 – 2.0 (m, 1H), 1.8 – 1.6 (m, 7H), 1.4 (tdd, $J$ = 11.0, 6.9, 3.6 Hz, 1H), 1.3 – 1.1 (m, 3H), 1.0 (dd, $J$ = 11.7, 6.7 Hz, 7H), 0.9 – 0.8 (m, 1H). $^{13}C$ NMR (100 MHz, $CD_3OD$-$d_4$) δ 174.5, 174.3, 158.5, 138.2, 129.4, 129.4, 129.0, 128.8, 128.8, 67.6, 61.9, 52.5, 51.3, 39.9, 35.2, 34.7, 33.1, 32.0, 27.5, 27.3, 27.1, 19.7, 18.7.

**(S)-2-((S)-2-(((benzyloxy)carbonyl)amino)-3-methylbutanamido)-3-cyclohexylpropanoic acid (4c).**

To a solution of **11c** (400 mg, 1.2 mmol, 1.0 equiv) in 1:1 THF/$H_2O$ (8 mL) was added LiOH·$H_2O$ (200 mg, 4.8 mmol, 4.0 equiv). The reaction was stirred at RT for 2 h. After completion, the reaction mixture was neutralized with 1M HCl solution and extracted with EtOAc. The organic layer was washed with sat. NaCl, dried over $Na_2SO_4$ and concentrated to afford the product **4c** (310 mg, yield 80%) as a white solid. The residue was used in the next without further purification.

**Methyl (5S,8S,11S)-8-(cyclohexylmethyl)-5-isopropyl-3,6,9-trioxo-11-(((S)-2-oxopyrrolidin-3-yl)methyl)-1-phenyl-2-oxa-4,7,10-triazadodecan-12-oate (6c).**

To a solution of **4c** (300 mg, 0.74 mmol, 1.0 equiv) in anhydrous DMF (5 mL) at 0°C, and then **5** (165 mg, 0.74 mmol, 1.0 equiv), HATU (400 mg, 1.05 mmol, 1.5 equiv), DIPEA (610 µL, 3.7 mmol, 5.0 equiv) was added sequentially. The mixture was stirred at RT for 6 h. The mixture was diluted with EtOAc and washed with water, 1M HCl, sat. NaCl, dried over $Na_2SO_4$, and concentrated. The residue was purified by column chromatography (MeOH: DCM = 1:20 v/v) to afford the pure product **6c** as a white solid (250 mg, 60%).

**Benzyl ((S)-1-(((S)-3-cyclohexyl-1-(((S)-1-hydroxy-3-((S)-2-oxopyrrolidin-3-yl)propan-2-yl)amino)-1-oxopropan-2-yl)amino)-3-methyl-1-oxobutan-2-yl)carbamate (12c).**

To a solution of **6c** (250 mg, 0.44 mmol, 1.0 equiv) in anhydrous THF (10 mL) at 0 °C was added $LiBH_4$ (1.0 M in THF, 1.32 mL, 1.32 mmol, 3.0 equiv). The mixture was stirred at RT for 2 h. After the reaction was completed, excess reactants were consumed by slow addition of H2O. The mixture was diluted with $H_2O$ and extracted with EtOAc, washed with sat. NaCl, dried over $Na_2SO_4$ and concentrated. The residue was purified by column chromatography (MeOH: DCM = 1:15 v/v) to afford the pure product **12c** as a white solid (130 mg, 54%). $^1$H NMR (400 MHz, $CD_3OD$-$d_4$) δ 7.3 – 7.1 (m, 5H), 5.1 – 4.9 (m, 2H), 4.3 (dd, $J$ = 9.0, 6.4 Hz, 1H), 3.9 – 3.8 (m, 2H), 3.5 (q, $J$ = 7.1 Hz, 1H), 3.5 – 3.3 (m, 2H), 3.2 – 3.1 (m, 2H), 2.4 – 2.3 (m, 1H), 2.2 (s, 1H), 2.0 – 1.8 (m, 2H), 1.7 – 1.4 (m, 9H), 1.1 (q, $J$ = 8.1, 7.1 Hz, 3H), 0.8 (dd, $J$ = 8.6, 6.7 Hz, 8H). $^{13}$C NMR (100 MHz, $CD_3OD$-$d$) δ 181.2, 173.5, 172.8, 157.4, 136.8, 128.1, 128.1, 127.6, 127.5, 127.5, 66.5, 64.2, 60.9, 51.4, 49.1, 40.1, 38.9, 38.1, 33.9, 33.5, 32.3, 32.0, 30.5, 27.6, 26.2, 26.0, 25.8, 18.4, 17.0.

**Benzyl ((S)-1-(((S)-3-cyclohexyl-1-oxo-1-(((S)-1-oxo-3-((S)-2-oxopyrrolidin-3-yl)propan-2-yl)amino)propan-2-yl)amino)-3-methyl-1-oxobutan-2-yl)carbamate (MPI5).**

To a solution of **12c**(130 mg, 0.24 mmol, 1.0 equiv) in anhydrous DCM (10 mL) was added Dess-Martin reagent (200 mg, 0.48 mmol,2.0 equiv) slowly at 0 °C. Then the reaction mixture was stirred at RT for 1 h. A solution of $NaHCO_3$ and $Na_2S_2O_3$ was added to quench the reaction. After 10 min, the mixture was washed with water, sat. NaCl, dried over $Na_2SO_4$ and concentrated. The residue was purified by column chromatography (MeOH: DCM = 1:15 v/v). Dissolved the obtained aldehyde in abundant $CCl_4$ and hexane, then concentrate in vacuo. Re-dissolved the residue in least amount of $CHCl_3$ and add abundant hexane to precipitate out a white solid **MPI5** (60 mg, 47%).$^1$H NMR (400 MHz, $CDCl_3$-$d$) δ 9.4 (s, 1H), 8.1 (d, $J$ = 6.9 Hz, 1H), 7.3 (s, 5H), 7.1 (d, $J$ = 8.5 Hz, 1H), 6.7 (s, 1H), 5.6 (d, $J$ = 8.8 Hz, 1H), 5.0 (q, $J$ = 12.3 Hz, 2H), 4.6 (td, $J$ = 8.9, 5.7 Hz, 1H), 4.3 (p, $J$ = 5.0 Hz, 1H), 4.0 (t, $J$ = 7.8 Hz, 1H), 3.3 – 3.1 (m, 2H), 2.4 – 2.2 (m, 2H), 2.1 – 1.9 (m, 2H), 1.8 (ddd, $J$ = 13.6, 7.3, 4.1 Hz, 1H), 1.7 – 1.4 (m, 8H), 1.3 – 1.2 (m, 1H), 1.1 – 1.0 (m, 3H), 0.8 (dd, $J$ = 13.4, 6.9 Hz, 8H). $^{13}$C NMR (100 MHz, $CDCl_3$-$d$) δ 199.6, 180.1, 173.5, 171.5, 156.7, 136.3, 128.7, 128.7, 128.3, 128.1, 128.1, 67.2, 60.7, 57.4, 51.2, 40.7, 40.3, 38.0, 34.3, 33.6, 32.6, 31.1, 30.0, 28.4, 26.5, 26.3, 26.2, 19.3, 19.3. ESI-MS calcd for $C_{29}H_{43}N_4O_6^+$ (M+H$^+$): 543.3; found 543.3.

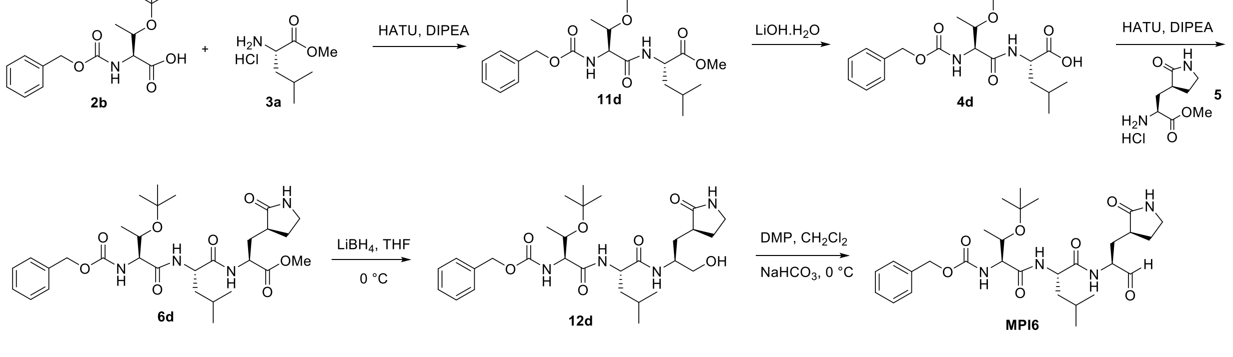


**Scheme S9.** The synthesis of compound **MPI6**.

**(S)-Methyl 2-((2S,3R)-2-(((benzyloxy)carbonyl)amino)-3-(tert-butoxy)butanamido)-4-methylpentanoate (11d)**

The amino acid methyl ester hydrochloride **3a** (1.0 g, 5.52 mmol) and the Cbz-protected amino acid **2b** (1.52 g, 6.08 mmol) were dissolved in dry DMF (20 mL) and the reaction was cooled to 0 °C. HATU (2.52 g, 6.62 mmol) and DIPEA (3.92 mL, 22.08 mmol) were added, and the reaction mixture was allowed warm up to room temperature and stirred for 12 h. The mixture was then poured into water (50 mL) and extracted with ethyl acetate (4×20 mL). The organic layer was washed with aqueous hydrochloric acid 10% v/v (2×20 mL), saturated aqueous $NaHCO_3$ (2×20 mL), brine (2×20 mL) and dried over $Na_2SO_4$. The organic phase was evaporated to dryness and the crude material purified by silica gel column chromatography (15-50% EtOAc in n-hexane as the eluent) to afford **11d** gummy liquid (1.71 g, 71%). $^1H$ NMR ($CDCl_3$, 400 MHz): δ 7.58 (d, *J*=7.64 Hz, 1H), 7.28-7.21 (m, 5H), 5.88 (d, *J*=4.68 Hz, 1H), 5.04 (ABq, *J*=12.16 Hz, 2H), 4.47-4.41 (m, 1H), 4.15-4.08 (m, 2H), 3.65 (s, 3H), 1.63-1.46 (m, 3H), 1.23 (s, 9H), 1.03 (d, *J*=6.28, 3H), 0.86 (dd, *J*=3.84, 5.92 Hz, 6H); $^{13}C$ NMR ($CDCl_3$, 100 MHz) δ 172.9, 169.4, 156.1, 136.3, 128.5 (2C), 128.0, 127.9, 75.5, 66.8, 60.4, 58.4, 52.2, 51.1, 41.3, 28.2 (3C), 25.0 22.8, 21.9, 16.4.

**(S)-2-((2S,3R)-2-(((Benzyloxy)carbonyl)amino)-3-(tert-butoxy)butanamido)-4-methylpentanoic acid (4d)**

The peptide **11d** (500 mg, 1.32 mmol) was dissolved in THF/$H_2O$ (1:1, 6.0 mL), and LiOH (138 mg, 3.30 mmol) was added at 0 °C. The mixture was stirred at room temperature overnight. Then THF was removed *on vacuum* and the aqueous layer was acidified with 1 M HCl and extracted with dichloromethane (3 x 10 mL). The organic layer was dried over anhydrous $Na_2SO_4$ and concentrated to yield **4d** as white solid (350 mg, 70%). $^1H$ NMR ($CDCl_3$, 400 MHz): δ 7.62 (d, *J*= 7.44 Hz, 1H), 7.29-7.22 (m, 5H), 5.94 (d, *J*=5.28 Hz, 1H), 5.04 (ABq, *J*=12.32 Hz, 2H), 4.45-4.40 (m, 1H), 4.17-4.08 (m, 2H), 1.68-1.50 (m, 3H), 1.21 (s, 9H), 1.02 (d, *J*=6.32 Hz, 3H), 0.87 (m, 6H); $^{13}C$ NMR ($CDCl_3$, 100 MHz) δ 177.1, 169.9, 156.2, 136.2, 128.5 (2C), 128.2, 128.0 (2C), 75.6, 66.97, 66.92, 58.4, 51.1, 41.0, 28.2 (3C), 25.0, 22.8, 21.8, 16.5.

**(5S,8S,11S)-Methyl 5-((R)-1-(tert-butoxy)ethyl)-8-isobutyl-3,6,9-trioxo-11-(((S)-2-oxopyrrolidin-3-yl)methyl)-1-phenyl-2-oxa-4,7,10-triazadodecan-12-oate (6d)**

The methyl (S)-2-amino-3-((S)-2-oxopyrrolidin-3-yl)propanoate hydrochloride **5** (140 mg, 0.606 mmol) and the peptide **4d** (242 mg, 0.666 mmol) were dissolved in dry DMF (6 mL), and the reaction was cooled to 0 °C. HATU (276 mg, 0.727 mmol) and DIPEA (0.43 mL, 2.42 mmol) were added, and the reaction mixture was allowed warm up to room temperature and stirred for 12 h. The mixture was then poured into water (20 mL) and extracted with ethyl acetate (4×20 mL). The organic layer was washed with aqueous hydrochloric acid 10% v/v (2×20 mL), saturated aqueous $NaHCO_3$ (2×20 mL), brine (2×20 mL) and dried over $Na_2SO_4$. The organic phase was evaporated to dryness and the crude material purified by silica gel column chromatography afford **6d** as white solid (240 mg, 64%). $^1H$ NMR ($CDCl_3$, 400 MHz): δ 7.64 (d, *J*=7.08 Hz, 1H), 7.39 (d, *J*=7.92 Hz, 1H), 7.31-7.20 (m, 5H), 6.34 (Brs, 1H), 5.87 (d, *J*=5.0 Hz, 1H), 5.03 (ABq, *J*=12.36 Hz, 2H), 4.50-4.43 (m, 1H), 4.50-4.43 (m, 1H), 4.40-4.34 (m, 1H), 4.11-4.09 (m, 2H), 3.67 (s, 3H), 3.29-3.15 (m, 2H), 2.38-2.23 (m, 2H), 2.10-2.02 (m, 1H), 1.81-1.61 (m, 4H), 1.51-1.44 (m, 1H),

1.18 (s, 9H), 0.99 (d, *J*=5.76, 3H), 0.86 (dd, *J*=5.6, 14.32, 6H).$^{13}$C NMR ($CDCl_3$, 100 MHz): δ 179.7, 172.2, 169.5, 136.2, 128.6 (2C), 128.3, 128.1 (2C), 77.1, 75.4, 67.0, 66.7, 58.9, 52.4, 51.9, 51.1, 41.7, 40.5, 38.2, 33.0, 28.2 (3C), 28.2, 24.7, 22.8, 22.2, 17.2.

**Benzyl ((2S,3R)-3-(tert-butoxy)-1-(((S)-1-(((S)-1-hydroxy-3-((S)-2-oxopyrrolidin-3-yl)propan-2-yl)amino)-4-methyl-1-oxopentan-2-yl)amino)-1-oxobutan-2-yl)carbamate (12d)**

To a stirring solution of compound **6d** (120mg, 0.225 mmol) in THF (5 mL) was added $LiBH_4$ (2.0 M in THF, 0.56 mL, 1.12 mmol) in several portions at 0 °C under a nitrogen atmosphere. The reaction mixture was stirred at 0 °C for 1 h, then allowed to warm up to room temperature, and stirred for an additional 2 h. The reaction was quenched by the drop wise addition of 1.0 M HCl(aq) (1.2 mL) with cooling in an ice bath. The solution was diluted with ethyl acetate and $H_2O$. The phases were separated, and the aqueous layer was extracted with ethyl acetate (3×15 mL). The organic phases were combined together, dried over $Na_2SO_4$, filtered, and concentrated on a rotorvap to give a yellow oily residue. Column chromatographic purification of the residue (6% MeOH in $CH_2Cl_2$ as the eluent) afforded a white solid **12d** (85 mg, 68%).$^{1}$H NMR ($CDCl_3$, 400 MHz) δ 7.66 (d, *J*=7.48 Hz, 1H), 7.41-7.30 (m, 5H), 6.13 (Brs, 1H), 6.02 (Brs, 1H), 5.13 (ABq, *J*=12.2 Hz, 2H), 4.42-4.36 (m, 1H), 4.18 (d, *J*=5.04, 2H), 4.07-3.97 (m, 1H), 3.65-3.58 (m, 2H), 3.35-3.28 (m, 2H), 2.46-2.37 (m, 2H), 2.10-2.02 (m, 1H), 1.97-1.78 (m, 2H), 1.71-1.53 (m, 3H), 1.28 (s, 9H), 1.09 (d, *J*=5.64 Hz, 3H), 0.94 (d, *J*=6.12, 10.36 Hz, 6H); $^{13}$C NMR ($CDCl_3$, 100 MHz): δ 180.9, 172.7, 169.8, 156.3, 136.1, 128.6 (2C), 77.2, 75.5, 66.9, 65.9, 59.1, 55.2, 50.5, 41.2, 40.5, 38.2, 32.4, 28.5, 28.2, 24.9, 22.8, 22.1, 17.4.

**Benzyl ((2S,3R)-3-(tert-butoxy)-1-(((S)-4-methyl-1-oxo-1-(((S)-1-oxo-3-((S)-2-oxopyrrolidin-3-yl)propan-2-yl)amino)pentan-2-yl)amino)-1-oxobutan-2-yl)carbamate (MPI6)**

To a solution of **12d** (70 mg, 0.138 mmol) in $CH_2Cl_2$ (6 mL) was added $NaHCO_3$ (46 mg, 4 equiv) and the Dess-Martin reagent (180 mg, 3 equiv). The resulting mixture was stirred at rt for 12 h. Then the reaction was quenched with a saturated $NaHCO_3$ solution containing 10 % $Na_2S_2O_3$. The layers were separated. The organic layer was then washed with saturated brine solution, dried over anhydrous $Na_2SO_4$ and concentrated *on vacuum*. The residue was then purified with flash chromatography afford **MPI6** as white solid (41 mg, 59%). $^{1}$H NMR (400 MHz, $CDCl_3$): δ 9.44 (s, 1H), 8.04 (d, J=6.24 Hz, 1H), 7.41 (d, *J*= 7.64 Hz, 1H), 7.31-7.27 (m, 5H), 6.14 (brs, 1H), 5.85 (d, *J*= 4.68 Hz, 1H), 5.05 (ABq, *J*= 12.2 Hz, 2H), 4.45-4.38 (m, 1H), 4.34-4.28 (m, 1H), 4.18-4.05 (m, 2H), 3.30-3.11 (m, 2H), 2.48-2.33 (m, 1H), 2.33-2.21 (m, 1H), 2.11-1.92 (m, 2H), 1.91-1.89 (m, 1H), 1.87-1.58 (m, 1H), 1.57-1.53 (m, 2H), 1.52-1.48 (m, 1H), 1.21 (s, 9H), 1.01 (d, *J*= 5.92 Hz, 3H), 0.89 (dd, *J*= 6.12, 11.56 Hz, 6H). $^{13}$C NMR ($CDCl_3$, 100 MHz): 199.6, 179.9, 172.8, 169.7, 156.2, 136.1, 128.6 (2C), 128.3, 128.1 (2C), 77.1, 75.4, 67.0, 66.7, 58.9, 57.6, 52.1, 41.6, 40.5, 37.9, 29.7, 28.5, 28.2 (3C), 24.9, 22.9, 22.1, 17.3.

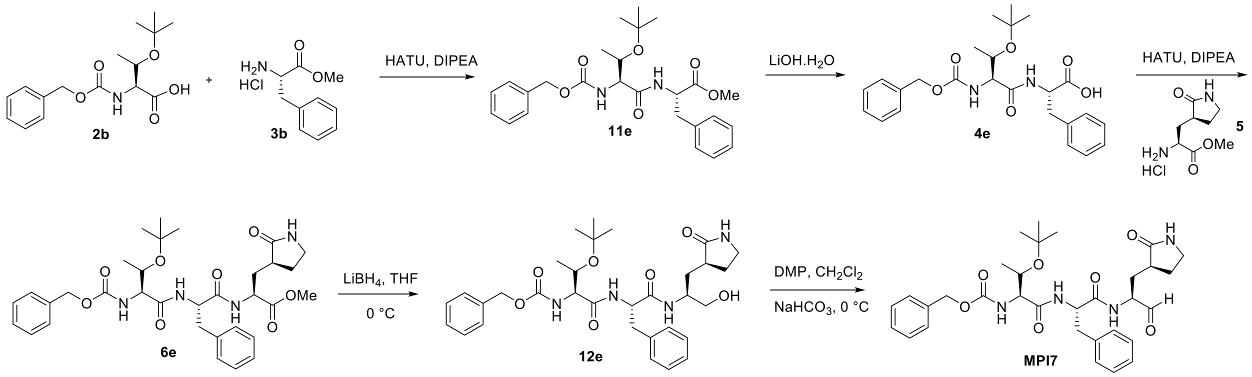


**Scheme S10.** The synthesis of compound **MPI7**.

#### Methyl N-((benzyloxy)carbonyl)-O-(tert-butyl)-L-allothreonyl-L-phenylalaninate (11e)

To a solution of **2b** (5 mmol, 1.55 g) and **3b** (5 mmol, 1.07g) in anhydrous DMF (20 mL) was added DIPEA (10 mmol, 1.29 g) and was cooled to 0 °C. HATU (5.5 mmol, 2.09 g) was added to the solution under 0 °C and then stirred at room temperature overnight. The reaction mixture was then diluted with ethyl acetate (100 mL) and washed with saturated $NaHCO_3$ solution (2×50 mL), 1 M HCl solution (2×50 mL), and saturated brine solution (2×50 mL) sequentially. The organic layer was dried over anhydrous $Na_2SO_4$ and then concentrated *on vacuo*. The residue was then purified with flash chromatography (15-50% EtOAc in hexanes as the eluent) to afford **11e** as colorless oil (1.64 g, 70%).

#### N-((benzyloxy)carbonyl)-O-(tert-butyl)-L-allothreonyl-L-phenylalanine (4e)

**11e** (1 mmol, 470 mg) was dissolved in 5 mL of THF. A solution of LiOH·$H_2O$ (2 mmol, 84 mg) in 5 mL $H_2O$ was added to the solution. The mixture was stirred at room temperature overnight. Then THF was removed *on vacuo* and the aqueous layer was acidified with 1 M HCl and extracted with dichloromethane (3×10 mL). The organic layer was dried over anhydrous $Na_2SO_4$ and concentrated to yield **4e** as white solid (330 mg, 72 %). $^1$H-NMR (400 MHz, $CD_3OD$): δ 7.29-7.43 (m, 5H), 7.11-7.29 (m, 5H), 5.11 (s, 2H), 4.71 (t, J = 6.2 Hz, 1H), 4.14 (d, J = 3.9 Hz, 1H), 3.98-4.07 (m, 1H), 3.17 (dd, J = 13.9, 5.6 Hz, 1H), 3.05 (dd, J = 13.8, 6.8 Hz, 1H), 0.98-1.16 (m, 12H).

#### Methyl (5S,8S,11S)-8-benzyl-5-((S)-1-(tert-butoxy)ethyl)-3,6,9-trioxo-11-(((S)-2-oxopyrrolidin-3-yl)methyl)-1-phenyl-2-oxa-4,7,10-triazadodecan-12-oate (6e)

To a solution of **4e** (0.4 mmol, 182 mg) and **5** (0.4 mmol, 88 mg) in anhydrous DMF (2 mL) was added DIPEA (0.8 mmol, 103 mg) and was cooled to 0 °C. HATU (0.44 mmol, 167 mg) was added to the solution under 0 °C and then stirred at room temperature overnight. The reaction mixture was then diluted with ethyl acetate (20 mL) and washed with saturated $NaHCO_3$ solution (2×10 mL), 1 M HCl solution (2×10 mL), and saturated brine solution (2×10 mL) sequentially. The organic layer was dried over anhydrous $Na_2SO_4$ and then concentrated *on vacuo*. The residue was then purified with flash chromatography (1-10% methanol in dichloromethane as the eluent) to afford **6b** as white solid (180 mg, 72%). $^1$H-NMR (400 MHz, $CDCl_3$): δ 7.44 – 7.19 (m, 10H), 5.90 (d, J = 5.4 Hz, 1H), 5.52 (s, 1H), 5.12 (q, J = 12.3 Hz, 2H), 4.77 (q, J = 6.8 Hz, 1H), 4.65 –

4.53 (m, 1H), 4.22 – 4.09 (m, 2H), 3.72 (s, 3H), 3.40 – 3.29 (m, 2H), 3.21 (dd, J = 14.0, 6.4 Hz, 1H), 3.09 (dd, J = 13.9, 6.1 Hz, 1H), 2.50 – 2.40 (m, 1H), 2.35 – 2.26 (m, 1H), 2.18 – 2.05 (m, 1H), 1.95 – 1.77 (m, 2H), 1.29 (s, 2H), 1.17 (s, 9H), 1.08 (d, J = 6.2 Hz, 3H).

**Benzyl ((2S,3S)-3-(tert-butoxy)-1-oxo-1-(((S)-1-oxo-1-(((S)-1-oxo-3-((S)-2-oxopyrrolidin-3-yl)propan-2-yl)amino)-3-phenylpropan-2-yl)amino)butan-2-yl)carbamate (MPI7)**

To a solution of **6e** (0.1 mmol, 62 mg) in anhydrous dichloromethane (5 mL) was added a solution of $LiBH_4$ in anhydrous THF (2 M, 0.1 mL, 0.2 mmol) at 0 °C. The resulting solution was stirred at the same temperature for 3 h. Then a saturated solution of $NH_4Cl$ (5 mL) was added dropwise to quench the reaction. The layers were separated, and the organic layer was washed with saturated brine solution (2×10 mL), dried over anhydrous $Na_2SO_4$. and evaporated to dryness. The residue was then dissolved in anhydrous dichloromethane (5 mL) and cooled to 0 °C. Dess-Martin periodinane (0.2 mmol, 85 mg) was added to the solution. The reaction mixture was then stirred at room temperature overnight. Then the reaction was quenched with a saturated $NaHCO_3$ solution containing 10 % $Na_2S_2O_3$. The layers were separated. The organic layer was then washed with saturated brine solution (2×10 mL), dried over anhydrous $Na_2SO_4$ and evaporated *on vacuo*. The residue was then purified with flash chromatography (1-10% methanol in dichloromethane as the eluent) to afford **MPI7** as white solid (27 mg, 45 %). $^1$H-NMR (400MHz, $CDCl_3$): δ 9.28 (s, 1H), 7.80 (d, J = 6.5 Hz, 1H), 7.44 – 7.09 (m, 10H), 5.87 (d, J = 5.6 Hz, 1H), 5.71 (s, 1H), 5.08 (q, J = 11.1 Hz, 2H), 4.76 (q, J = 7.1 Hz, 1H), 4.32 (q, J = 7.0 Hz, 1H), 4.19 – 4.04 (m, 2H), 3.38 – 3.26 (m, 2H), 3.19 (dd, J = 13.8, 6.3 Hz, 1H), 3.07 (dd, J = 13.8, 6.8 Hz, 1H), 2.41 – 2.22 (m, 2H), 1.88 (t, J = 6.7 Hz, 2H), 1.84 – 1.73 (m, 1H), 1.30 – 1.22 (m, 1H), 1.16 (s, 9H), 1.05 (d, J = 6.1 Hz, 3H); $^{13}$C NMR (100 MHz, $CDCl_3$) δ 199.60, 179.68, 171.55, 169.49, 156.32, 136.34, 136.28, 129.49, 128.87, 128.73, 128.40, 128.20, 127.35, 84.26, 75.61, 67.15, 66.80, 59.18, 57.74, 54.53, 40.53, 38.32, 37.82, 28.80, 28.23, 17.63. ESI-MS calcd for $C_{32}H_{43}N_4O_7$ (M+H$^+$): 595.3; found: 595.4.

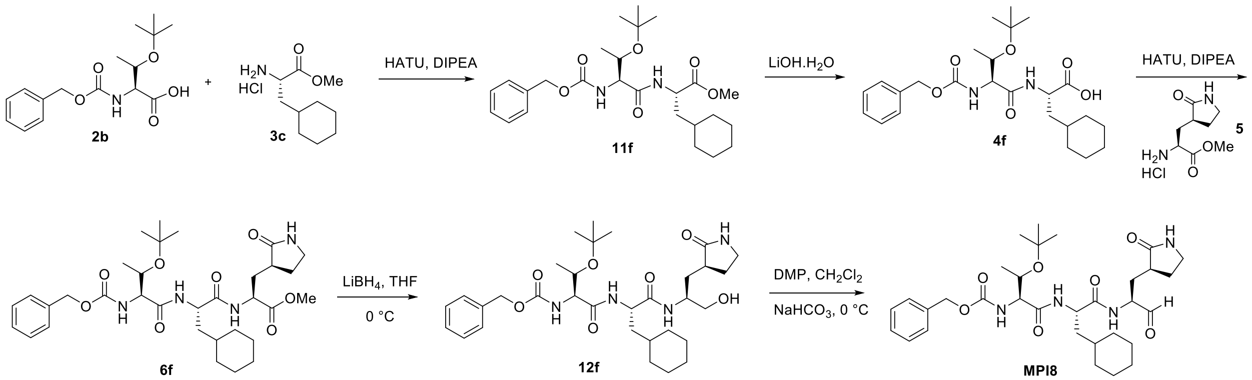


**Scheme S11.** The synthesis of compound **MPI8**.

**Methyl (S)-2-((2S,3R)-2-(((benzyloxy)carbonyl)amino)-3-(tert-butoxy)butanamido)-3-cyclohexylpropanoate (11f).**

To a solution of **2b** (2 g, 6.46 mmol, 1.0 equiv), in anhydrous DMF (15 mL) at 0°C, and then **3c** (1.4 g, 6.46 mmol, 1.0 equiv), HATU (3.7 g, 9.70 mmol, 1.5 equiv) and DIPEA (5.7 mL, 32.3 mmol, 5.0 equiv) was added sequentially. The mixture was stirred at RT for 6 h. The mixture was diluted with EtOAc and washed with water, 1M HCl, sat. NaCl, dried over $Na_2SO_4$, and concentrated. The residue was purified by column chromatography (EtOAc: Hexane = 1:4 v/v) to afford the pure product **11f** as a colorless oil (2.5 g, 83%). $^1$H NMR (400 MHz, $CDCl_3$-*d*) δ 7.6 (d, *J* = 7.8 Hz, 1H), 7.3 – 7.1 (m, 5H), 5.9 (d, *J* = 5.3 Hz, 1H), 5.1 – 4.9 (m, 2H), 4.4 (td, *J* = 8.4, 5.1 Hz, 1H), 4.2 – 4.1 (m, 2H), 3.6 (s, 3H), 1.7 – 1.5 (m, 6H), 1.5 (ddd, *J* = 14.2, 9.0, 5.7 Hz, 1H), 1.2 (s, 9H), 1.2 – 1.1 (m, 3H), 1.0 (d, *J* = 6.3 Hz, 4H), 0.9 – 0.7 (m, 2H). $^{13}$C NMR (100 MHz, $CDCl_3$-*d*) δ 172.7, 169.2, 155.9, 136.2, 128.3, 128.3, 127.8, 127.7, 127.7, 75.2, 66.7, 66.5, 60.0, 51.8, 50.2, 39.5, 34.0, 33.3, 32.3, 28.0, 28.0, 28.0, 26.1, 26.0, 25.8, 20.7.

**(S)-2-((2S,3R)-2-(((benzyloxy)carbonyl)amino)-3-(tert-butoxy)butanamido)-3-cyclohexylpropanoic acid (4f).**

To a solution of **11f** (400 mg, 0.84 mmol, 1.0 equiv) in 1:1 THF/$H_2O$ (8 mL) was added $LiOH\cdot H_2O$ (140 mg, 3.4 mmol, 4.0 equiv). The reaction was stirred at RT for 2 h. After completion, the reaction mixture was neutralized with 1M HCl solution and extracted with EtOAc. The organic layer was washed with sat. NaCl, dried over $Na_2SO_4$ and concentrated to afford the product **4f** (330 mg, 85%) as a white solid. The residue was used in the next without further purification.

**Methyl (5S,8S,11S)-5-((S)-1-(tert-butoxy)ethyl)-8-(cyclohexylmethyl)-3,6,9-trioxo-11-(((S)-2-oxopyrrolidin-3-yl)methyl)-1-phenyl-2-oxa-4,7,10-triazadodecan-12-oate (6f).**

To a solution of **4f** (300 mg, 0.65 mmol, 1.0 equiv) in anhydrous DMF (5 mL) at 0°C, and then **5** (144 mg, 0.65 mmol, 1.0 equiv), HATU (370 mg, 0.98 mmol, 1.5 equiv), DIPEA (580μL, 3.25 mmol, 5.0 equiv) was added sequentially. The mixture was stirred at RT for 6 h. The mixture was diluted with EtOAc and washed with water, 1M HCl, sat. NaCl, dried over $Na_2SO_4$, and concentrated. The residue was purified by column chromatography (MeOH: DCM = 1:15 v/v) to afford the pure product **6f** as a white solid (265 mg, 65%).

**Benzyl ((2S,3S)-3-(tert-butoxy)-1-(((S)-3-cyclohexyl-1-(((S)-1-hydroxy-3-((S)-2-oxopyrrolidin-3-yl)propan-2-yl)amino)-1-oxopropan-2-yl)amino)-1-oxobutan-2-yl)carbamate (12f).**

To a solution of **6f** (250 mg, 0.40 mmol, 1.0 equiv) in anhydrous THF (10 mL) at 0 °C was added $LiBH_4$ (1.0 M in THF, 1.2 mL, 1.20 mmol, 3.0 equiv). The mixture was stirred at RT for 2 h. After the reaction was completed, excess reactants were consumed by slow addition of $H_2O$. The mixture was diluted with $H_2O$ and extracted with EtOAc, washed with sat. NaCl, dried over $Na_2SO_4$ and concentrated. The residue was purified by column chromatography (MeOH: DCM = 1:15 v/v) to afford the pure product **12f**as a white solid (160 mg, 66%).

**Benzyl ((2S,3S)-3-(tert-butoxy)-1-(((S)-3-cyclohexyl-1-oxo-1-(((S)-1-oxo-3-((S)-2-oxopyrrolidin-3-yl)propan-2-yl)amino)propan-2-yl)amino)-1-oxobutan-2-yl)carbamate (MPI8).**

To a solution of **12f** (160 mg, 0.26 mmol, 1.0 equiv) in anhydrous DCM (10 mL) was added Dess-Martin reagent (225 mg, 0.52 mmol,2.0 equiv) slowly at 0 °C. Then the reaction mixture was stirred at RT for 1 h. A solution of $NaHCO_3$ and $Na_2S_2O_3$ was added to quench the reaction. After 10 min, the mixture was washed with water, sat. NaCl, dried over $Na_2SO_4$ and concentrated. The residue was purified by column chromatography (MeOH: DCM = 1:15 v/v). Dissolved the obtained aldehyde in abundant $CCl_4$ and hexane, then concentrate in vacuo. Re-dissolved the residue in least amount of $CHCl_3$ and add abundant hexane to precipitate out a white solid **MPI8** (82 mg, 53%).$^1$H NMR (400 MHz, $CDCl_3$-*d*) δ 9.5 (s, 1H), 8.1 (d, *J* = 6.8 Hz, 1H), 7.5 (q, *J* = 8.4, 7.6 Hz, 1H), 7.4 – 7.3 (m, 5H), 6.8 (s, 1H), 5.9 (d, *J* = 5.6 Hz, 1H), 5.1 (q, *J* = 12.1 Hz, 2H), 4.5 (td, *J* = 8.6, 5.6 Hz, 1H), 4.4 (q, *J* = 9.9, 7.2 Hz, 1H), 4.2 – 4.1 (m, 2H), 3.2 (p, *J* = 8.4, 7.1 Hz, 2H), 2.5 – 2.4 (m, 1H), 2.3 – 2.2 (m, 1H), 2.0 (ddt, *J* = 16.2, 11.3, 5.6 Hz, 1H), 1.8 (ddd, *J* = 13.3, 7.9, 4.2 Hz, 1H), 1.8 – 1.5 (m, 8H), 1.4 – 1.1 (m, 12H), 1.0 (d, *J* = 6.2 Hz, 4H), 1.0 – 0.8 (m, 2H). $^{13}$C NMR (100 MHz, $CDCl_3$-*d*) δ 199.7*,* 180.0*,* 173.0*,* 169.8*,* 156.3*,* 136.2*,* 128.6, 128.6, 128.3*,* 128.1, 128.1, 75.4*,* 67.0*,* 66.8*,* 59.0*,* 57.3*,* 51.4*,* 40.6*,* 40.2*,* 37.9*,* 34.2, 34.2, 33.6*,* 32.7*,* 29.9*,* 28.3, 28.3, 28.3, 26.4*,* 26.2*,* 26.0*,* 17.4. ESI-MS calcd for $C_{32}H_{49}N_4O_7^+$ (M+H$^+$): 601.3; found 601.3.

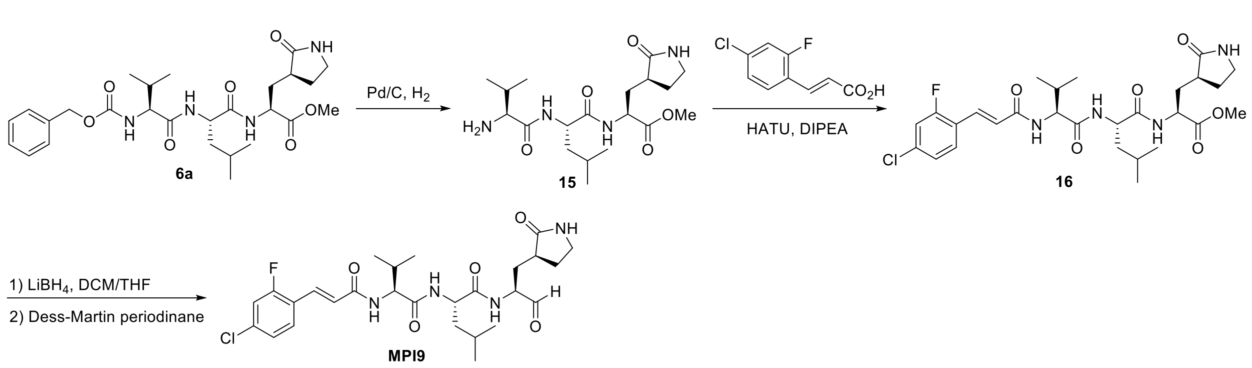


**Scheme S12**. The synthesis of compound **MPI9**.

#### Methyl (S)-2-((S)-2-((S)-2-((E)-3-(4-chloro-2-fluorophenyl)acrylamido)-3-methylbutanamido)-4-methylpentanamido)-3-((S)-2-oxopyrrolidin-3-yl)propanoate (16)

To a solution of **6a** (106 mg, 0.2 mmol) in methanol (5 mL) was added 10% Pd/C (21 mg). The mixture was then stirred with hydrogen balloon at room temperature for 3 h. The reaction mixture was then filtered, and the filtrate was concentrated *in vacuo* to afford **15** as colorless oil, which was used without further purification.

To a solution of **15** in 2 mL anhydrous DMF was added 2-fluoro-4-chlorocinnamic acid (40 mg, 0.2 mmol) and DIPEA (0.4 mmol, 52 mg). The solution was cooled to 0 °C, follow by the addition of HATU (0.24 mmol, 91 mg). The solution was then allowed to warm up to room temperature and stirred overnight. The reaction mixture was then diluted with ethyl acetate (20 mL) and washed with saturated $NaHCO_3$ solution (2×10 mL), 1 M HCl solution (2×10 mL), and saturated brine solution (2×10 mL) sequentially. The organic layer was dried over anhydrous $Na_2SO_4$ and then concentrated *in vacuo*. The residue was then purified with flash chromatography (1-10% methanol in dichloromethane as the eluent) to afford **16** as white solid (75 mg, 65%). $^1$H NMR (400 MHz, DMSO-$d_6$) δ 8.46 (d, J = 8.2 Hz, 1H), 8.25 (d, J = 8.9 Hz, 1H), 8.11 (d, J = 7.7 Hz, 1H), 7.66 (dd, J = 17.7, 9.4 Hz, 2H), 7.54 (dd, J = 10.7, 2.1 Hz, 1H), 7.44 (d, J = 16.0 Hz, 1H),

7.38 (dd, J = 8.4, 2.1 Hz, 1H), 7.00 (d, J = 16.0 Hz, 1H), 4.42 – 4.24 (m, 3H), 3.62 (s, 3H), 3.16 (t, J = 9.2 Hz, 1H), 3.06 (q, J = 9.3, 8.9 Hz, 1H), 2.36 – 2.27 (m, 1H), 2.12 – 1.95 (m, 3H), 1.62 (dtd, J = 20.8, 11.7, 10.8, 6.9 Hz, 3H), 1.46 (t, J = 7.3 Hz, 2H), 0.95 – 0.81 (m, 12H).

**(S)-2-((S)-2-((E)-3-(4-chloro-2-fluorophenyl)acrylamido)-3-methylbutanamido)-4-methyl-N-((S)-1-oxo-3-((S)-2-oxopyrrolidin-3-yl)propan-2-yl)pentanamide (MPI9)**

To a solution of **16** (0.1 mmol, 58 mg) in anhydrous tetrahydrofuran (5 mL) was dropwise added a solution of $LiBH_4$ in tetrahydrofuran (2 M, 0.1 mL, 0,2 mmol) at 0 °C. The solution was stirred under the same temperature for 3 h. Then a saturated solution of $NH_4Cl$ (5 mL) was added dropwise to quench the reaction. The layers were separated, and the organic layer was washed with saturated brine solution (2×10 mL), dried over anhydrous $Na_2SO_4$. and evaporated to dryness. The residue was then dissolved in anhydrous dichloromethane (5 mL) and cooled to 0 °C. Dess-Martin periodinane (0.2 mmol, 85 mg) was added to the solution. The reaction mixture was then stirred at room temperature overnight. Then the reaction was quenched with a saturated $NaHCO_3$ solution containing 10% $Na_2S_2O_3$. The layers were separated. The organic layer was then washed with saturated brine solution (2×10 mL), dried over anhydrous $Na_2SO_4$ and evaporated *in vacuo*. The residue was then purified with flash chromatography (1-10% methanol in dichloromethane as the eluent) to afford **MPI9** as white solid (32 mg, 58%). $^1$H NMR (400 MHz, DMSO-$d_6$) δ 9.41 (s, 1H), 8.46 (d, J = 7.9 Hz, 1H), 8.26 (d, J = 8.7 Hz, 1H), 8.17 (d, J = 7.6 Hz, 1H), 7.66 (t, J = 8.3 Hz, 1H), 7.62 (s, 1H), 7.53 (dd, J = 10.6, 2.1 Hz, 1H), 7.43 (d, J = 15.9 Hz, 1H), 7.37 (dd, J = 8.5, 2.0 Hz, 1H), 7.00 (d, J = 15.9 Hz, 1H), 4.42 – 4.18 (m, 3H), 3.16 (t, J = 9.1 Hz, 1H), 3.06 (td, J = 9.4, 7.1 Hz, 1H), 2.36 – 2.21 (m, 1H), 2.13 (dt, J = 14.0, 7.8 Hz, 1H), 2.01 (h, J = 6.6 Hz, 1H), 1.88 (ddd, J = 14.9, 11.4, 4.0 Hz, 1H), 1.70 – 1.56 (m, 3H), 1.55 – 1.39 (m, 2H), 0.97 – 0.73 (m, 12H); $^{13}$C NMR (101 MHz, DMSO) δ 201.24, 178.73, 173.08, 171.32, 165.00, 161.93, 159.41, 135.01, 134.90, 130.74, 130.63, 126.15, 125.84, 122.35, 122.23, 117.35, 117.09, 58.07, 56.54, 51.68, 41.08, 37.59, 31.34, 29.82, 27.75, 24.67, 23.30, 22.24, 19.59, 18.53. ESI-MS: calcd for $C_{27}H_{37}ClFN_4O_5$ ($M+H^+$): 551.2; found 551.4.

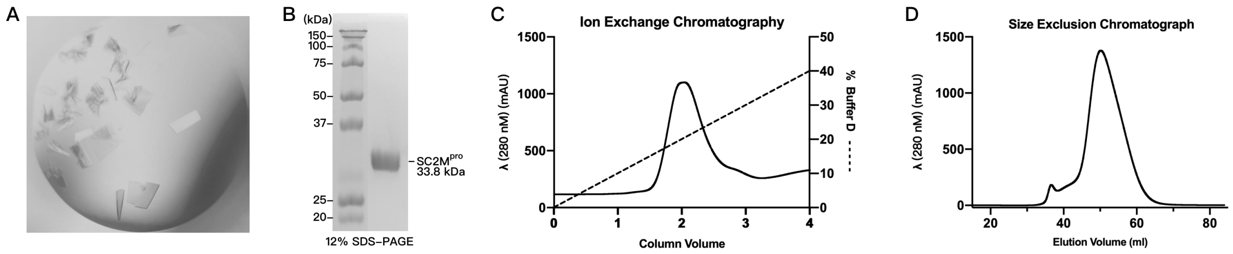


**Figure S1.** Purification and Crystallization of SC2M$^{pro}$. (A) Sitting-drop vapor-diffusion method was employed to produce plate-like crystals of SC2M$^{pro}$. (B) The purified SC2M$^{pro}$ protein was analyzed by reducing 12% SDS-PAGE with Coomassie blue staining. (C) and (D) Ion exchange and size exclusion chromatography were performed to purify the SC2M$^{pro}$ protein.

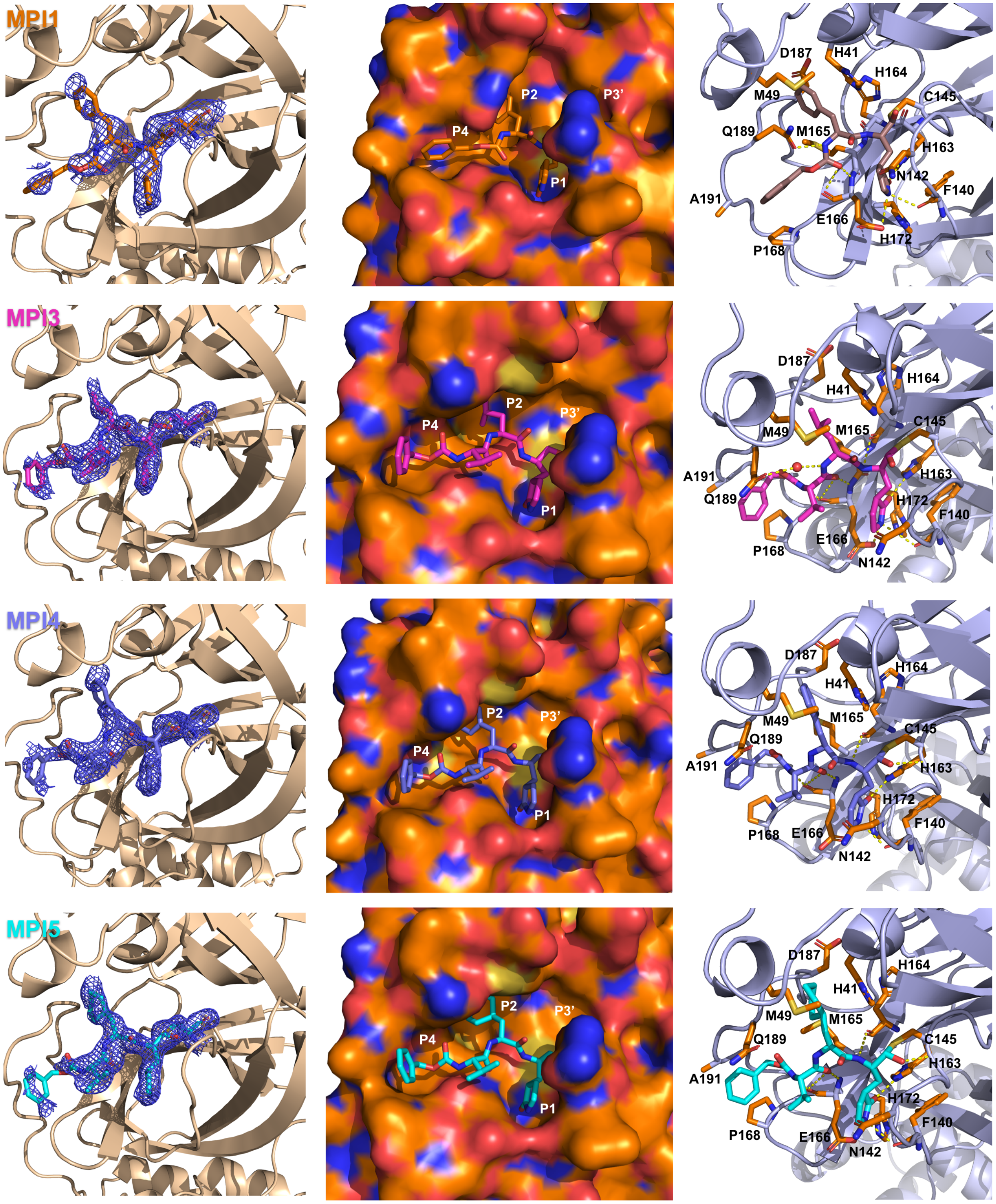
MPI1
P2
P3'
P4
P1
D187
H41
H164
M49
C145
Q189
M165
H163
N142
F140
A191
E166
H172
P168
MPI3
D187
H164
H41
P2
P3'
P4
M49
M165
C145
A191
H163
Q189
H172
F140
P1
E166
P168
N142
MPI4
D187
H164
H41
P2
P3'
M49
M165
C145
Q189
P4
A191
H163
H172
P1
F140
P168
E166
N142
MPI5
D187
H164
H41
P2
P3'
M49
M165
P4
Q189
C145
H163
A191
H172
P1
F140
P168
E166
N142

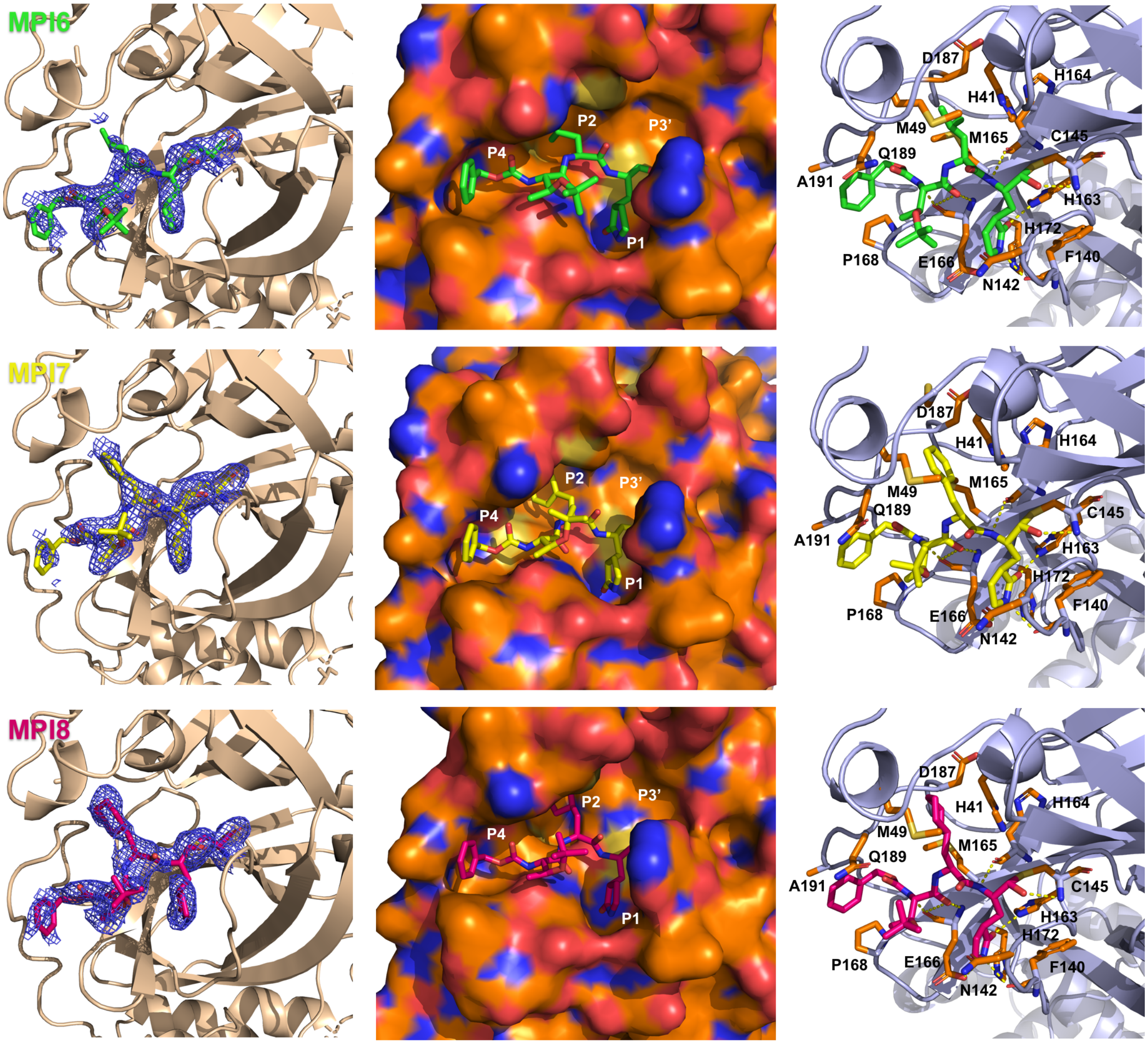


**Figure S2**. X-ray crystallography analysis of SC2M$^{Pro}$ in complexes with different inhibitors. A contoured 2Fo-Fc map at the 0.5σlevel around inhibitors and C145 are shown in the active site of SC2M$^{Pro}$ at left panel. The occupation of the active site cavity of SC2M$^{Pro}$ by MPI3 is shown in the middle panel. Extensive hydrogen bonding and Van der Waals interactions between SC2M$^{Pro}$ and inhibitors are shown in the right panel.

| | apo | MPI1 | MPI3 | MPI4 |
|---|---|---|---|---|
| Data Collection | | | | |
| Space group | C121 | C121 | C121 | C121 |
| cell dimensions | | | | |
| *a*, *b*, *c* (Å) | 114.89, 53.44, 44.88 | 114.12, 53.90, 45.42 | 114.63, 53.48, 45.38 | 114.26, 53.59, 45.34 |
| α, β, γ (°) | 90.00, 101.46, 90.00 | 90.00, 101.48, 90.00 | 90.00, 101.73, 90.00 | 90.00, 101.60, 90.00 |
| Resolution (Å) | 56.30-1.77 (2.05-2.00) | 48.56-2.20 (2.27-2.20) | 48.28-1.8 (1.84-1.80) | 48.34-1.80 (1.84-1.80) |
| $R_{merge}$ | 7.8 (87.1) | 10.9 (83.9) | 11.3 (81.8) | 8.9 (92.3) |
| $I/\sigma I$ | 9.9 (1.4) | 6.7 (2.0) | 6.5 (1.0) | 7.7 (1.2) |
| Completeness (%) | 95.2 (92.6) | 88.8 (80.6) | 94.8 (91.5) | 94.5 (91.3) |
| Redundancy | 3.6 (3.5) | 3.3 (3.0) | 3.7 (3.8) | 4.1 (4.1) |
| Refinement | | | | |
| Resolution (Å) | 31.74-2.00 | 48.56-2.20 | 48.28-1.80 | 48.34-1.80 |
| No. Reflections | 17216 (1673) | 12021 (917) | 23657 (2277) | 23533 (2247) |
| $R_{work}/R_{free}$ | 0.1734/0.2266 | 0.2145/0.3027 | 0.1769/0.2266 | 0.1964/0.2501 |
| No. atoms | | | | |
| Protein | 2367 | 2443 | 2438 | 2441 |
| Water | 135 | 116 | 232 | 188 |
| *B* factors | | | | |
| Protein | 35.465 | 56.617 | 33.154 | 43.068 |
| Water | 40.602 | 62.841 | 40.951 | 51.413 |
| R.m.s deviations | | | | |
| Bond lengths (Å) | 0.007 | 0.010 | 0.007 | 0.008 |
| Bond angles (°) | 0.88 | 1.19 | 1.07 | 1.11 |

| | MPI5 | MPI6 | MPI7 | MPI8 |
|---|---|---|---|---|
| Data Collection | | | | |
| Space group | C121 | C121 | C121 | C121 |
| cell dimensions | | | | |
| *a*, *b*, *c* (Å) | 114.67, 53.18, 45.22 | 114.08, 53.76, 45.24 | 115.07, 53.84, 45.43 | 114.83, 53.72, 45.39 |
| α, β, γ (°) | 90.00, 101.72, 90.00 | 90.00, 101.77, 90.00 | 90.00, 101.47, 90.00 | 90.00, 101.88, 90.00 |
| Resolution (Å) | 48.06-1.70 (1.73-1.70) | 48.44-2.10 (2.16-2.10) | 48.59-2.10 (2.16-2.10) | 48.47-1.90 (1.94-1.90) |
| $R_{merge}$ | 7.0 (38.2) | 7.7 (75.9) | 10.5 (86.3) | 10.2 (66.5) |
| $I/\sigma I$ | 13.3 (4.0) | 8.8 (1.4) | 6.7 (1.3) | 5.5 (2.0) |
| Completeness (%) | 93.4 (88.5) | 99.9 (99.1) | 99.1 (97.7) | 96.6 (91.1) |
| Redundancy | 4.0 (4.0) | 4.2 (3.9) | 3.6 (3.5) | 3.1 (2.6) |
| Refinement | | | | |
| Resolution (Å) | 48.06-1.70 | 48.44-2.10 | 44.52-2.10 | 48.47 - 1.90 |
| No. Reflections | 27392 (2625) | 15764 (1570) | 15881 (1560) | 19879 (1655) |
| $R_{work}/R_{free}$ | 0.1795/0.2239 | 0.1965/0.2501 | 0.1912/0.2412 | 0.2522/0.3364 |
| No. atoms | | | | |
| Protein | 2437 | 2442 | 2445 | 2445 |
| Water | 240 | 104 | 109 | 122 |
| *B* factors | | | | |
| Protein | 33.886 | 57.073 | 54.357 | 46.009 |
| Water | 41.660 | 63.681 | 61.346 | 51.370 |
| R.m.s deviations | | | | |
| Bond lengths (Å) | 0.007 | 0.008 | 0.008 | 0.008 |
| Bond angles (°) | 1.00 | 1.11 | 1.13 | 1.12 |

**Table S1.** Statistics of crystallographic analysis of $SC2M^{Pro}$ in its apo-form and complexed with different inhibitors.

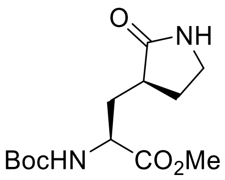

**14**

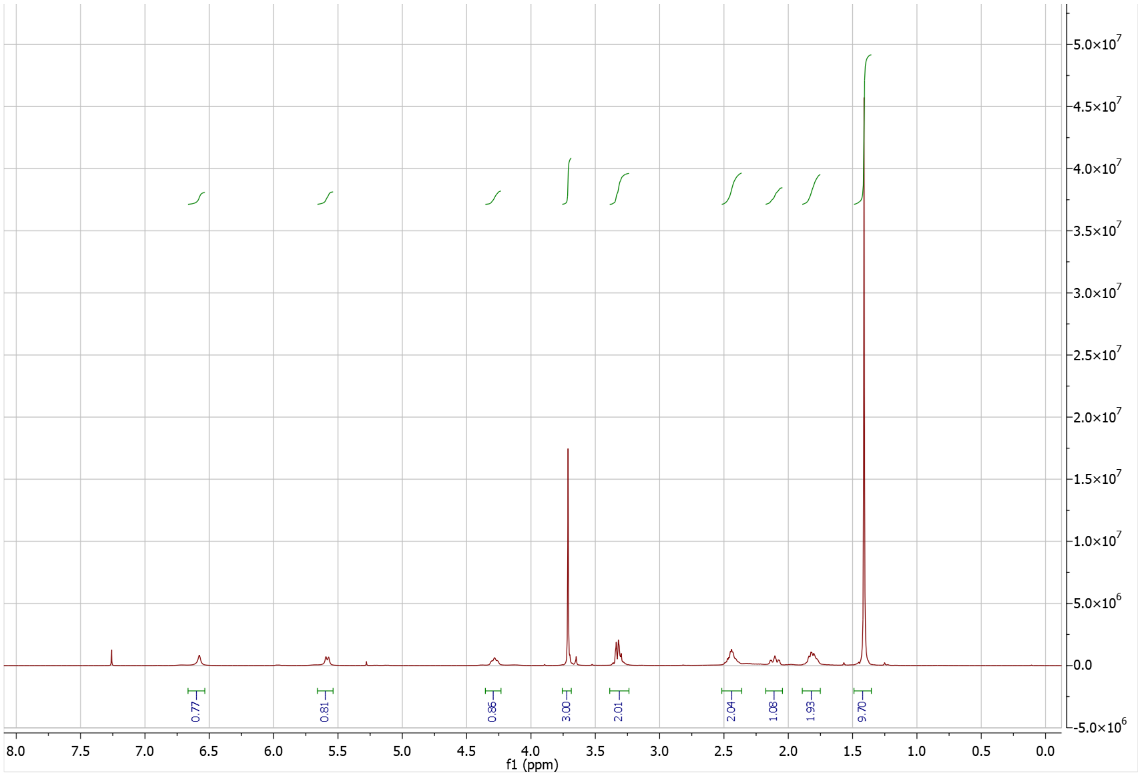

0.77
0.81
0.86
3.00
2.01
2.04
1.08
1.93
9.70
8.0 7.5 7.0 6.5 6.0 5.5 5.0 4.5 4.0 3.5 3.0 2.5 2.0 1.5 1.0 0.5 0.0
f1 (ppm)
5.0×10^7
4.5×10^7
4.0×10^7
3.5×10^7
3.0×10^7
2.5×10^7
2.0×10^7
1.5×10^7
1.0×10^7
5.0×10^6
0.0
-5.0×10^6

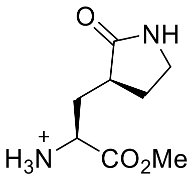
O
NH
Cl
H3N
CO2Me

**5**

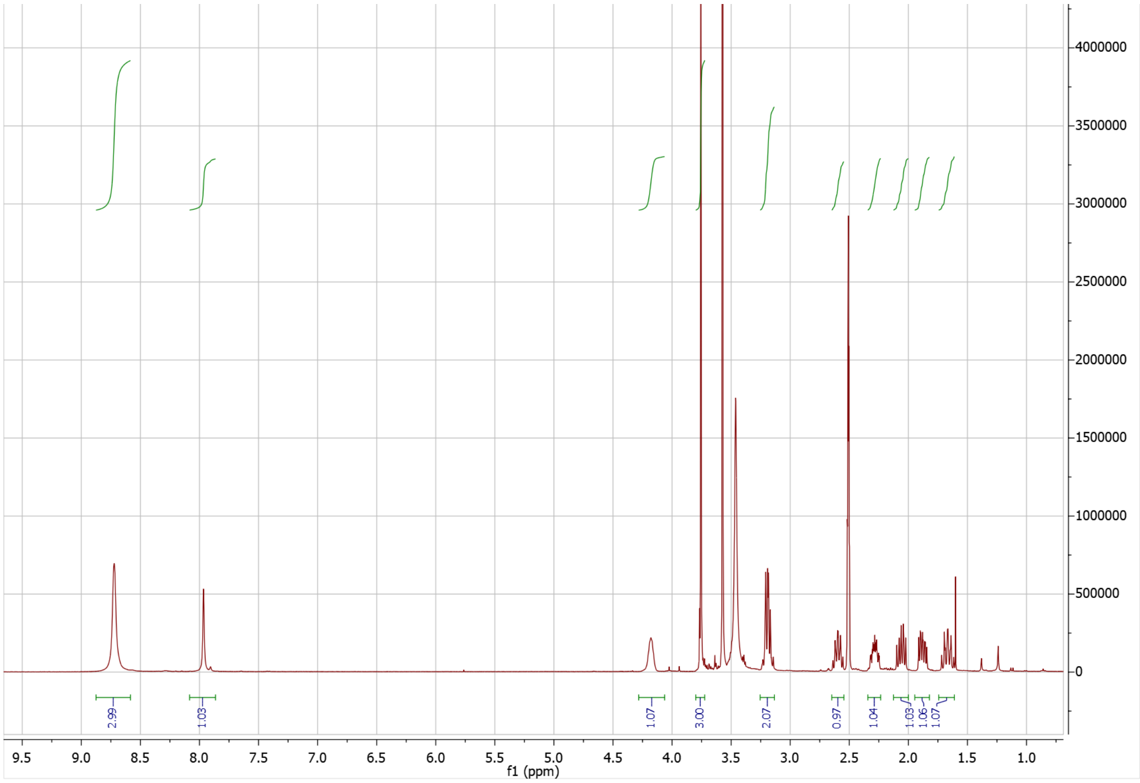
4000000
3500000
3000000
2500000
2000000
1500000
1000000
500000
0
2.99
1.03
1.07
3.00
2.07
0.97
1.04
1.03
1.06
1.07
9.5
9.0
8.5
8.0
7.5
7.0
6.5
6.0
5.5
5.0
4.5
4.0
3.5
3.0
2.5
2.0
1.5
1.0
f1 (ppm)

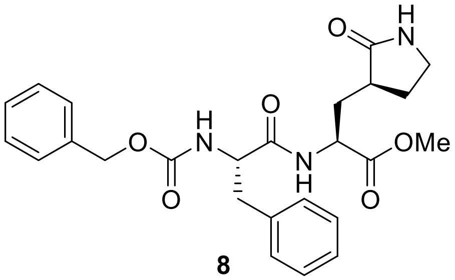
O
H
N
O
N
H
O
N
H
OMe
O
O

**8**

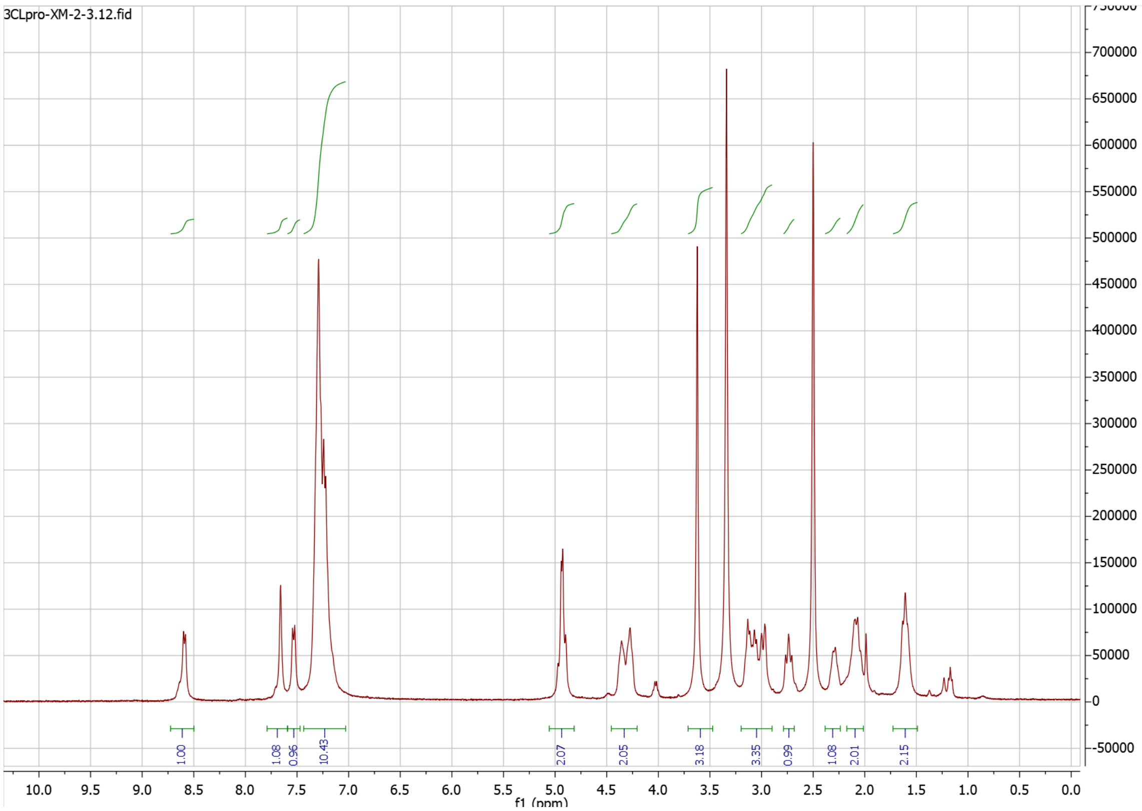
3CLpro-XM-2-3.12.fid
1.00
1.08
0.96
10.43
2.07
2.05
3.18
3.35
0.99
1.08
2.01
2.15
10.0 9.5 9.0 8.5 8.0 7.5 7.0 6.5 6.0 5.5 5.0 4.5 4.0 3.5 3.0 2.5 2.0 1.5 1.0 0.5 0.0
f1 (ppm)
700000
650000
600000
550000
500000
450000
400000
350000
300000
250000
200000
150000
100000
50000
0
-50000

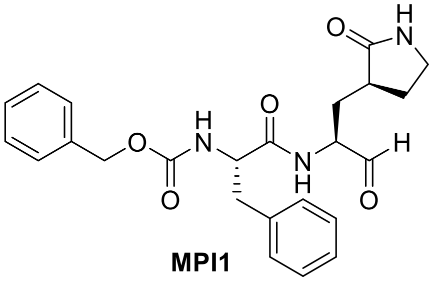

O
H
N
O
H
N
O
O
N
H
H
O


**MPI1**

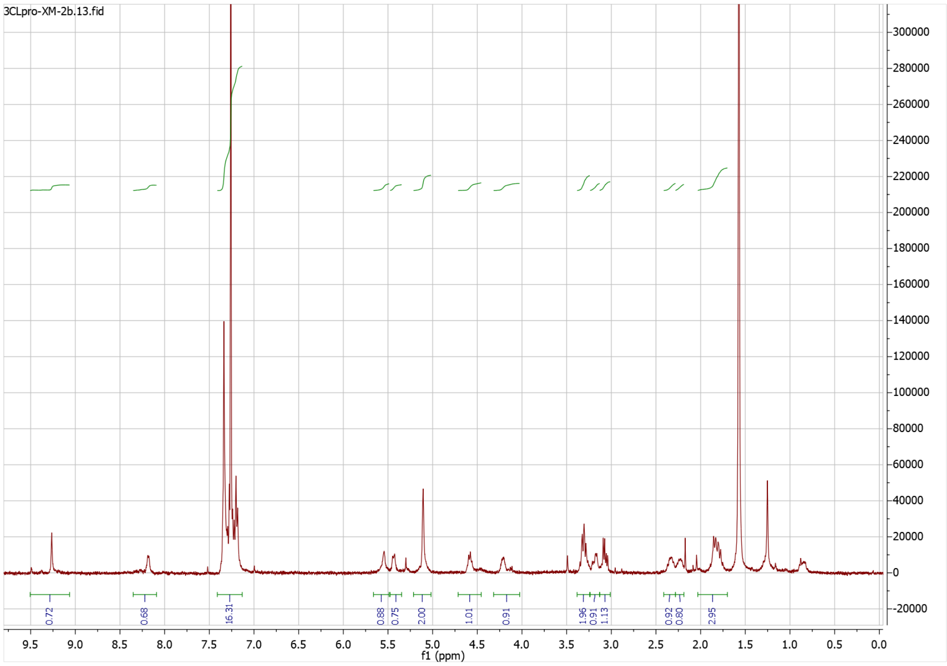

3CLpro-XM-2b.13.fid
0.72
0.68
16.31
0.88
0.75
2.00
1.01
0.91
1.96
0.91
1.13
0.92
0.80
2.95
f1 (ppm)


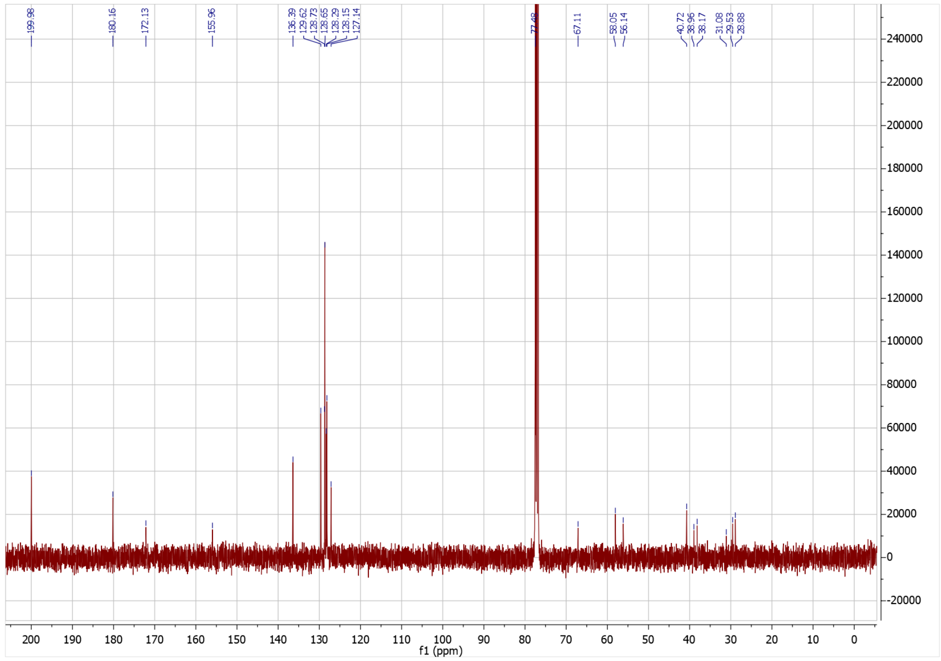

199.98
180.16
172.13
155.96
136.39
129.62
128.73
128.65
128.29
128.15
127.14
67.11
58.05
56.14
40.72
38.96
38.17
31.08
29.53
28.88
f1 (ppm)

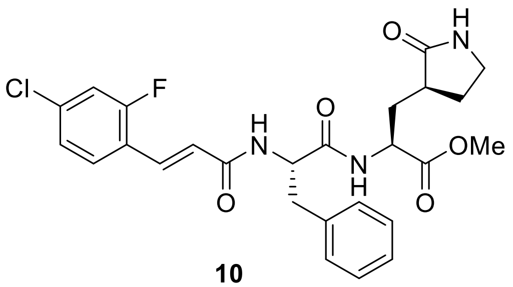

**10**

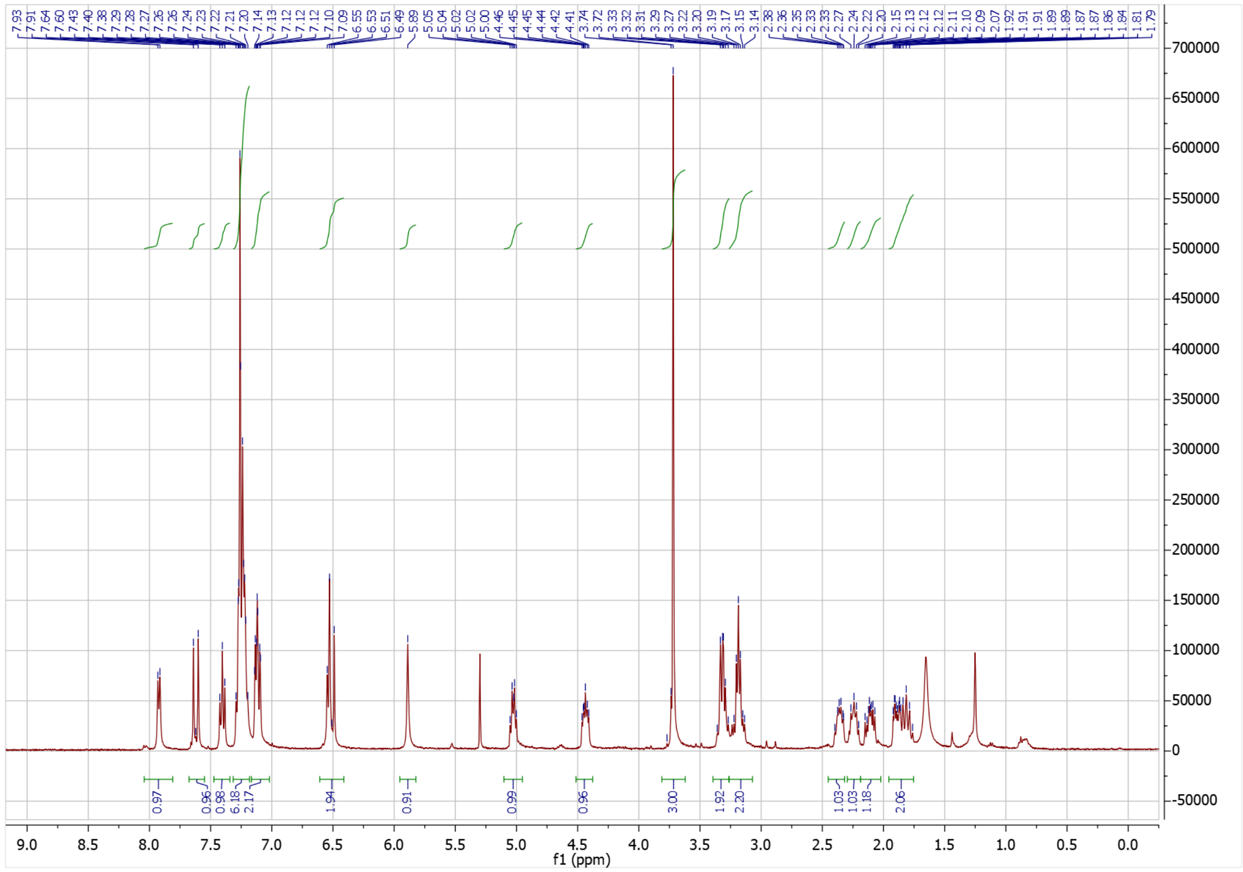

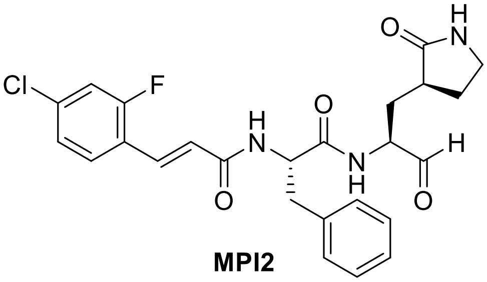
Cl
F
O
H
N
O
N
H
O
H
N
H
O

**MPI2**

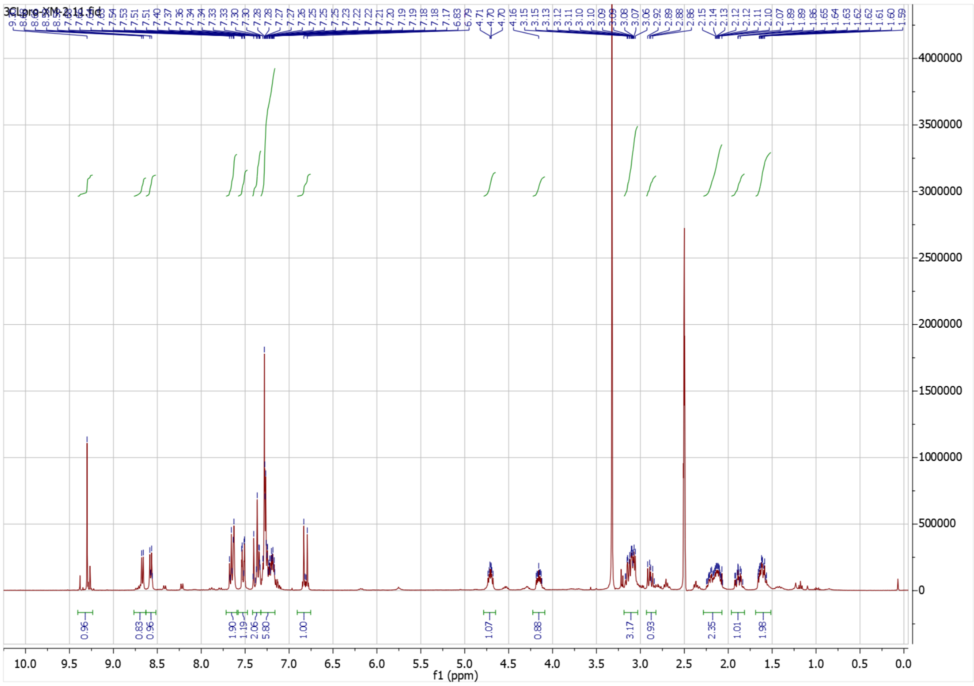
3CLpro-XM-2_14.fid
f1 (ppm)

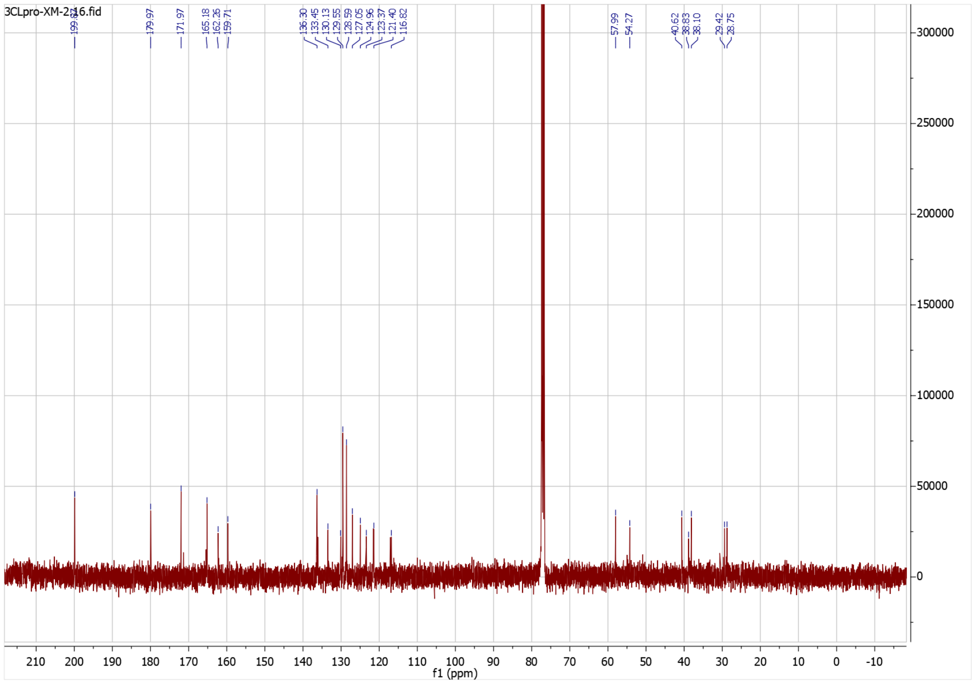
3CLpro-XM-2_16.fid
179.97
171.97
165.18
162.26
159.71
136.30
133.45
130.13
129.55
128.59
127.05
124.96
123.37
121.40
116.82
57.99
54.27
40.62
38.83
38.10
29.42
28.75
f1 (ppm)

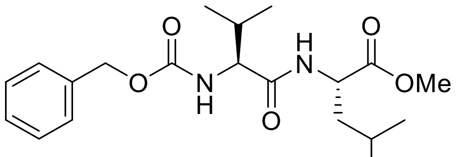

**11a**

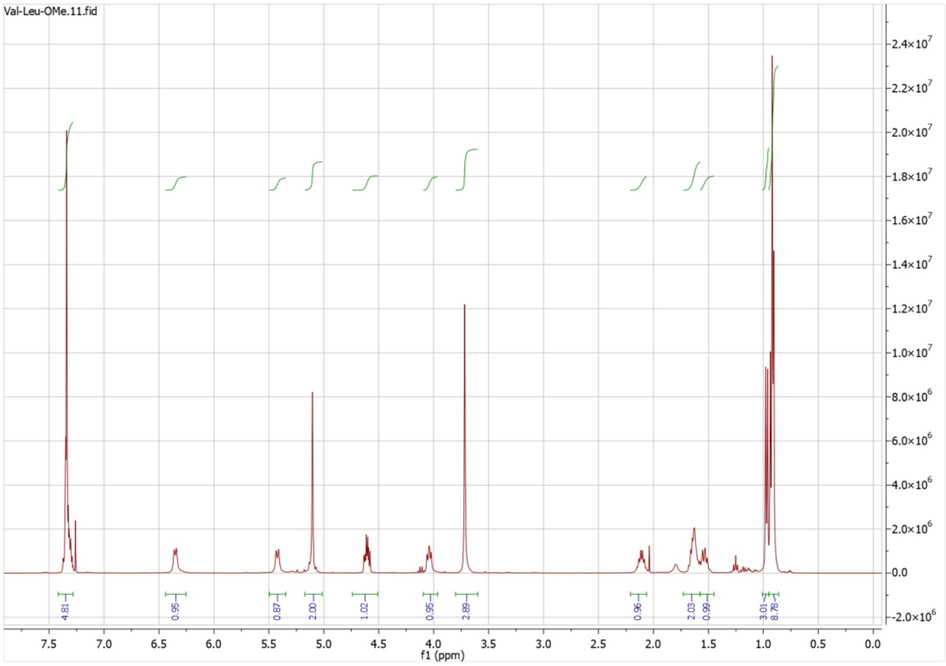


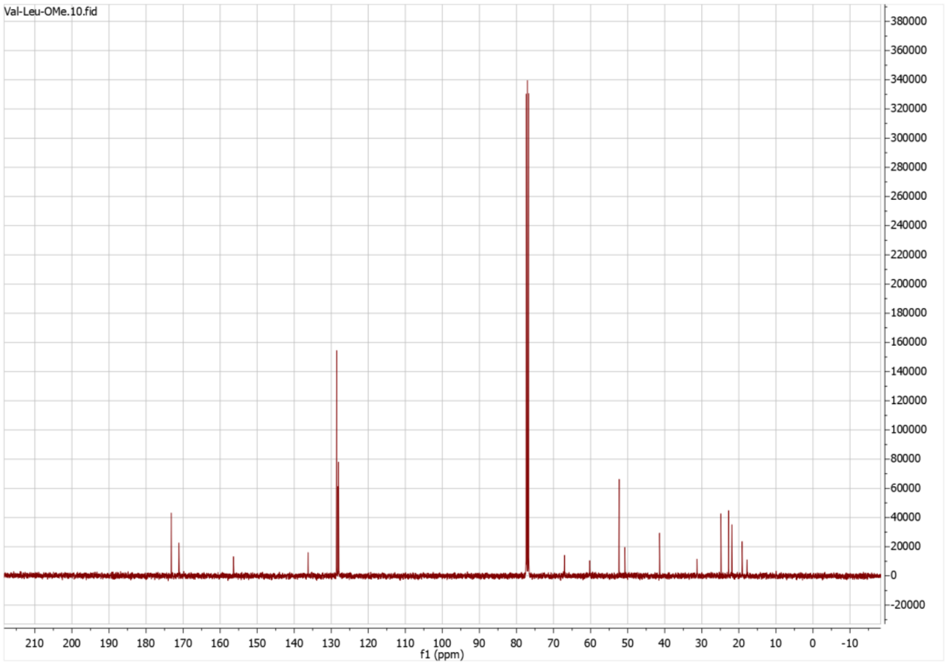

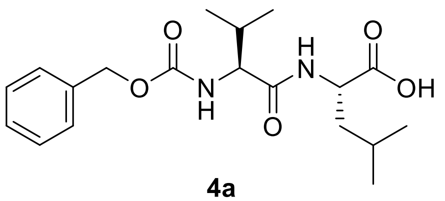

**4a**

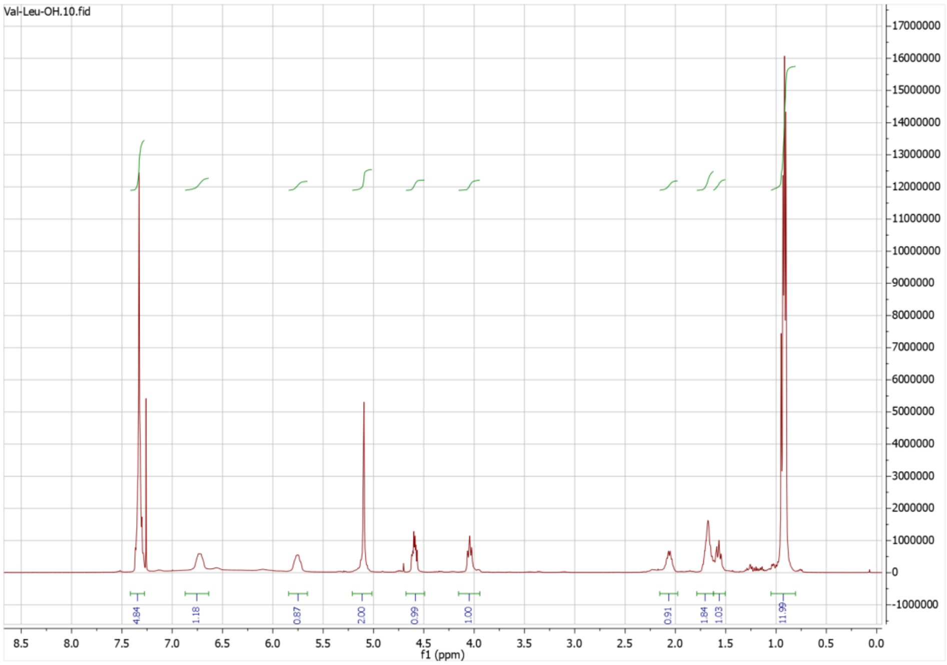
Val-Leu-OH.10.fid
4.84
1.18
0.87
2.00
0.99
1.00
0.91
1.84
1.03
11.99
f1 (ppm)

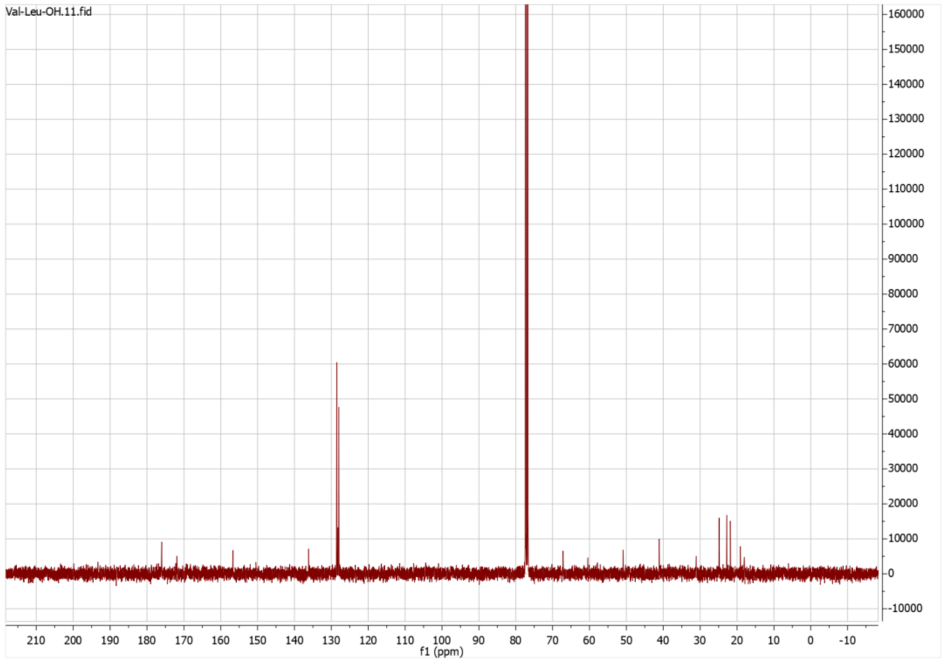
Val-Leu-OH.11.fid
f1 (ppm)

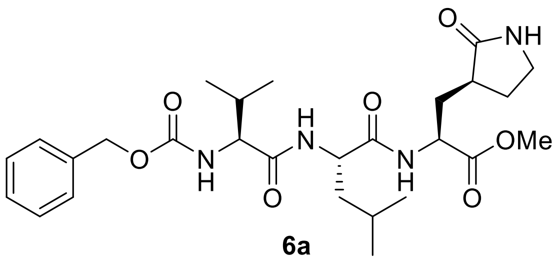
O
NH
O
H
N
O
OMe
O
N
H
O
N
H
O

**6a**

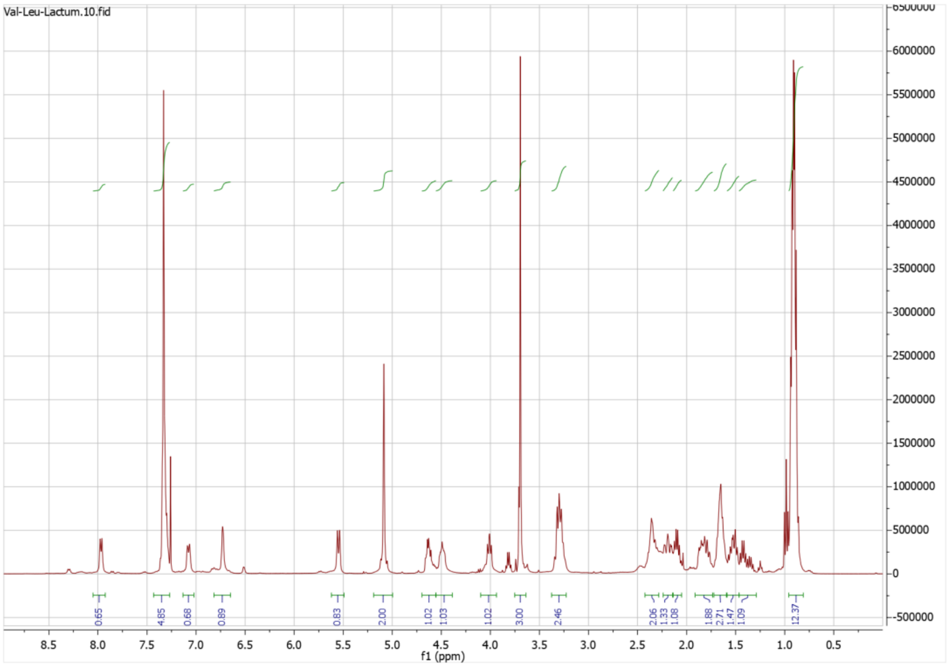
Val-Leu-Lactum.10.fid
0.65
4.85
0.68
0.89
0.83
2.00
1.02
1.03
1.02
3.00
2.46
2.06
1.33
1.08
1.88
2.71
1.47
1.09
12.37
f1 (ppm)

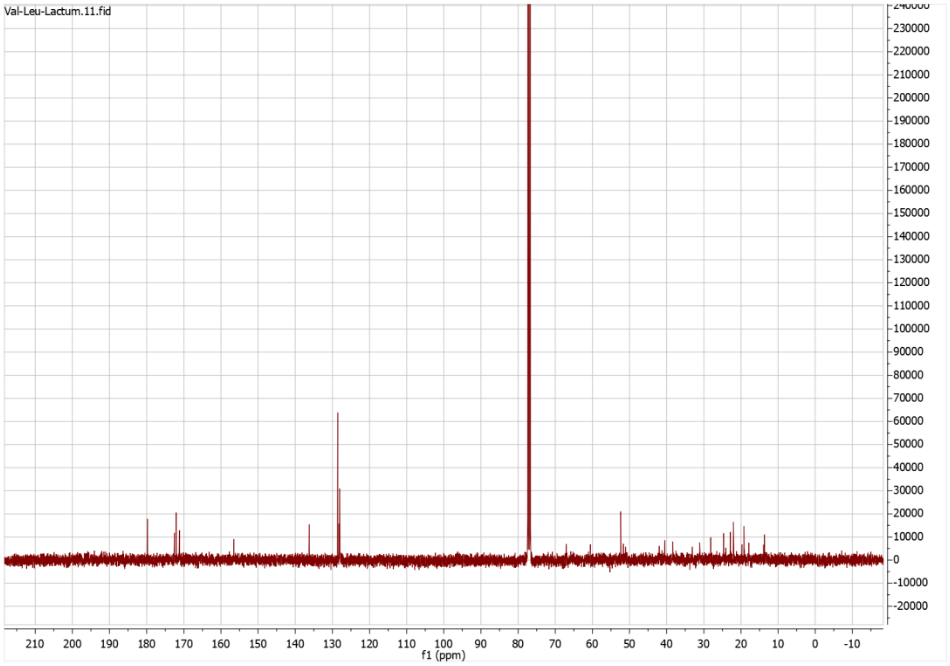
Val-Leu-Lactum.11.fid
f1 (ppm)

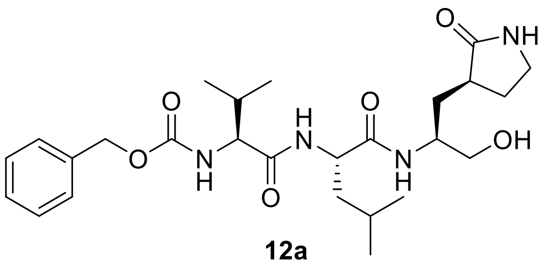
O
NH
O
O
H
N
OH
O
N
H
N
H
O

**12a**

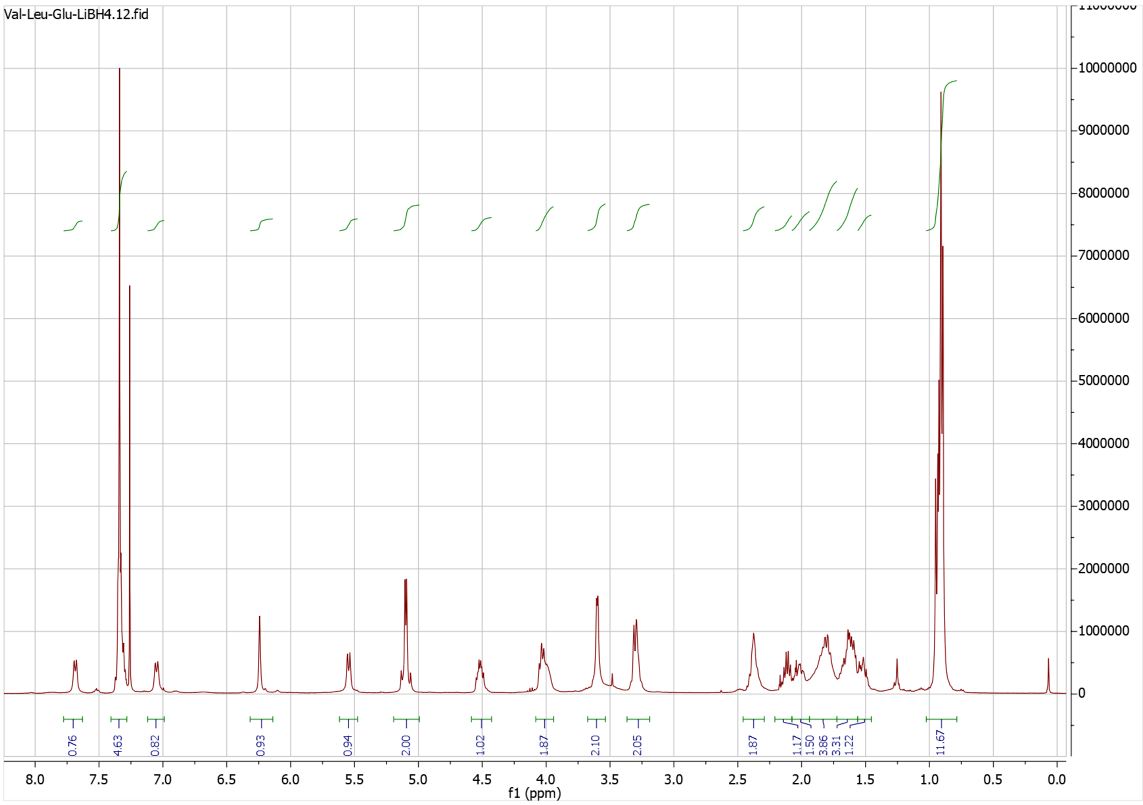
Val-Leu-Glu-LiBH4.12.fid
0.76
4.63
0.82
0.93
0.94
2.00
1.02
1.87
2.10
2.05
1.87
1.17
1.50
3.86
3.31
1.22
11.67
f1 (ppm)

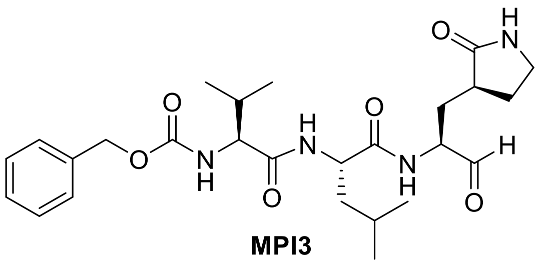

**MPI3**

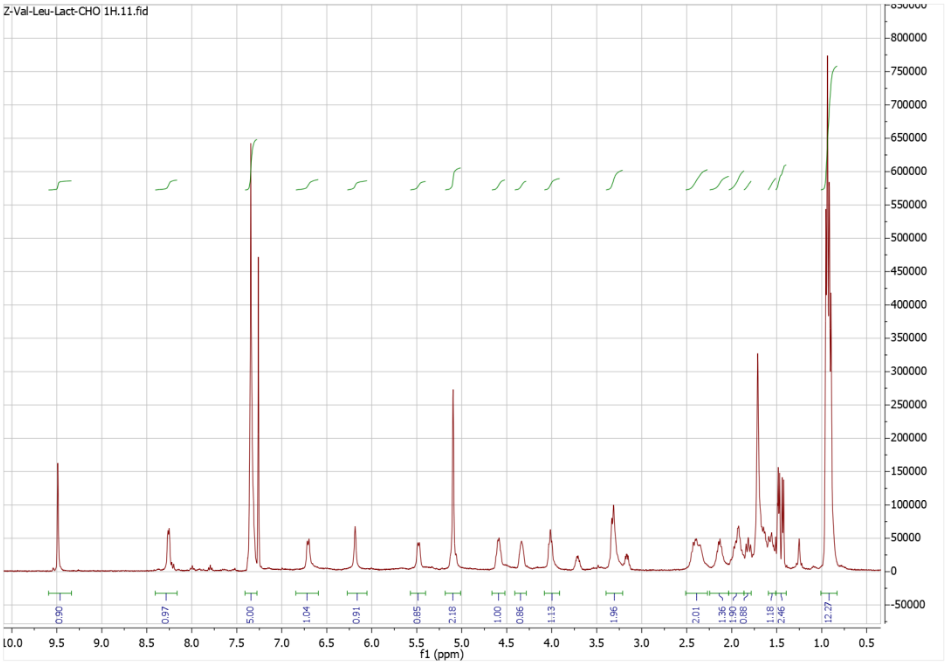
Z-Val-Leu-Lact-CHO 1H.11.fid
f1 (ppm)

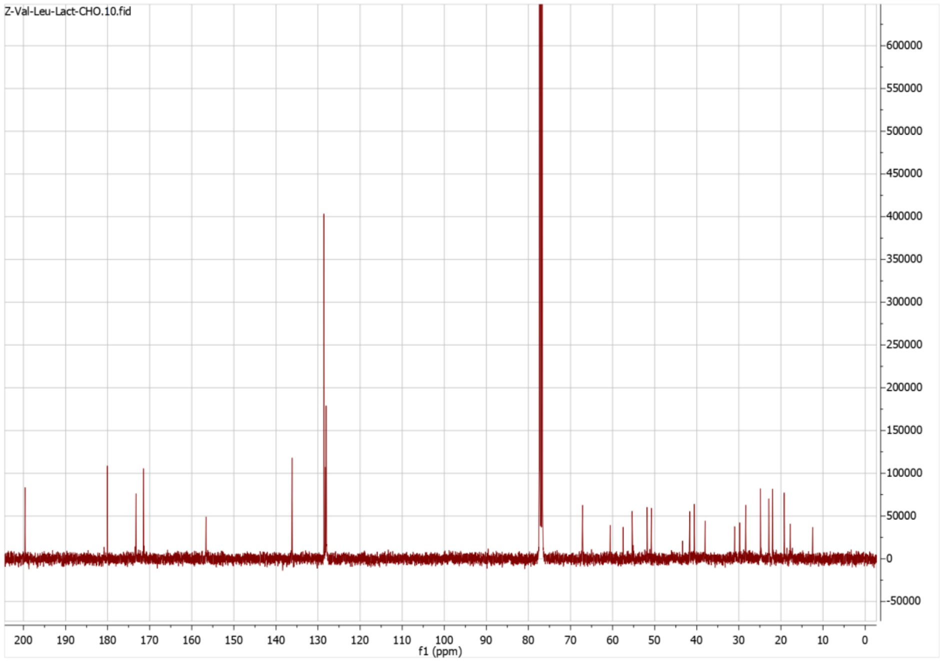
Z-Val-Leu-Lact-CHO.10.fid
f1 (ppm)

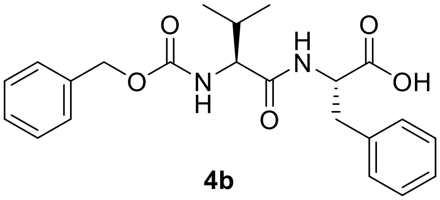

**4b**

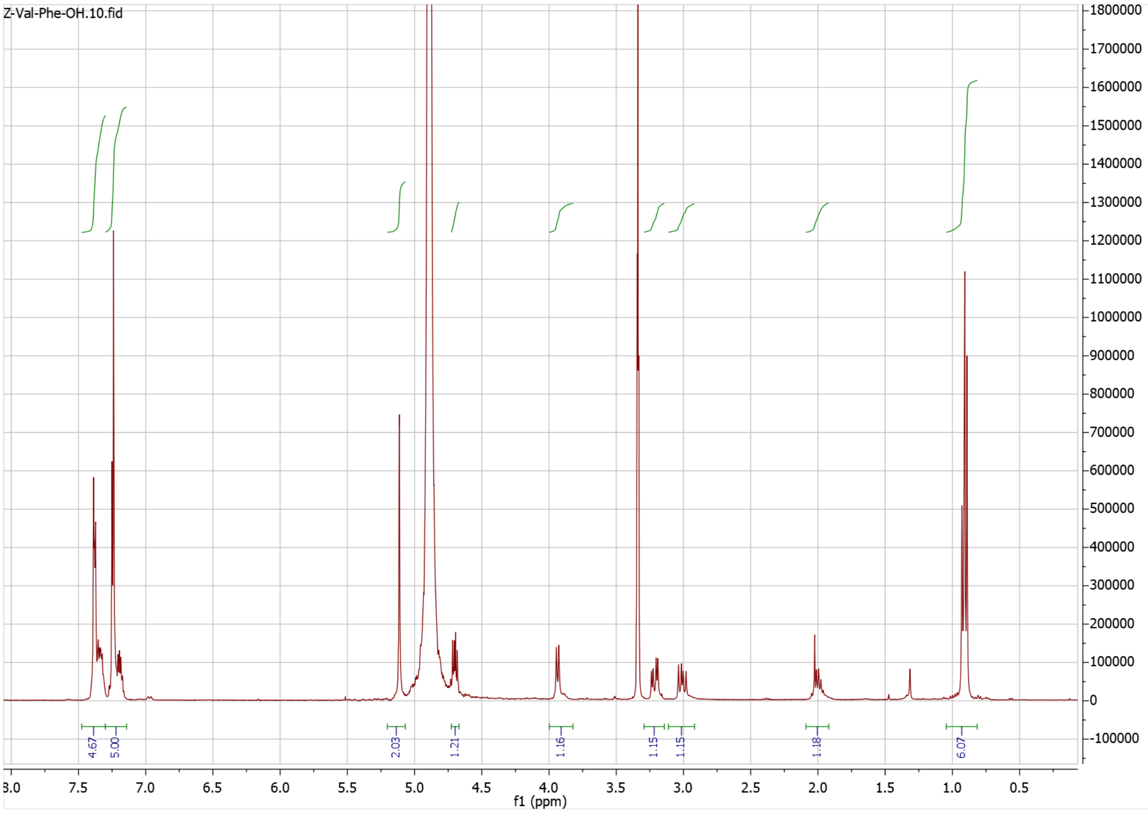

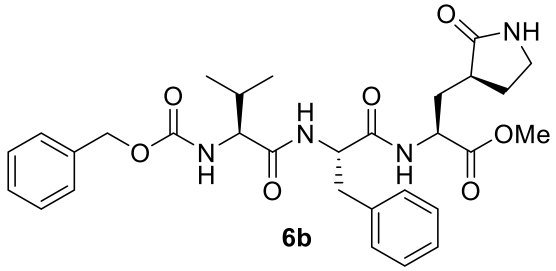

**6b**

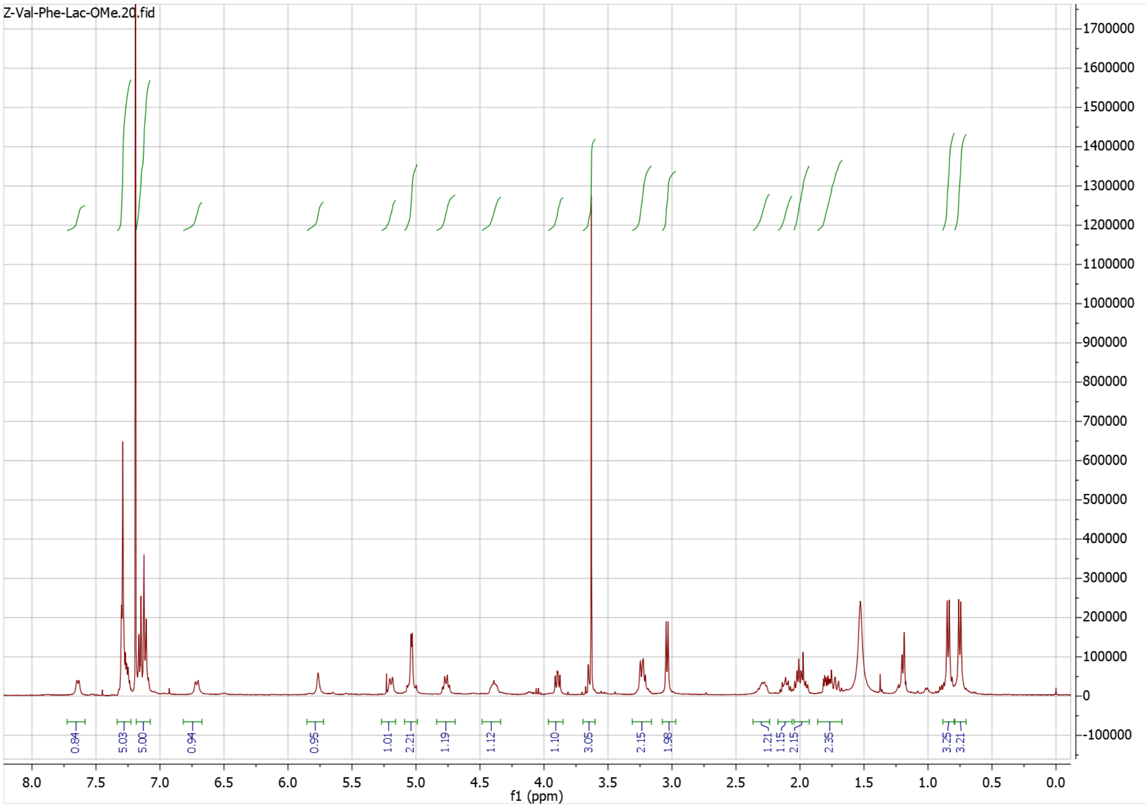

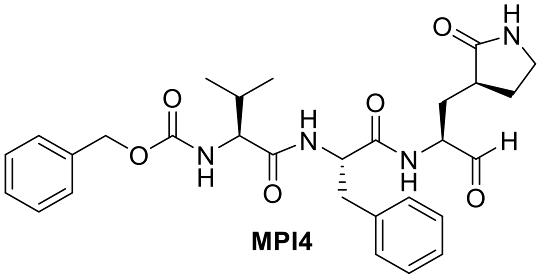

**MPI4**

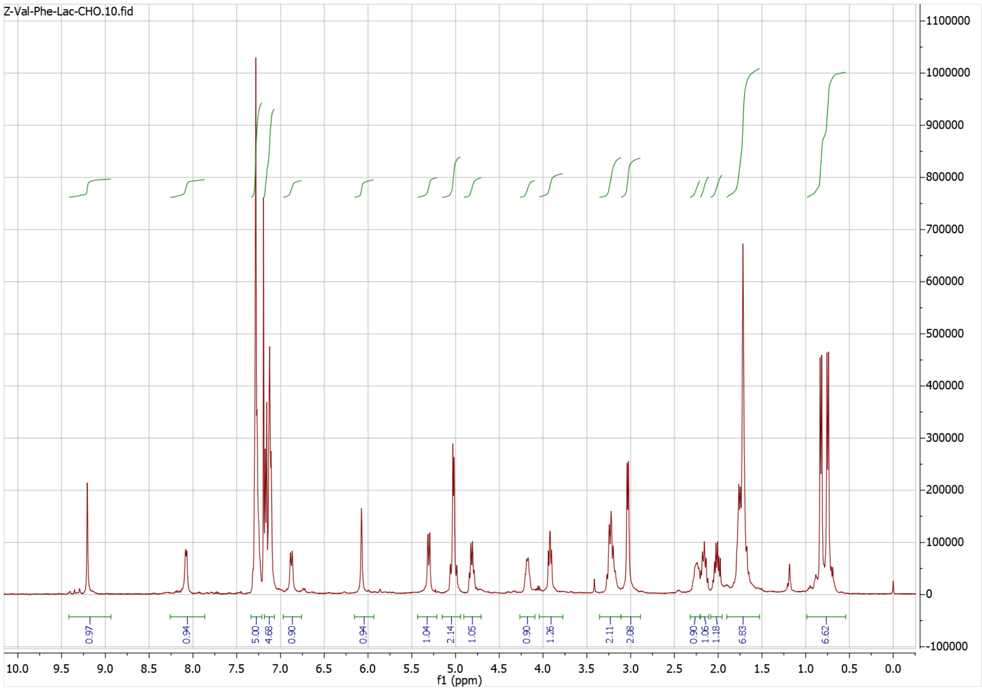


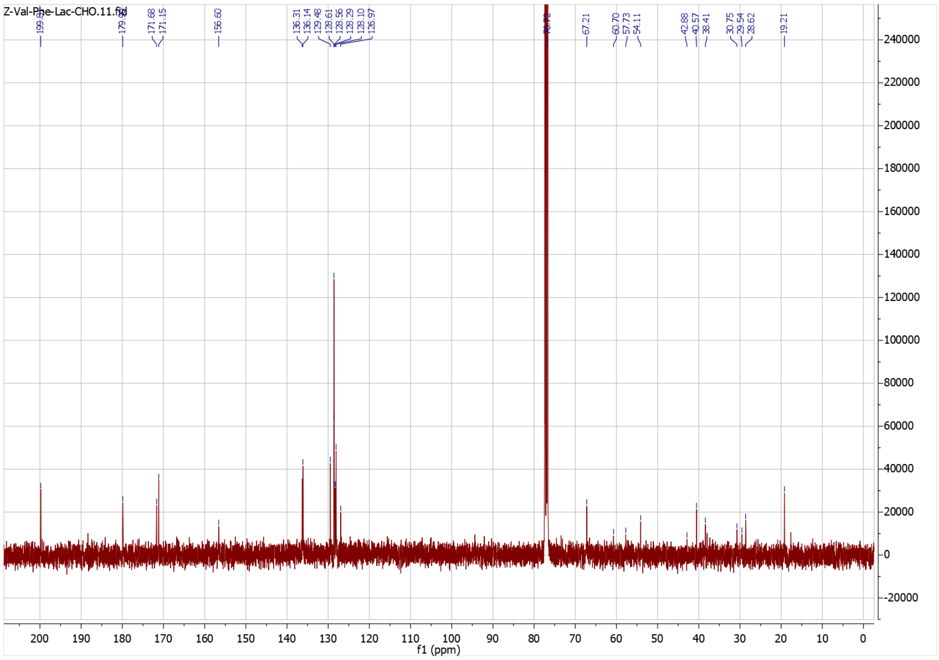

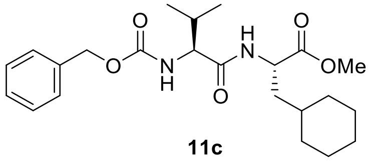
O
N
H
OMe

**11c**

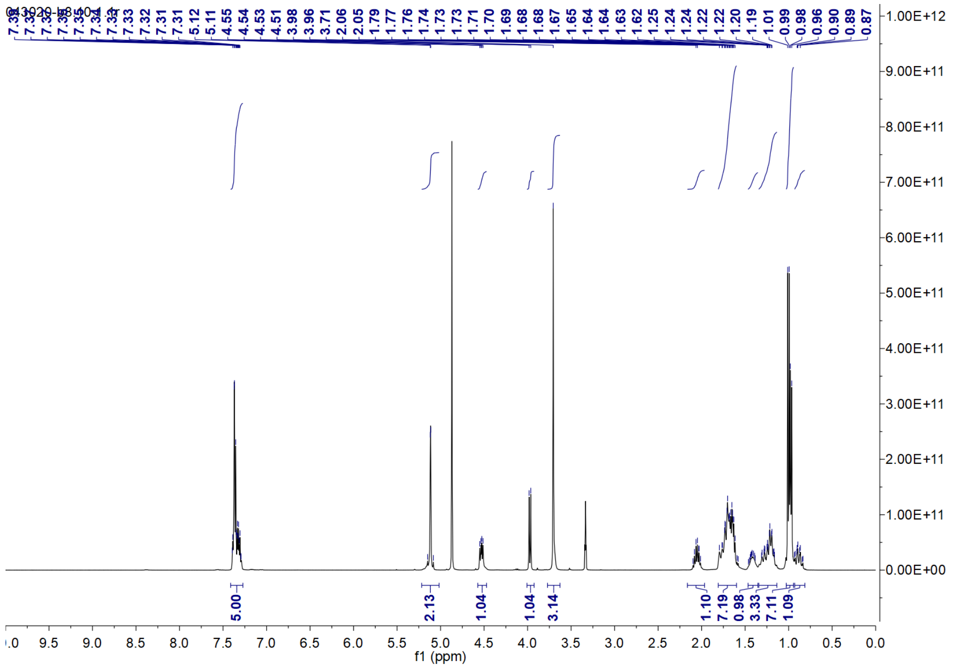
043020-b8.10.1.1r
7.33
7.32
7.31
7.31
5.12
5.11
4.55
4.54
4.53
4.51
3.98
3.96
3.71
2.06
2.05
1.79
1.77
1.76
1.74
1.73
1.73
1.71
1.70
1.69
1.68
1.68
1.67
1.65
1.64
1.64
1.63
1.62
1.25
1.24
1.24
1.22
1.22
1.20
1.19
1.01
0.99
0.98
0.96
0.90
0.89
0.87
5.00
2.13
1.04
1.04
3.14
1.10
7.19
0.98
3.33
7.11
1.09
f1 (ppm)

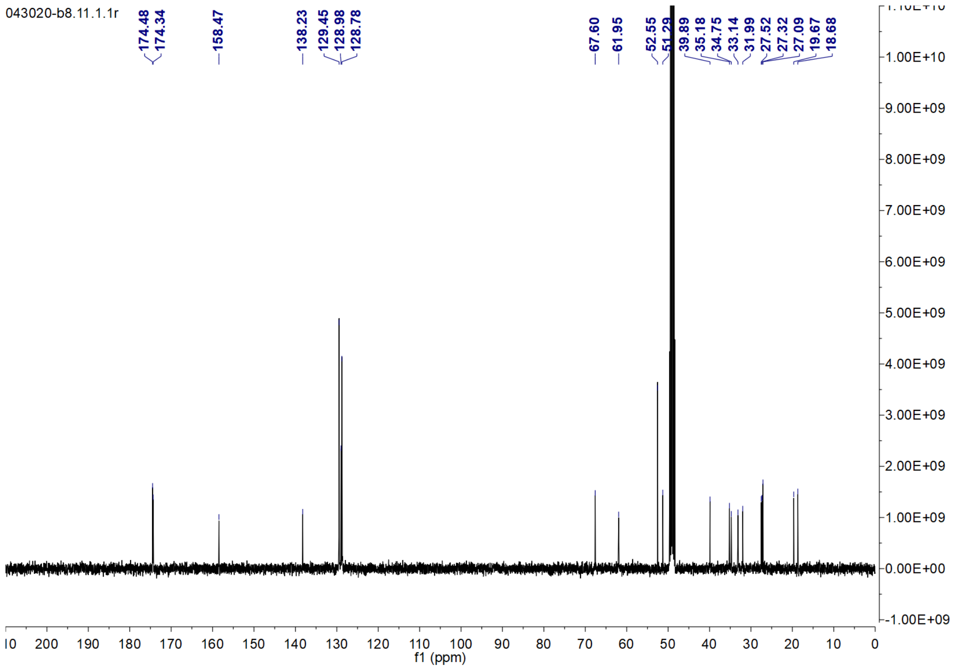
043020-b8.11.1.1r
174.48
174.34
158.47
138.23
129.45
128.98
128.78
67.60
61.95
52.55
51.29
39.89
35.18
34.75
33.14
31.99
27.52
27.32
27.09
19.67
18.68
f1 (ppm)

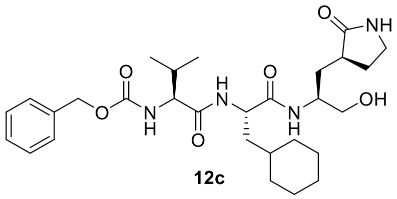
O
NH
H
N
O
O
O
N
H
N
H
OH

**12c**

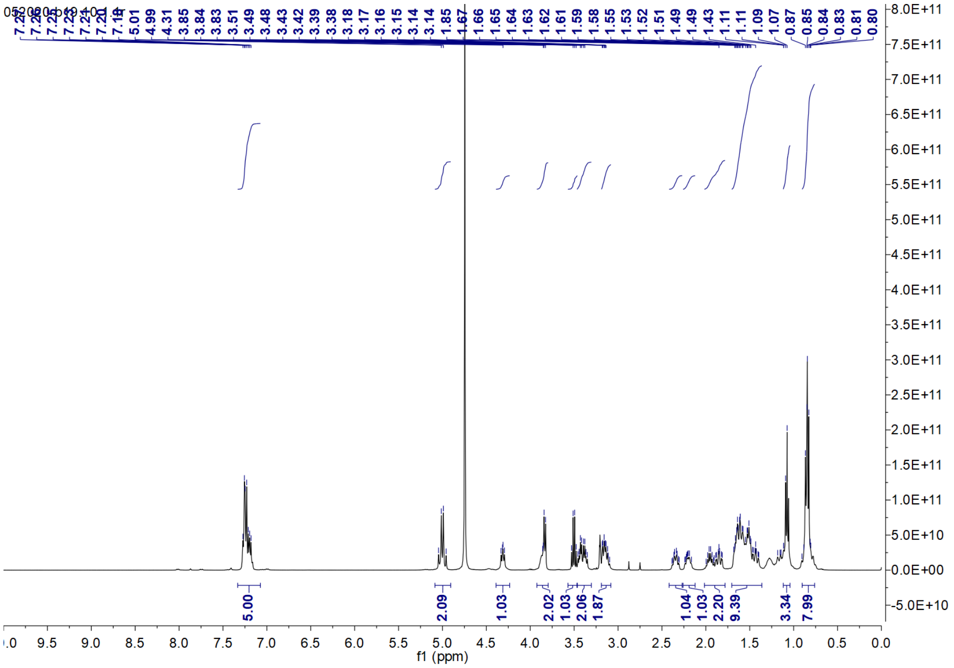
5.01
4.99
4.31
3.85
3.84
3.83
3.51
3.49
3.48
3.43
3.42
3.39
3.38
3.18
3.17
3.16
3.15
3.14
3.14
1.85
1.67
1.66
1.65
1.64
1.63
1.62
1.61
1.59
1.58
1.55
1.53
1.52
1.51
1.49
1.49
1.43
1.11
1.11
1.09
1.07
0.87
0.85
0.84
0.83
0.81
0.80
5.00
2.09
1.03
2.02
1.03
2.06
1.87
1.04
1.03
2.20
9.39
3.34
7.99
f1 (ppm)

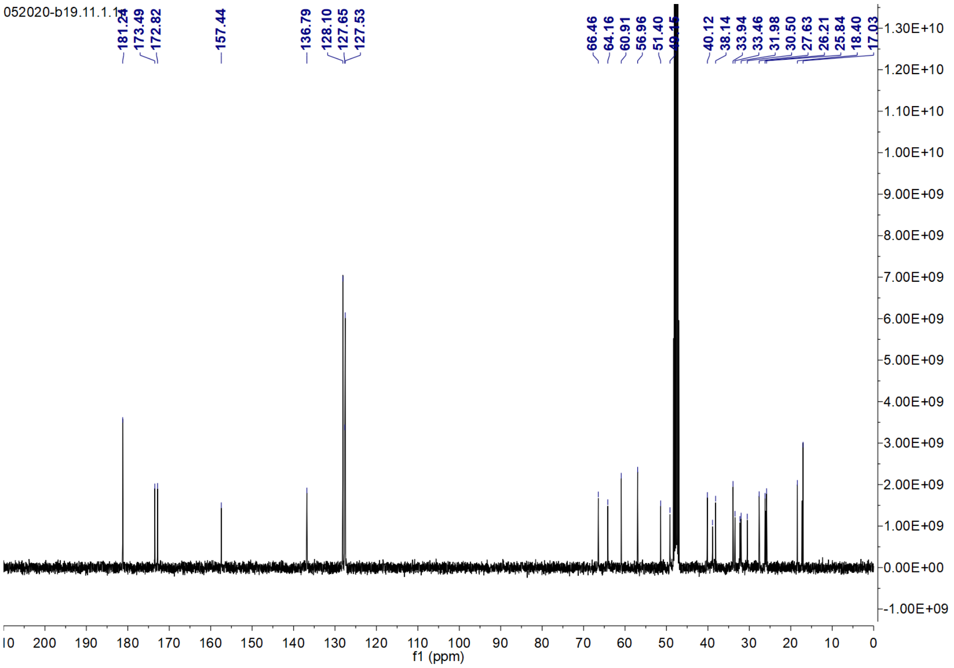
052020-b19.11.1.1
181.24
173.49
172.82
157.44
136.79
128.10
127.65
127.53
66.46
64.16
60.91
56.96
51.40
40.12
38.14
33.94
33.46
31.98
30.50
27.63
26.21
25.84
18.40
17.03
f1 (ppm)

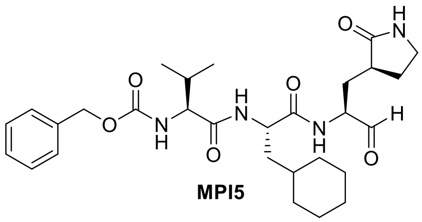

**MPI5**

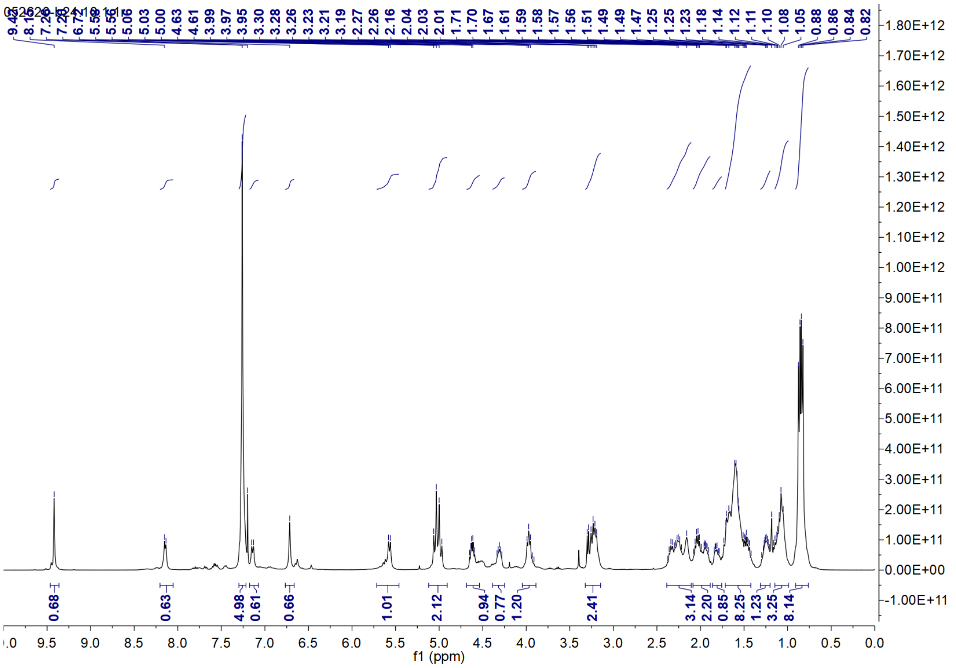
052620-b24.10.1r
f1 (ppm)
0.68
0.63
4.98
0.61
0.66
1.01
2.12
0.94
0.77
1.20
2.41
3.14
2.20
0.85
8.25
1.23
3.25
8.14

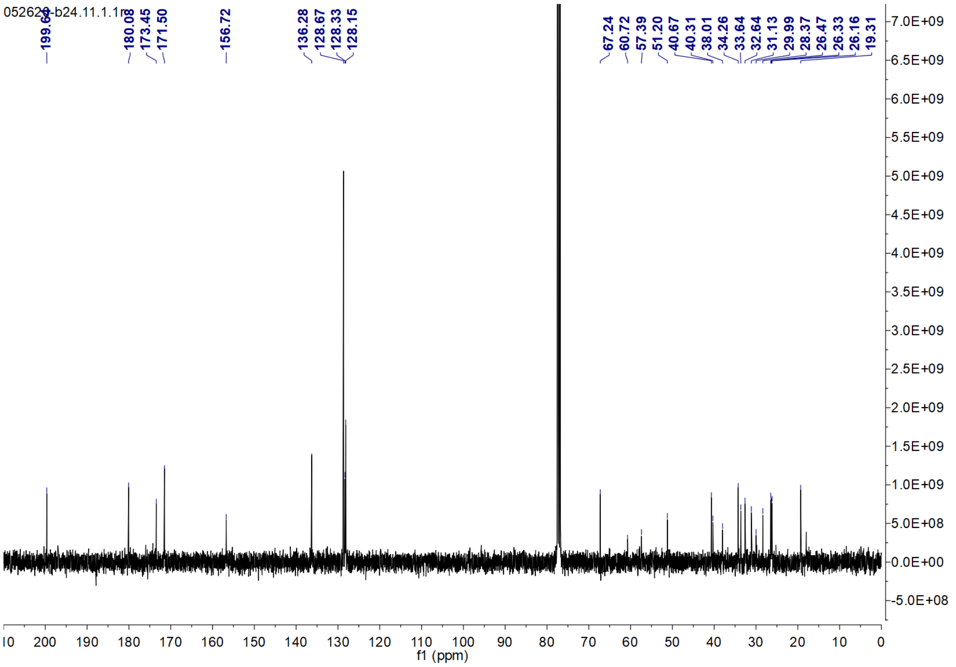
052620-b24.11.1.1r
199.60
180.08
173.45
171.50
156.72
136.28
128.67
128.33
128.15
67.24
60.72
57.39
51.20
40.67
40.31
38.01
34.26
33.64
32.64
31.13
29.99
28.37
26.47
26.33
26.16
19.31
f1 (ppm)

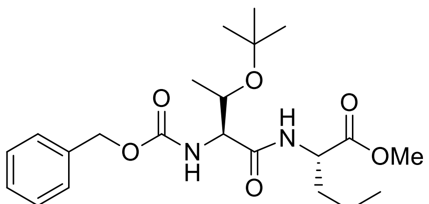


**11d**

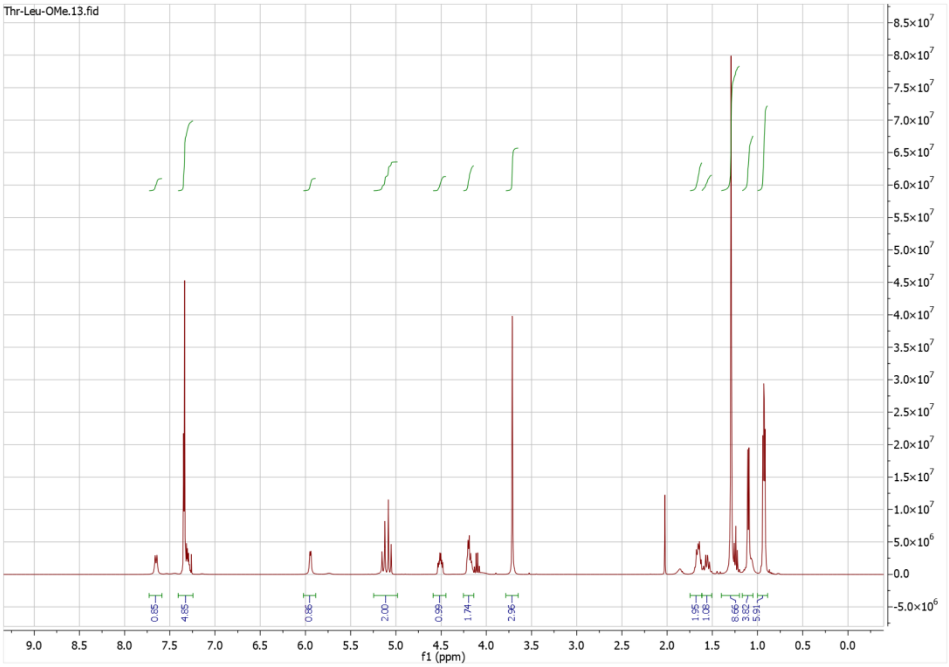


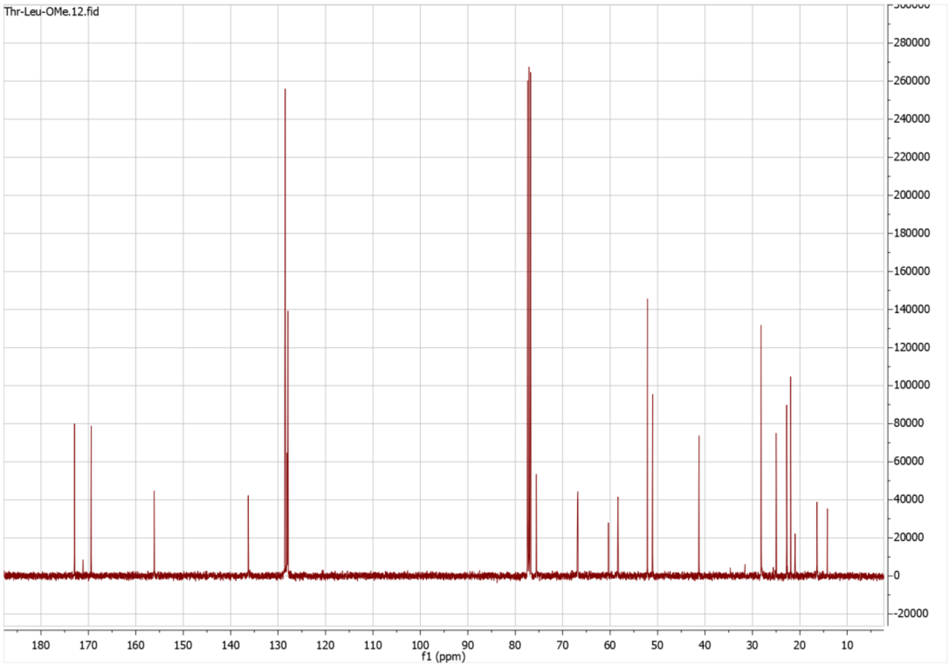

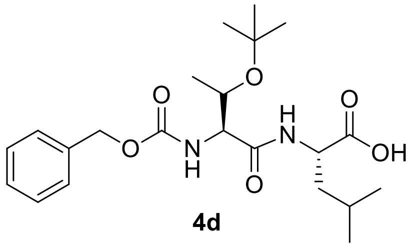

**4d**

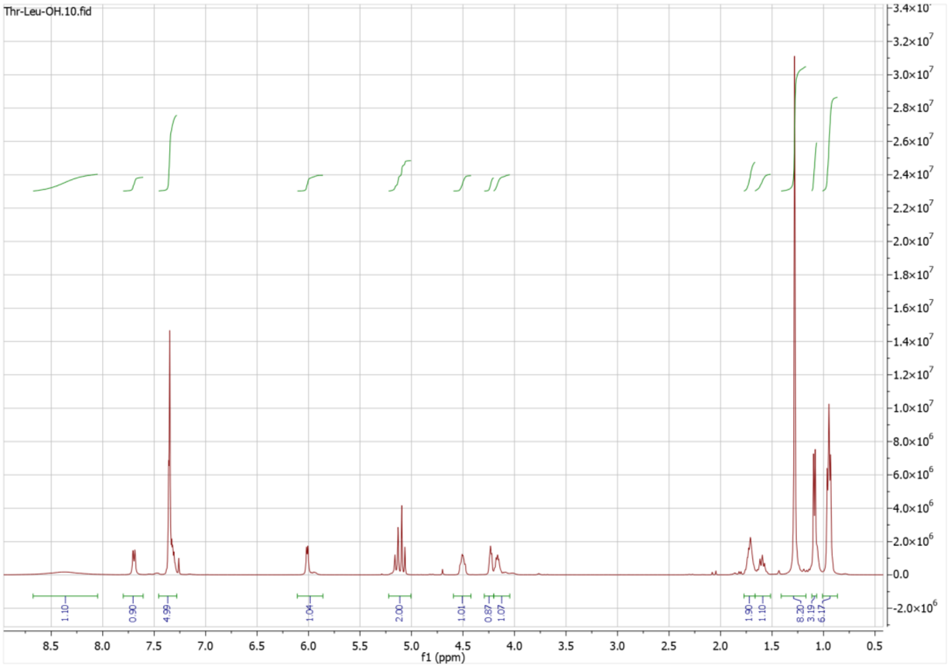


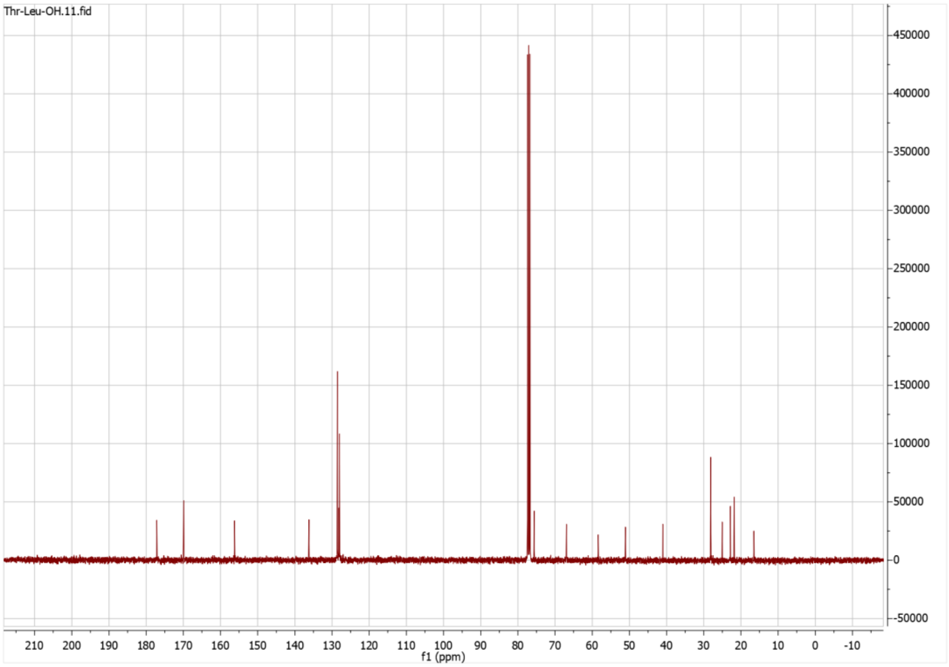

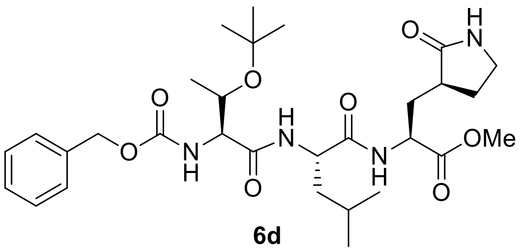

**6d**

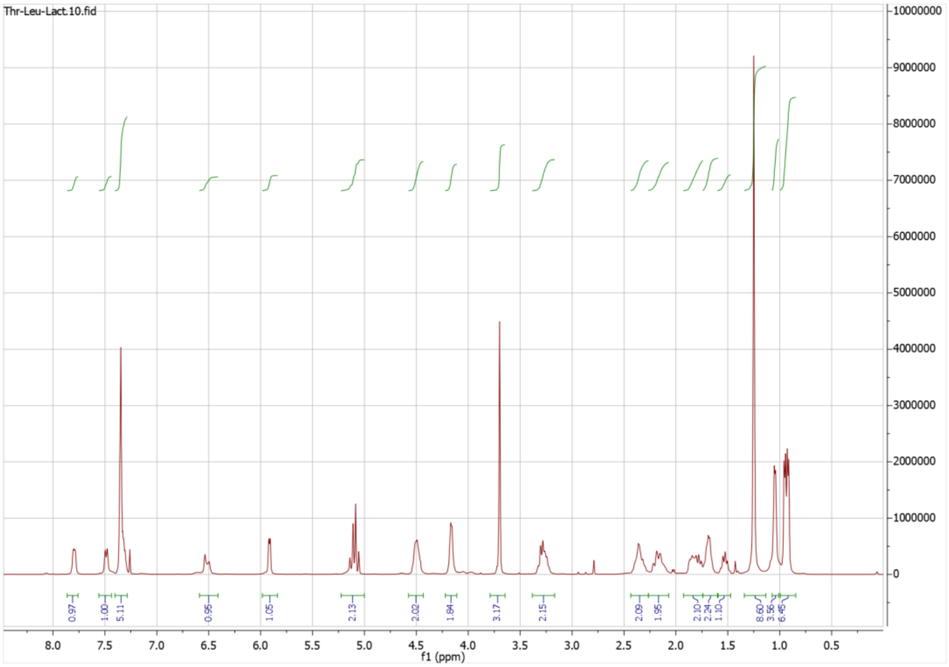


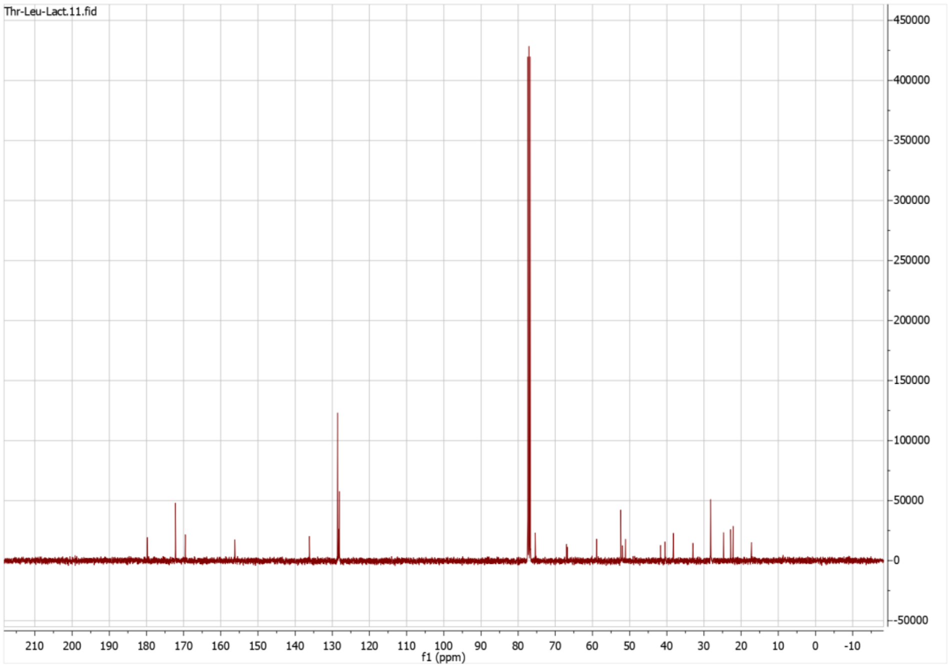

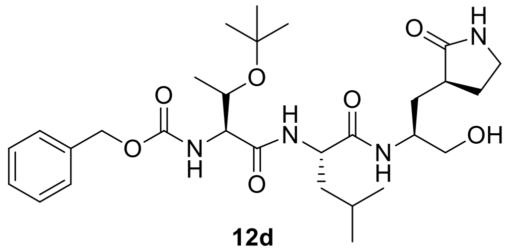

**12d**

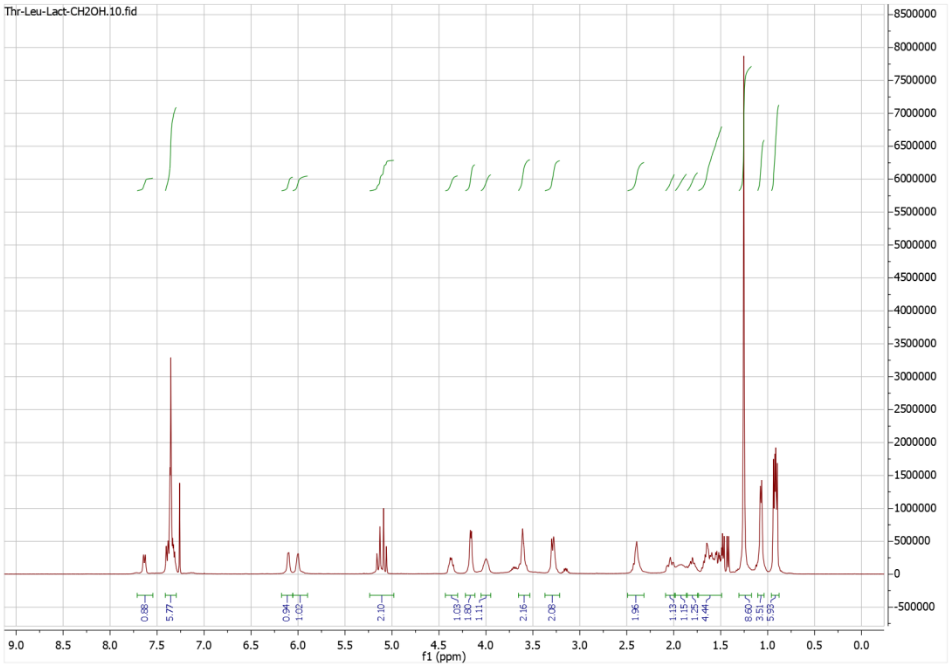
Thr-Leu-Lact-CH2OH.10.fid
0.88
5.77
0.94
1.02
2.10
1.03
1.80
1.11
2.16
2.08
1.96
1.13
1.15
1.25
4.44
8.60
3.51
5.93
f1 (ppm)

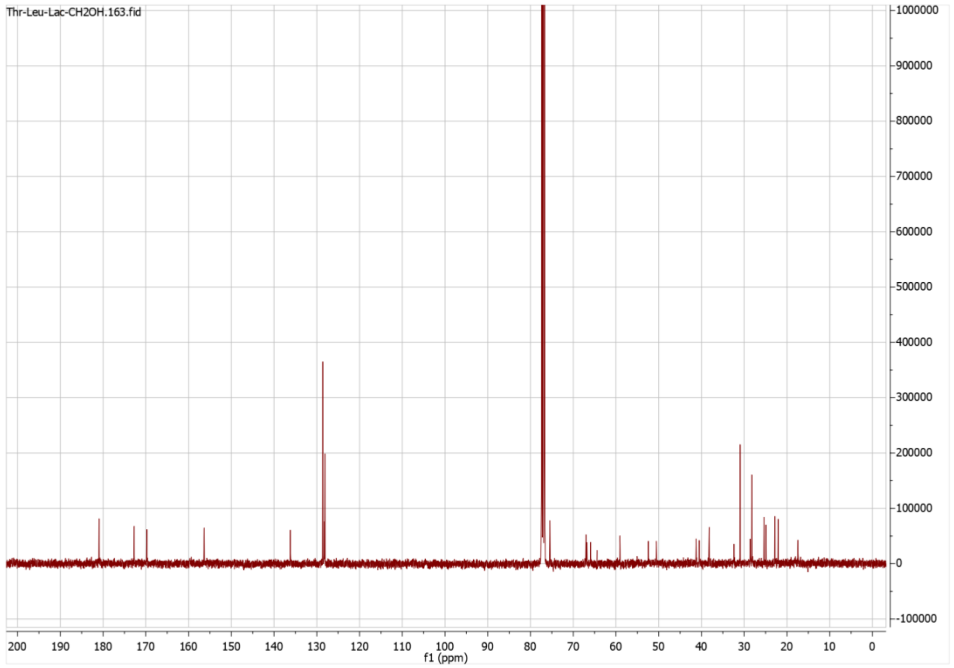
Thr-Leu-Lac-CH2OH.163.fid
f1 (ppm)

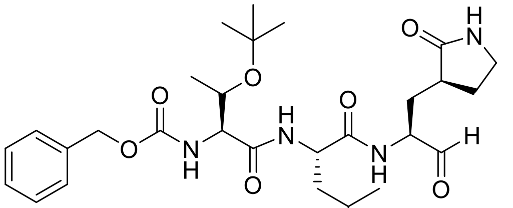

**MPI6**

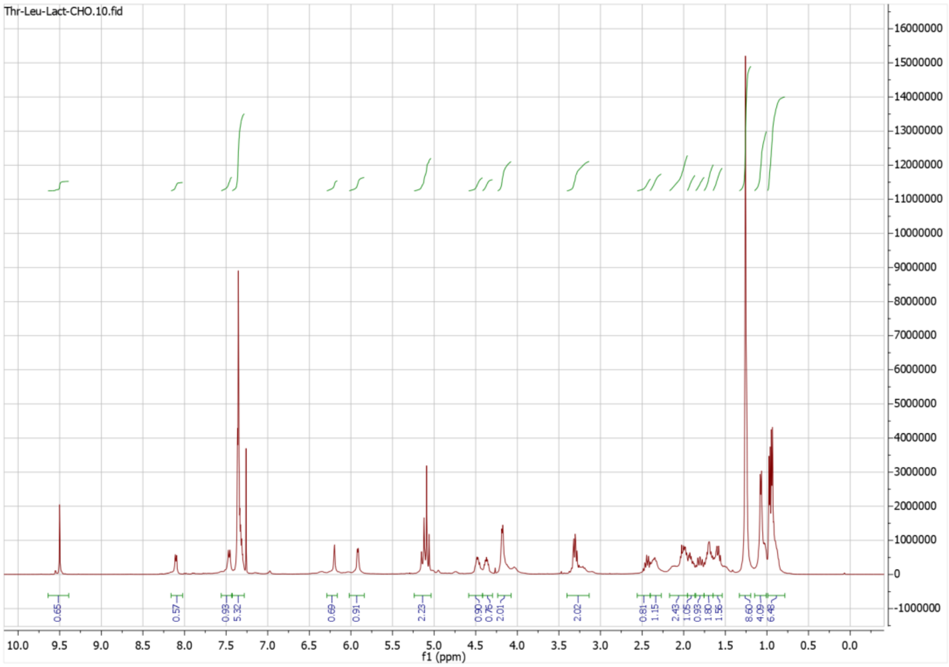


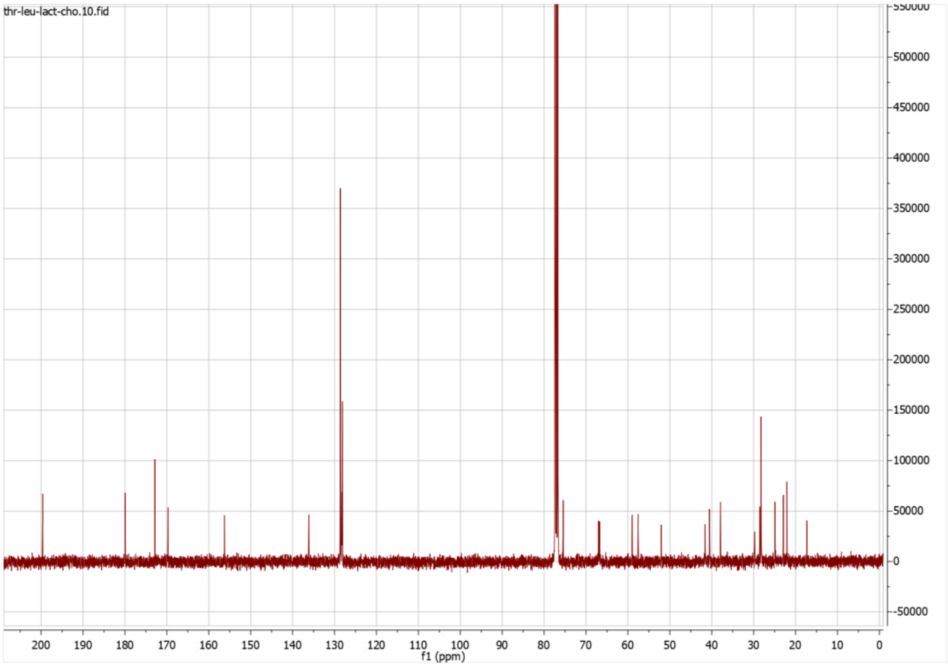

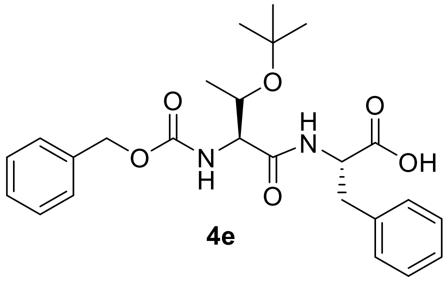

**4e**

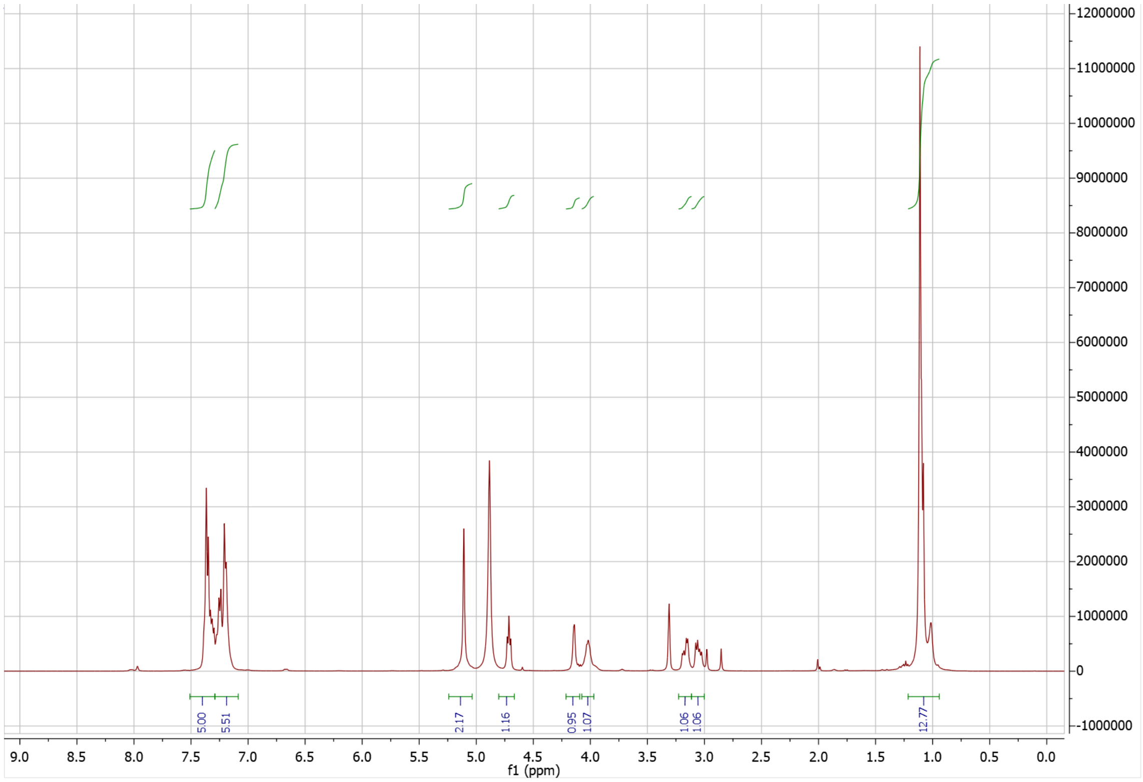

12000000
11000000
10000000
9000000
8000000
7000000
6000000
5000000
4000000
3000000
2000000
1000000
0
-1000000
5.00
5.51
2.17
1.16
0.95
1.07
1.06
1.06
12.77
9.0 8.5 8.0 7.5 7.0 6.5 6.0 5.5 5.0 4.5 4.0 3.5 3.0 2.5 2.0 1.5 1.0 0.5 0.0
f1 (ppm)

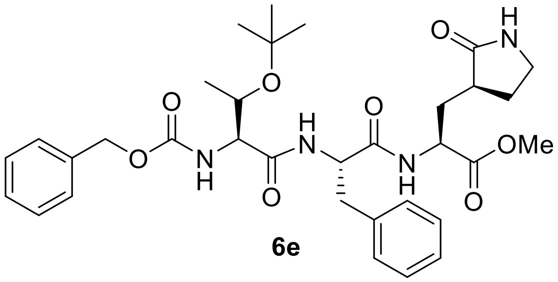

**6e**

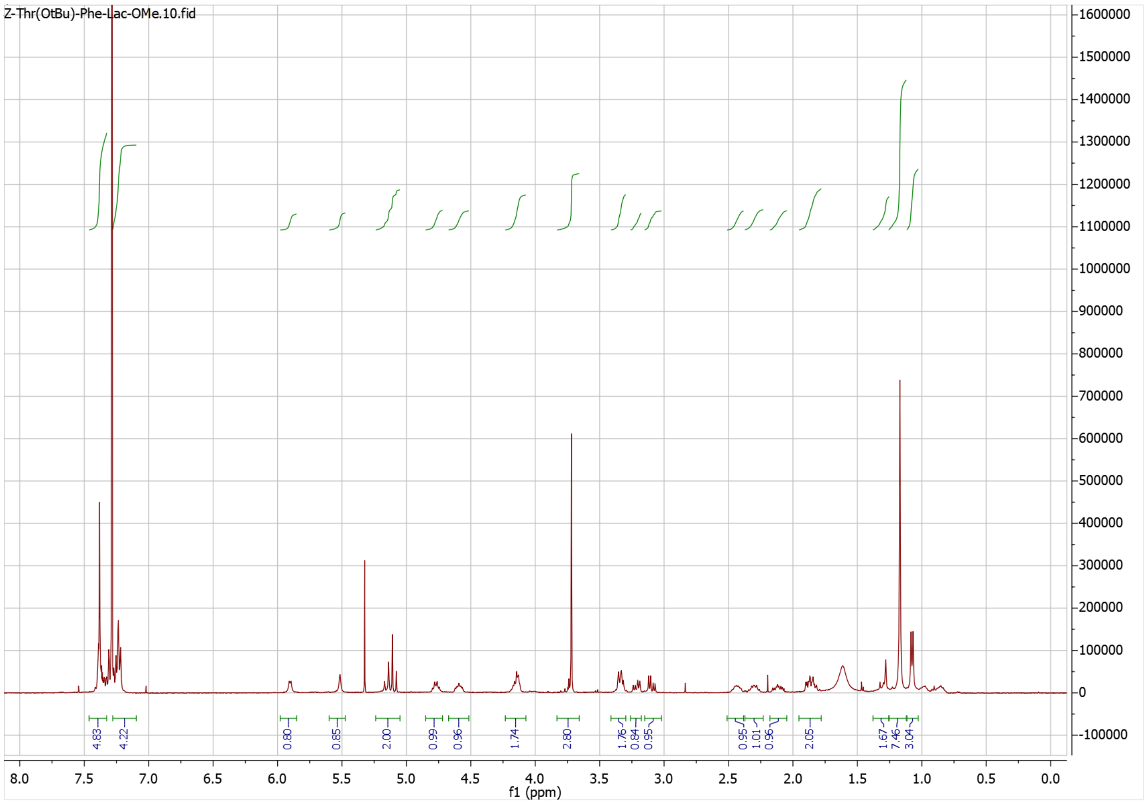

Z-Thr(OtBu)-Phe-Lac-OMe.10.fid
1600000
1500000
1400000
1300000
1200000
1100000
1000000
900000
800000
700000
600000
500000
400000
300000
200000
100000
0
-100000
4.83
4.22
0.80
0.85
2.00
0.99
0.96
1.74
2.80
1.76
0.84
0.95
0.95
1.01
0.96
2.05
1.67
7.46
3.04
8.0
7.5
7.0
6.5
6.0
5.5
5.0
4.5
4.0
3.5
3.0
2.5
2.0
1.5
1.0
0.5
0.0
f1 (ppm)

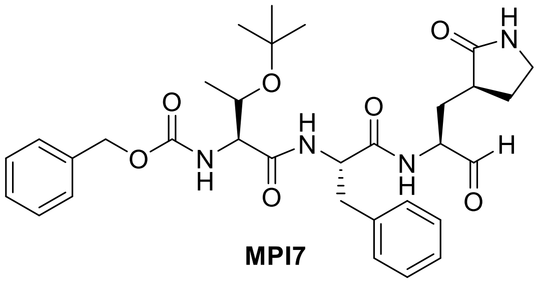

**MPI7**

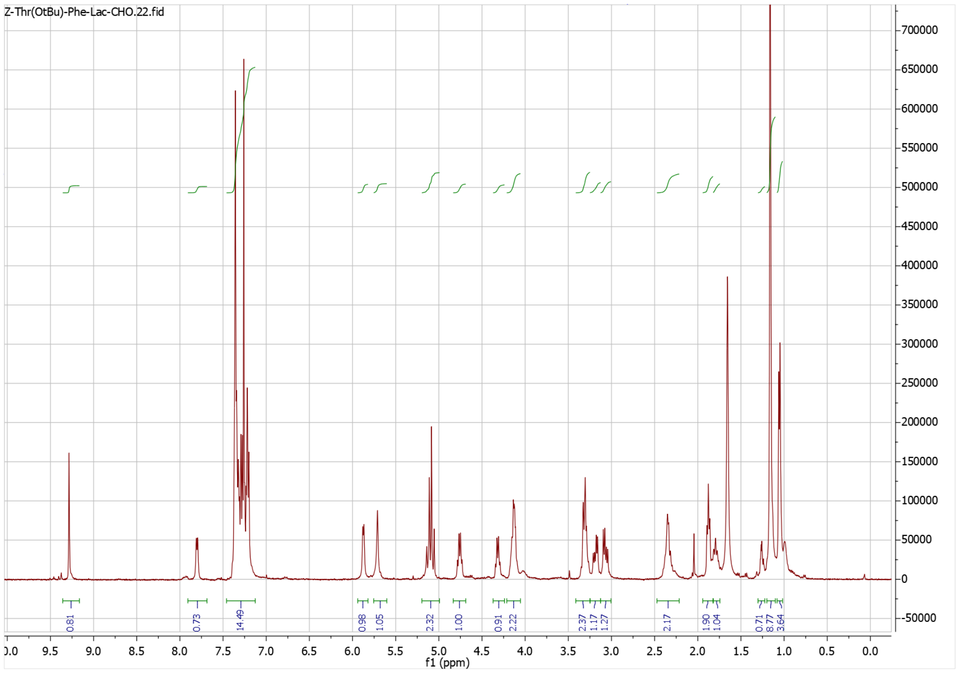

Z-Thr(OtBu)-Phe-Lac-CHO.22.fid
0.81
0.73
14.49
0.98
1.05
2.32
1.00
0.91
2.22
2.37
1.17
1.27
2.17
1.90
1.04
0.71
8.77
3.64
f1 (ppm)


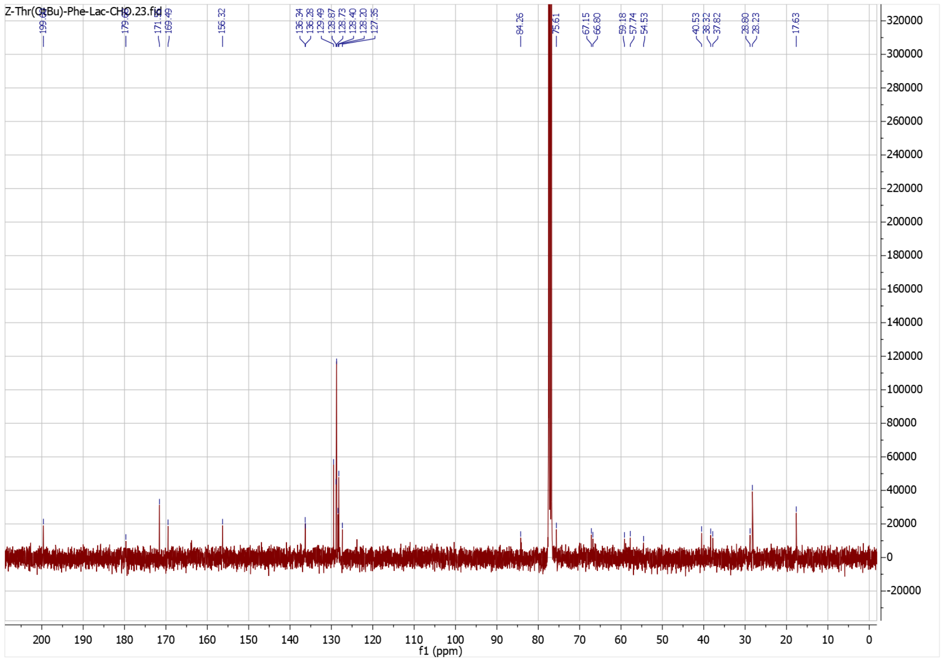

156.32
136.34
136.28
129.49
128.87
128.73
128.40
128.20
127.35
84.26
75.61
67.15
66.80
59.18
57.74
54.53
40.53
38.32
37.82
28.80
28.23
17.63
f1 (ppm)

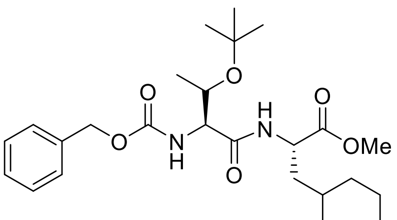

O
N
H
O
OMe


**11f**

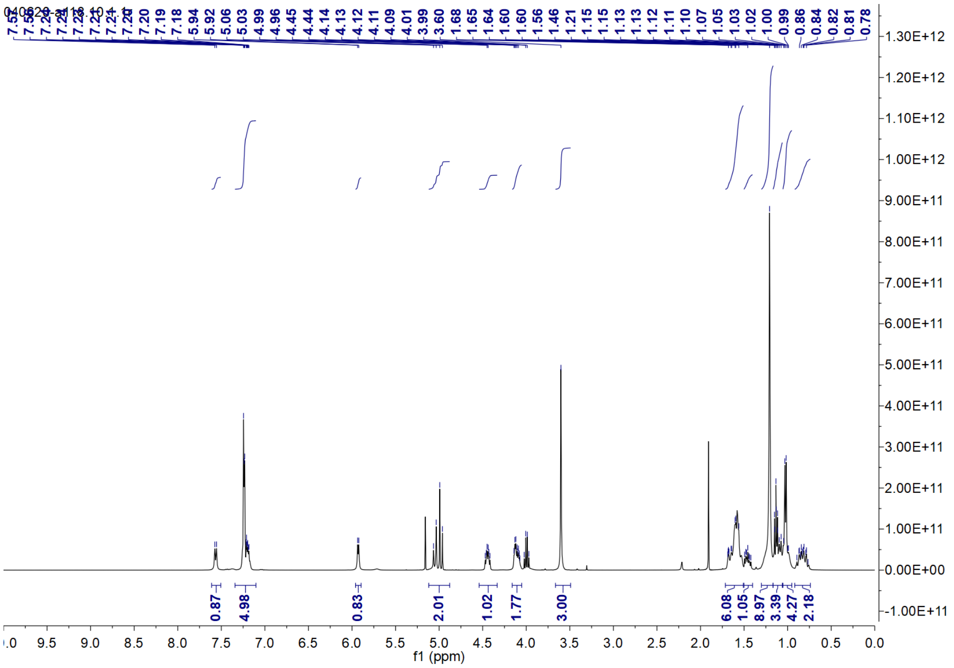

7.22
7.21
7.21
7.20
7.20
7.19
7.18
5.94
5.92
5.06
5.03
4.99
4.96
4.45
4.44
4.14
4.13
4.12
4.11
4.09
4.01
3.99
3.60
1.68
1.65
1.64
1.60
1.60
1.56
1.46
1.21
1.15
1.15
1.13
1.13
1.12
1.11
1.10
1.07
1.05
1.03
1.02
1.00
0.99
0.86
0.84
0.82
0.81
0.78
0.87
4.98
0.83
2.01
1.02
1.77
3.00
6.08
1.05
8.97
3.39
4.27
2.18
f1 (ppm)


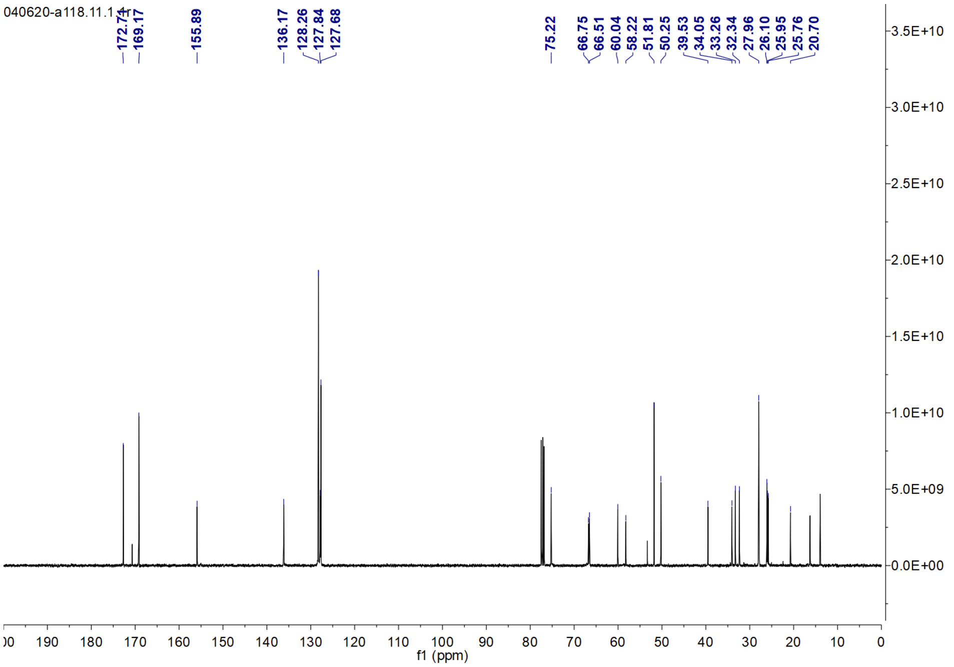

172.71
169.17
155.89
136.17
128.26
127.84
127.68
75.22
66.75
66.51
60.04
58.22
51.81
50.25
39.53
34.05
33.26
32.34
27.96
26.10
25.95
25.76
20.70
f1 (ppm)

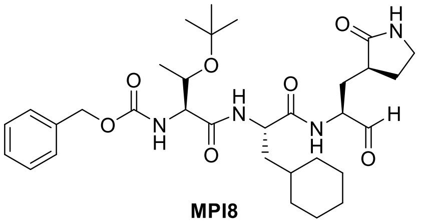

**MPI8**

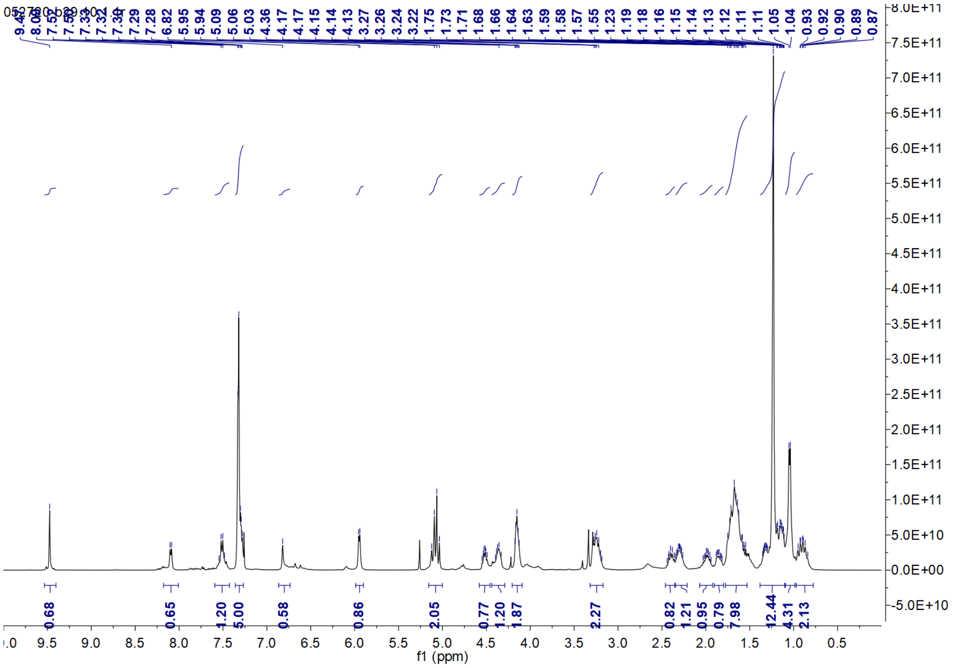
f1 (ppm)

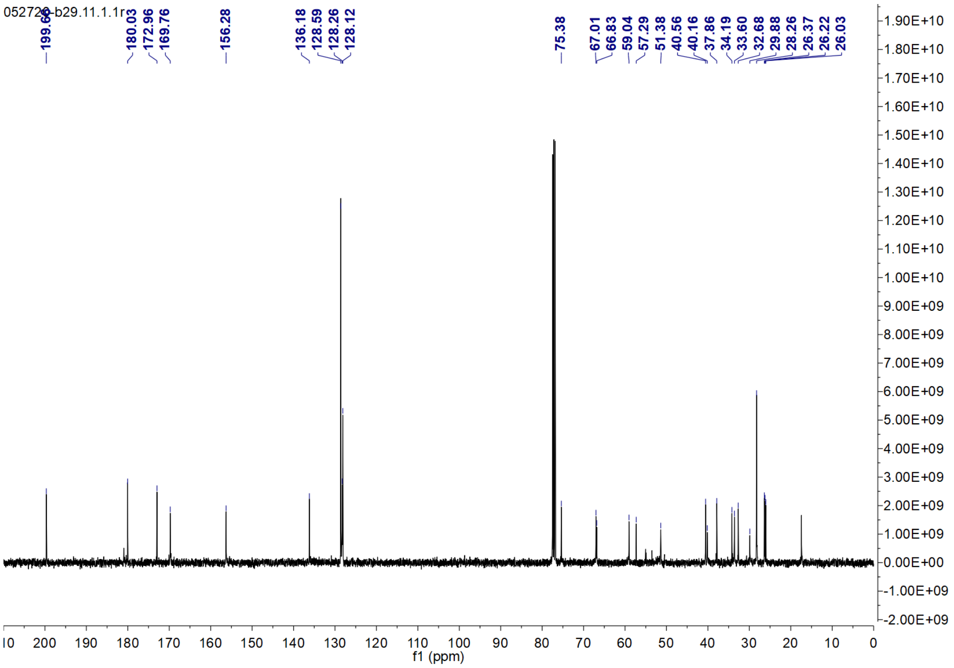
052720-b29.11.1.1r
199.60
180.03
172.96
169.76
156.28
136.18
128.59
128.26
128.12
75.38
67.01
66.83
59.04
57.29
51.38
40.56
40.16
37.86
34.19
33.60
32.68
29.88
28.26
26.37
26.22
26.03
f1 (ppm)

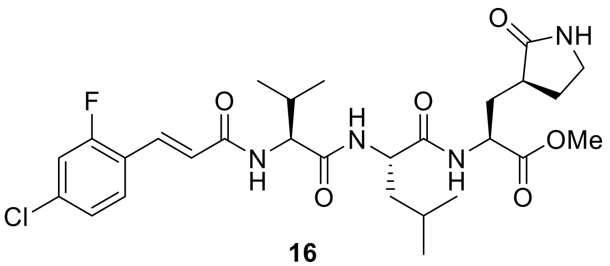

**16**

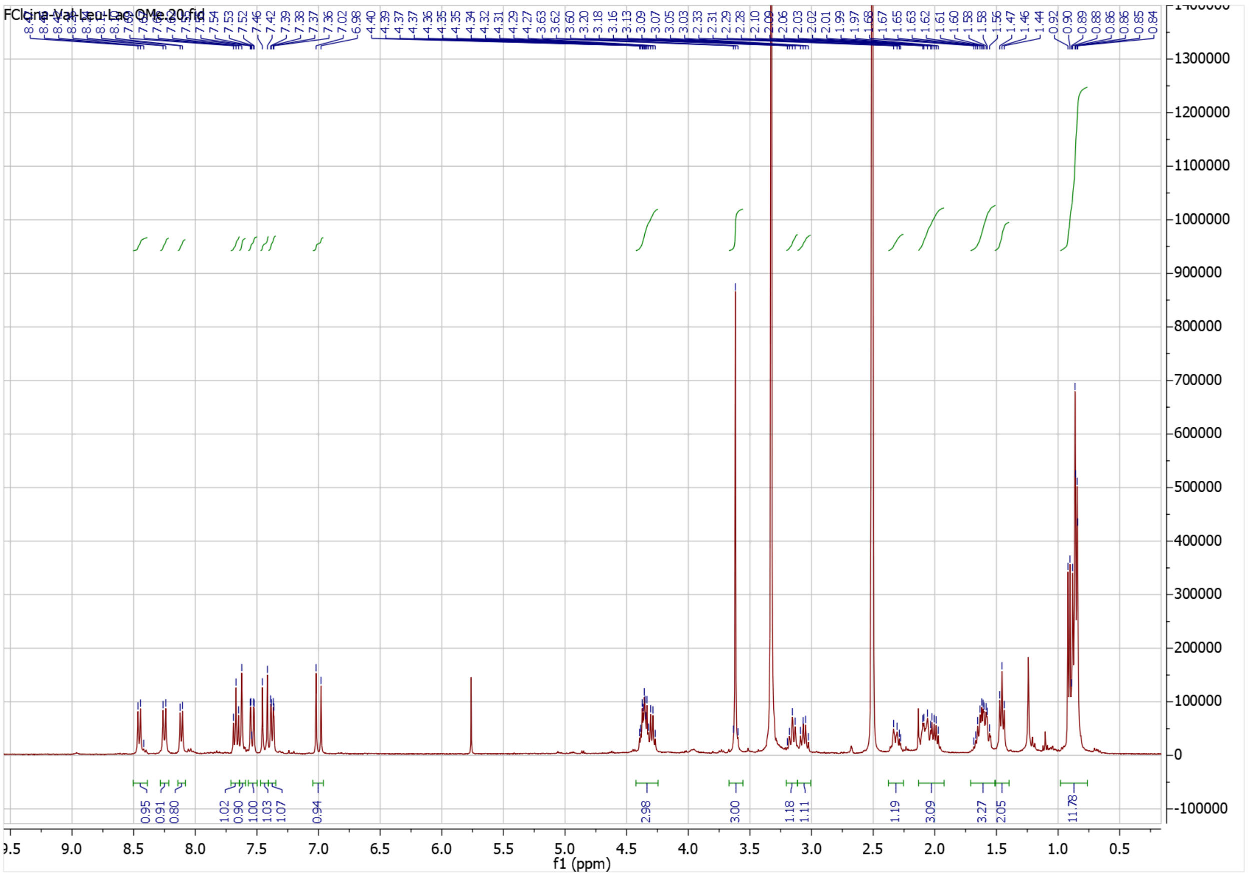

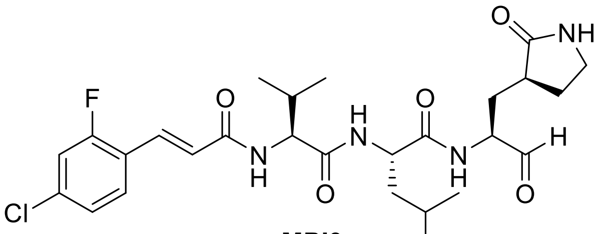

**MPI9**

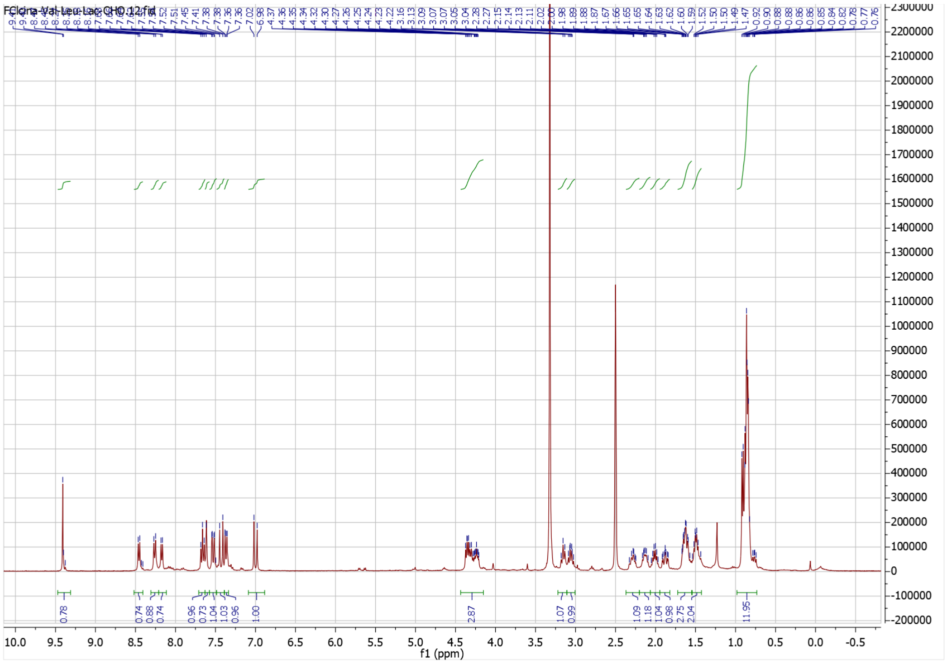
FClcina-Val-Leu-Lac-CHO.12.fid
f1 (ppm)

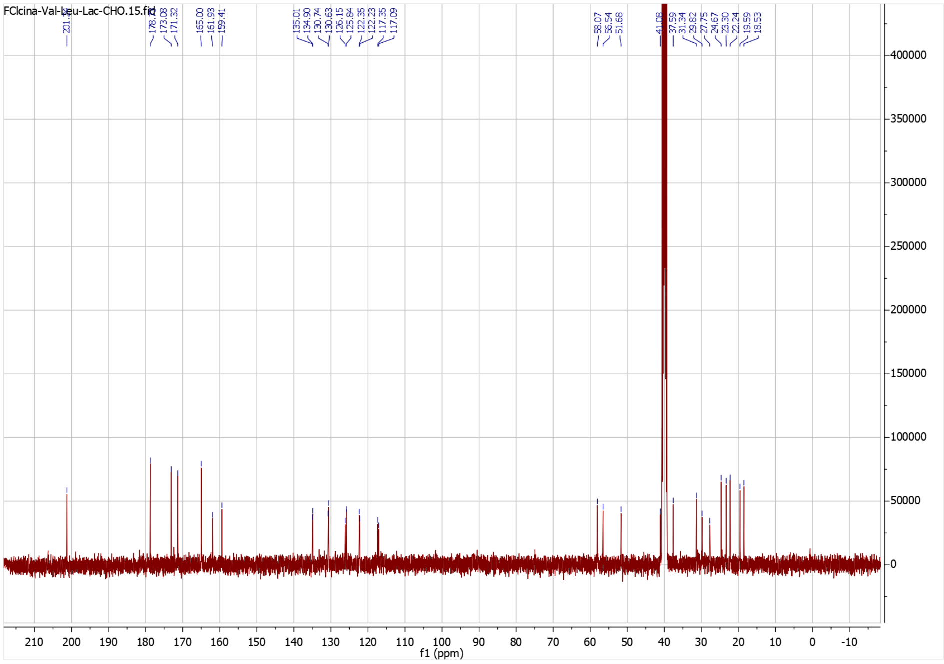
FClcina-Val-Leu-Lac-CHO.15.fid
f1 (ppm)
